# Atypical activation and molecular glue-like dimerization mechanism of an intrinsically-biased chemokine receptor

**DOI:** 10.1101/2025.10.30.685530

**Authors:** Nilanjana Banerjee, Manisankar Ganguly, Ashna Reyaz, Sudha Mishra, Annu Dalal, Xue Guo, Xingyu Qiu, Max Meyrath, Christie B. Palmer, Hongchen Guo, Siyuan Song, Shachie Sinha, Nabarun Roy, Debdatta Mukherjee, Divyanshu Tiwari, Manish K. Yadav, Andy Chevigne, Carol V. Robinson, Xin Chen, Ramanuj Banerjee, Arun K. Shukla

## Abstract

CXCR7, also known as atypical chemokine receptor 3 (ACKR3), is a naturally-biased, β-arrestin-coupled seven transmembrane receptor (7TMR) that lacks productive coupling with heterotrimeric G-proteins. Despite a critical involvement in cancer metastasis, cardiovascular pathophysiology, and inflammatory disorders, the molecular basis of non-canonical activation and functional divergence of CXCR7 remains elusive. Here, we present a complete landscape of CXCR7 activation using a series of cryo-EM structures, and discover an atypical activation mechanism that is distinct from prototypical GPCRs. CXCR7 is maintained in a basal conformation by a unique tripartite ionic-lock involving TM5-TM6, in contrast to a broadly conserved TM3-TM6 ionic-lock in GPCRs, which is disrupted upon receptor activation. Importantly, activation of CXCR7 results in a constricted pocket and distinct surface topology on the intracellular side compared to prototypical GPCRs. Serendipitously, we capture novel dimeric arrangements of CXCR7 with an inter-protomer stitching by a native phospholipid serving as a molecular glue, and identify previously unanticipated extrahelical allosteric sites on the receptor. Surprisingly, in an intermediate state structure of CXCR7, the second extracellular loop (ECL2) displays a self-blocking conformation, in stark contrast to ECL2-mediated self-activating mechanism reported recently for some orphan GPCRs. Finally, we unequivocally establish CXCR7 as an atypical opioid receptor via a large peptide library screening and structure elucidation in complex with distinct opioid peptides imparting full receptor activation. In summary, our study elucidates an atypical mechanism of CXCR7 activation, and establishes it as an alternative, non-canonical opioid receptor target with potential for novel pain therapeutics.

## Introduction

G protein-coupled receptors (GPCRs) constitute a large superfamily of integral membrane proteins, and they are critically involved in nearly all aspects of human physiology^1,2^. Aberrant GPCR signaling is linked to a broad spectrum of human disorders, and as a result, they are targeted by approximately one third of the currently prescribed medicines^3–5^. The broadly conserved paradigm of GPCR activation and signaling involves agonist-induced interaction of the receptors with heterotrimeric G-proteins, GRKs, and β-arrestins (βarrs). Structural biology efforts using crystallography and cryo-EM have illuminated the intricate details of receptor activation and transducer-coupling mechanisms over the past two decades^6,7^. Interestingly, the classical paradigm of GPCR activation has been expanded further with the discovery and integration of functional selectivity framework, also referred to as signaling-bias, and it has profoundly influenced the ongoing drug discovery efforts^8–11^. A series of recent studies combining biophysical and structural methodologies have started to provide generalizable insights into functional selectivity at the level of receptor and/or transducer conformations^12–14^. However, these studies are primarily limited to synthetic ligands and mutant receptors, and relatively little is known about the molecular basis and underlying principles of naturally-encoded signaling-bias at GPCRs.

Several members of the GPCR superfamily are incapable of activating heterotrimeric G-proteins, despite harboring a conserved seven transmembrane (7TM) architecture, but they still maintain robust GRK and βarr recruitment^15–18^. These arrestin-coupled receptors (ACRs) represent naturally-encoded GRK/βarr-biased 7TMRs, and offer a unique opportunity to decode the fundamental mechanisms directing distinct modalities of receptor activation and ensuing signaling-bias^19,20^. Most of these receptors belong to the subfamily of chemokine receptors, and also referred to as Atypical Chemokine Receptors (ACKRs)^16,17^, although additional examples from other receptor subfamilies have also emerged^19,21^. Previous studies on these receptors have focused predominantly on establishing their ligand-binding properties, transducer-coupling patterns, and non-canonical downstream signaling^19,22^, but the structural mechanism of their functional divergence remains mostly elusive. For example, these receptors appear to preferentially engage GRK5/6 for their phosphorylation and subsequent βarr recruitment, which aligns with the lack of G-protein activation and hence, inefficient translocation of GRK2/3 to the plasma membrane^19^.^23,24^ Unlike prototypical GPCRs, ACRs do not seem to elicit canonical downstream signaling cascades, potentially suggesting distinct modalities of transducer-coupling and receptor activation compared to GPCRs^19^. For instance, agonist-stimulation does not result in a measurable ERK1/2 phosphorylation downstream of ACRs, which has been used as a quintessential readout to define the spatiotemporal aspects of βarr signaling at prototypical GPCRs^19^. These unique functional manifestations distinguish these receptors from the functional selectivity scenario operational at prototypical GPCRs imparted by synthetic biased ligands, and establish them as unique system to explore the molecular mechanisms of naturally-encoded signaling-bias.

One of the naturally-biased 7TMRs, CXCR7, also referred to as ACKR3, is expressed widely in the endothelial and immune cells, and plays a central role in cellular migration, embryonic development, immune regulation, tissue homeostasis, and vascular functions^25^. Aberrant CXCR7 signaling is implicated in numerous disease contexts including tumour growth, metastasis and angiogenesis, vascular remodelling, multiple sclerosis, neuroinflammation, and autoimmune disorders^26^. CXCR7 recognizes two distinct C-X-C chemokines namely, CXCL11 and CXCL12, which also activate prototypical GPCRs i.e. CXCR3 and CXCR4, respectively, and couples efficiently to βarrs^27^. Similar to class A GPCRs, CXCR7 harbors the conserved sequence motifs such as the DRY in TM3 and NPXXY in TM7, which are implicated in G-protein-coupling and activation^28^. Therefore, its inability to functionally engage and activate G-proteins is even more perplexing, and potentially suggests an atypical activation mechanism driving the functional divergence. Two recent studies have provided the first glimpse of CXCL12 binding^29^ and βarr-coupling to CXCR7 using cryo-EM^30^. In these structural snapshots, CXCL12 appears to adopt a significantly different binding pose, compared to that by other chemokines on their cognate receptors, as visualized in the structures reported previously^29,30^. Still however, considering that the inactive structure of CXCR7, and that of any other ACRs, still remains unresolved, deciphering the activation mechanism, in particular, the distinguishing features compared to prototypical GPCRs, remains rather speculative. These knowledge gaps represent major lacunae in our current understanding of activation, divergent signaling, and non-canonical functions mediated by ACRs including CXCR7. Interestingly, CXCR7 is also proposed to recognize and scavenge opioid peptides, and potentially modulate the endogenous opioid system implicated in nociception^31–33^. However, the molecular basis of opioid-recognition by CXCR7 and ensuing receptor activation has not yet been studied at molecular level, which in turn, has limited the efforts to explore the potential of modulating CXCR7 for developing novel pain therapeutics.

Here, we elucidate an atypical activation mechanism of CXCR7 involving a significant rearrangement of unique ionic-locks distinct from prototypical GPCRs, and discover a pivotal switch that is also broadly conserved in chemoattractant receptors. We identify previously unanticipated extrahelical allosteric sites on the receptor for agonists and phospholipids, and surprisingly, also trap receptor dimers using a native lipid as molecular glue with an architecture that is reminiscent of distinct conformational states of membrane transporters. Moreover, using a combination of peptide library screening and cryo-EM-based structural elucidation, we unequivocally establish CXCR7 as an atypical opioid receptor, thereby, expanding the subfamily of opioid receptors beyond the conventional members. Our findings offer important insights into CXCR7 activation and signaling with broad and generalizable implications for the framework of naturally-encoded signaling-bias and development of novel therapeutics.

## Results

### A rich tapestry of CXCR7 ligand pharmacology

CXCR7 is one of the most well characterized chemokine receptors with respect to a wide spectrum of ligands having distinct chemical structures and molecular efficacies^34^. Therefore, we selected a combination of small molecules, peptide, nanobody, and natural chemokine ligands, each having a distinct pharmacology, to capture the complete landscape of CXCR7 activation. Of these, VUF16840 is a small molecule inverse agonist in terms of βarr-coupling, VUN701 is a nanobody-based antagonist, VUF11207 is a CXCR7-selective agonist, VUF10661 is a small molecule dual agonist at CXCR3 and CXCR7, TC14012 is a cyclic peptide agonist at CXCR7 but an inverse agonist at CXCR4, and CXCL11 is a natural chemokine agonist^34,35^ (**Figure 1a-b**). We first validated these ligands in extensive cellular assays in terms of βarr recruitment, which confirmed their pharmacology and efficacy profile, and also established TC14012 as CXCR7-selective cyclic-peptide agonist (**Figure 1c and Supplementary Figure S1A-D**). We also observed that CXCL11 is nearly as efficacious agonist as CXCL12 albeit with slightly lower potency, and VUN701 and VUF16840 are able to antagonize both, CXCL11 and CXCL12, although only VUF16840 exhibits an inverse agonism for βarr recruitment. Taken together, these ligands provide an opportunity to visualize distinct conformations of CXCR7 and fully understand receptor activation.

**Figure 1.**
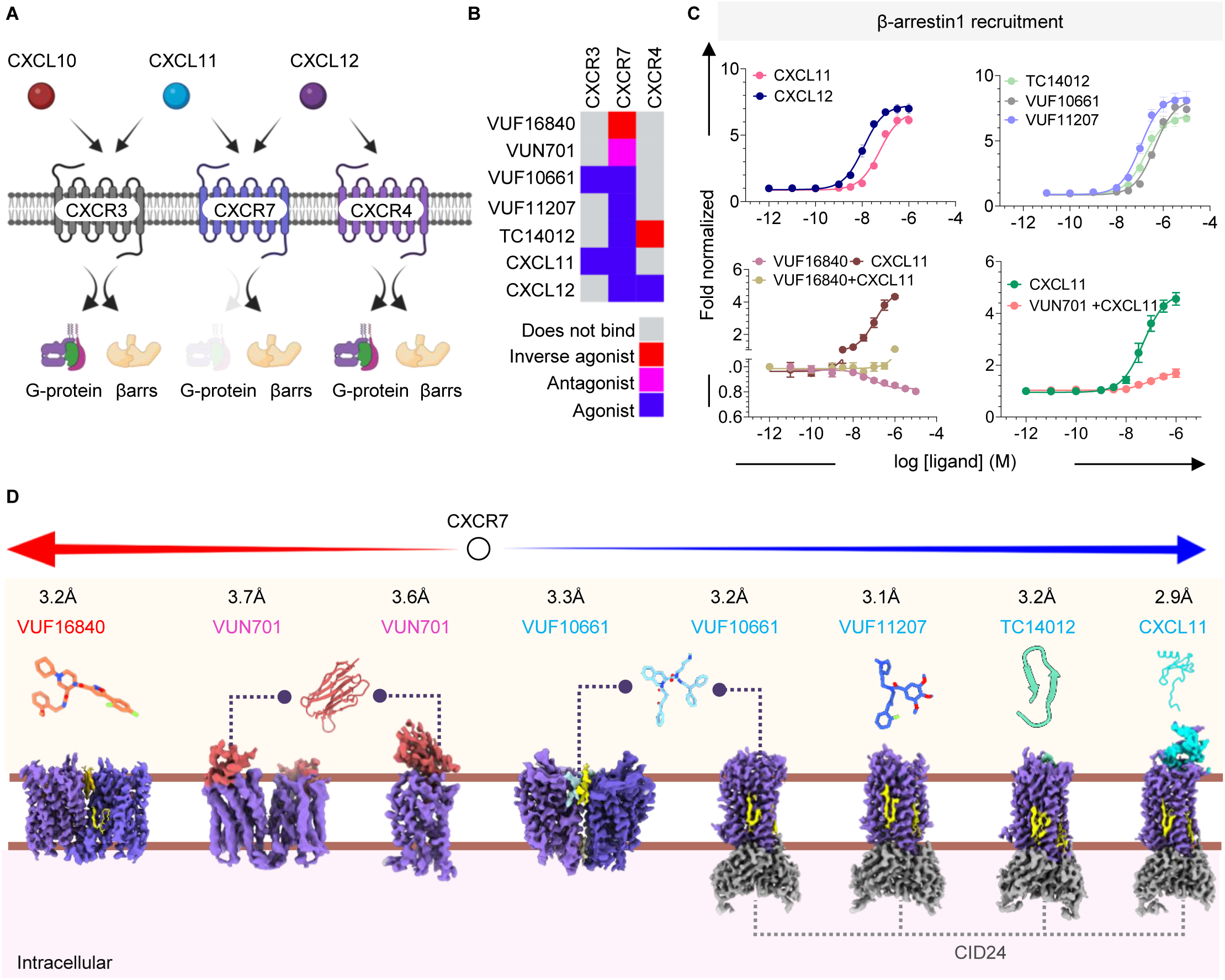
Ligand pharmacology and structure determination of CXCR7. **(A)** A schematic illustration showing chemokine-binding and transducer-coupling to CXCR3, CXCR4, and CXCR7. **(B)** Distinct pharmacology of selected CXCR7 ligands presented based on published literature. **(C)** Ligand-induced βarr1 recruitment at CXCR7 measured using NanoBiT-based assay in HEK-293T cells (mean±sem; n=3; fold normalized with basal signal). **(D)** Structures of CXCR7 in complex with indicated ligands determined using cryo-EM, stabilized using CID24 (an antibody fragment) as indicated.

### Structure determination of CXCR7 using cryo-EM

In order to determine the structures of agonist-bound CXCR7, we leveraged a previously described antibody fragment, CID24, which binds on the intracellular side of the receptor, and presumably recognizes an active conformation^29^. We observed that CID24 is not able to recognize antagonist or inverse-agonist-bound CXCR7 (**Supplementary Figure S2A**), and therefore, we anticipated that capturing the inverse agonist/antagonist-bound CXCR7 will be technically challenging due to the small size of the receptor. Serendipitously however, we observed during cryo-EM data processing that VUF16840-CXCR7 predominantly exhibited a dimeric population yielding a structure at 3.2Å resolution (**Figure 1D and Supplementary Figure S3**). For the VUN701-CXCR7 complex, both, monomeric and dimeric populations were apparent during cryo-EM data processing, and we determined their structures at 3.6Å and 3.7Å, respectively (**Figure 1D and Supplementary Figure S3**). On the other hand, the agonist-bound CXCR7 complexes, stabilized by CID24, yielded structures ranging from 2.9Å to 3.2Å, where the receptor was present mostly as a monomer except VUF10661 (**Figure 1D and Supplementary Figure S5**). We observed both, monomer and dimer population for VUF10661-bound CXCR7, and determined their structures at 3.2Å and 3.3Å, respectively (**Figure 1D and Supplementary Figure S4 and S5)**. Interestingly, the monomer structure contains CID24 on the intracellular side of the receptor, but the dimer structure did not show discernible evidence of CID24 binding (**Figure 1D**). All the structures showed clear densities for the bound ligands, majority of the receptor segment, and also distinct lipid molecules bound to the receptor, leading to the discovery of several fundamental insights as described subsequently. The technical details of cryo-EM data collection and processing are outlined in **Table S1**, and the cryo-EM densities for the TMs and ligands are presented in the **Supplementary Figure S6-S8**. Notably, we observed a significantly larger portion of the N-terminus of CXCR7 in the CXCL11-bound structure due to an engagement with the ligand, which was not resolved in other structures as they do not extensively engage the N-terminus of the receptor (**Supplementary Figure S9A-B)**. In addition, we also observed a clear disulfide bond between Cys34^N-term^ and Cys287^7.25^ in the active structures of CXCR7, while it was absent in the inactive state structures due to a linear shift of the receptor N-terminus (**Supplementary Figure S9C-D**). Interestingly, this disulfide bond was not resolved in the previously determined CXCR7 structure in complex with CXCL12^29^, although it appears to be a critical determinant of receptor stabilization in an active state based on our structural analysis and extensive site-directed mutagenesis studies^36^.

### Overall ligand binding to CXCR7

Most ligands expectedly engage primarily in the orthosteric binding pocket of the receptor with the exception of VUF10661, which shows a dual binding mode. For example, VUF16840 occupies the base of the orthosteric pocket with a distance of ∼4Å from the conserved Trp265^6.48^ and encompasses a buried surface area of ∼1600Å^2^ (**Figure 2a**). The orthosteric binding pocket is characterized by the presence of several aromatic amino acid residues such as Tyr51^1.39^, Trp100^2.60^, His121^3.29^, Phe124^3.32^, Phe129^3.37^, Trp265^6.48^, and Tyr268^6.51^, and hydrophobic residues such as Leu128^3.36^, Ile132^3.40^, Leu305^7.43^, and Leu183^4.64^. Together, these residues create an optimal environment for the binding of VUF16840 (**Supplementary Figure S10A)**. Small molecule agonists VUF10661 and VUF11207 also occupy a similar space in the orthosteric binding pocket although they penetrate a little deeper compared to VUF16840 in the pocket (**Figure 2a and Supplementary Figure S10B-C**). As mentioned earlier, VUF10661 is a dual agonist for CXCR3 and CXCR7, and it occupies a similar position in the orthosteric of both, CXCR3 and CXCR7, and engages mostly a similar set of residues (**Supplementary Figure S10D-E)**. This is in excellent agreement with our site-directed mutagenesis data showing a comparable impact of analogous residues in both receptors on VUF10661 binding^37^. On the other hand, CXCL11 exhibits a two-site binding mechanism on CXCR7, wherein the core domain engages the N-terminus and ECL2 of the receptor, while the N-terminus penetrates deep into the orthosteric binding pocket (**Figure 2A and Supplementary Figure S11A**). Overall, CXCL11 binding encompasses more than 3,000Å² buried surface area, and is stabilized by an array of hydrogen bonds, hydrophobic interactions, salt bridges, and polar contacts (**Supplementary Figure S11B**). A partially negatively charged segment in the N-terminus of CXCR7 spanning residues Ser^23^-Met^37^ docks into a positively charged groove formed by helix-1 and β3 strand of the CXCL11 core domain as a part of the first binding site (**Supplementary Figure S11B**). On the other hand, the proximal N-terminus of CXCL11 makes a series of contacts with distinct residues in TM1-7 in the orthosteric binding pocket as the second binding site on the receptor.

**Figure 2.**
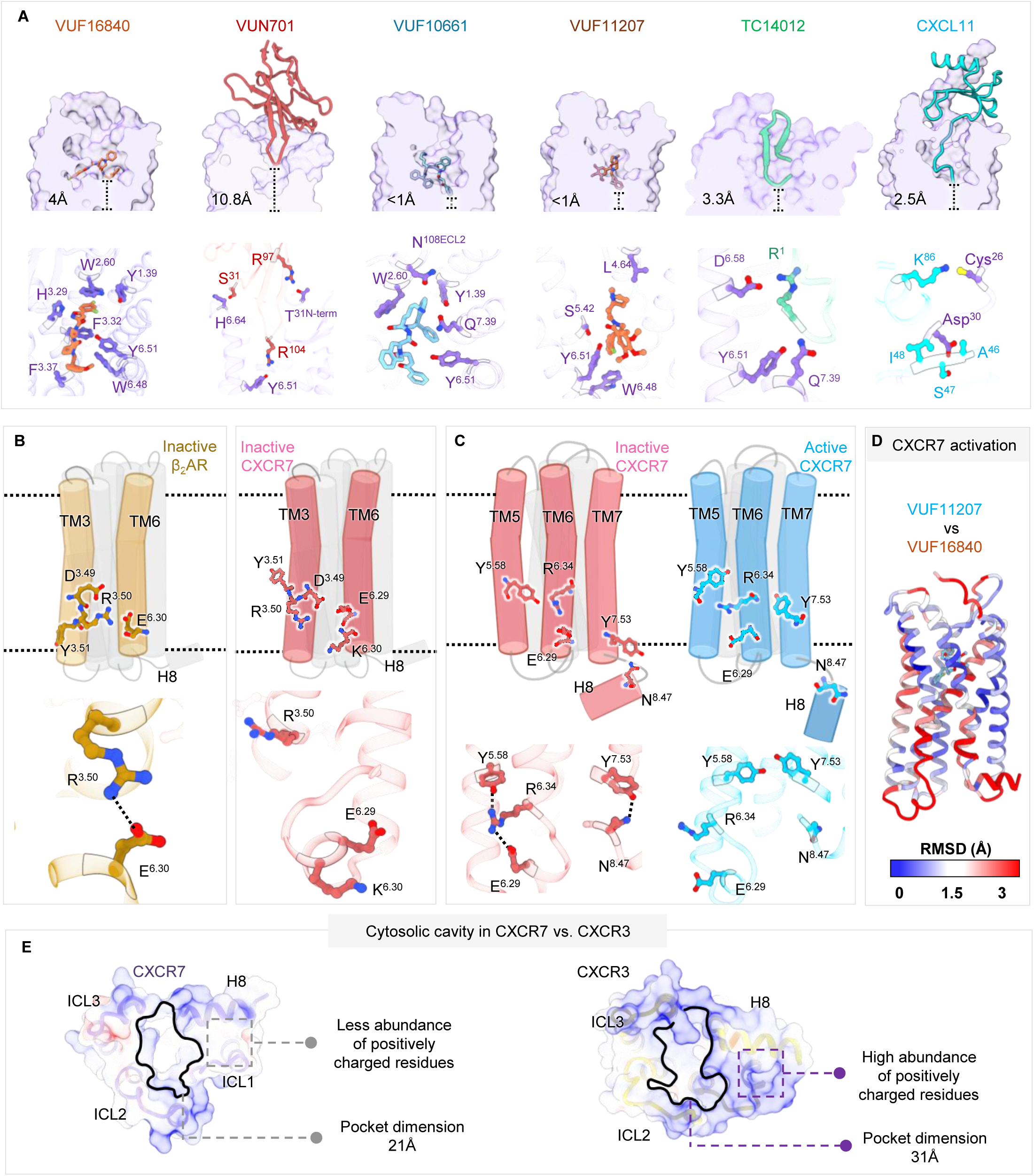
Ligand-recognition and atypical activation of CXCR7. **(A)** Overall binding pose (upper panel) and key interactions for the indicated ligands in the orthosteric binding pocket. The distance between the deepest point of the ligands from the conserved W6.48 is indicated in the upper panel. **(B)** A schematic representation and ribbon diagram showing the presence and absence of a broadly conserved TM3-TM6 ionic-lock in β2AR and CXCR7, respectively. The orientations of R3.50 (TM3) and E6.30/6.29 (TM6) are presented based on the structural snapshots (PDB IDs: 2RH1 for β2AR; 9WLD for CXCL11-CXCR7). **(C)** A schematic representation and ribbon diagram showing two distinct ionic locks i.e. TM5-TM6 and TM7-H8 in the inactive state of CXCR7 limiting the TM movement (left panel; red), and their disruption upon receptor activation (right panel; blue). **(D)** Activation dependent conformational changes in CXCR7 depicted in terms of RMSD between inverse-agonist vs. agonist-bound structures (VUF16840 vs. VUF11207-bound structures) mapped onto a ribbon representation. **(E)** Distinct cytoplasmic surface of the active CXCR7 compared to CXCR3 indicating a relatively constricted dimension of the cytoplasmic pocket and lesser abundance of positively charged residues on a patch involved in αN-helix engagement (PDB IDs: 9WLD for CXCL11-CXCR; 8HNK for CXCR3).

Structural alignment of CXCL11-bound CXCR7 complex with the previously reported CXCL12-CXCR7 structure indicates that despite adopting broadly similar binding orientations, the chemokine core domains are laterally displaced relative to each another (**Supplementary Figure S11C**). Structural superimposition of currently available C-X-C type chemokine receptors in complex with different C-X-C chemokines indicates that the chemokines adopt an overall similar binding pose with minor localized variations. In contrast, in the CXCL11–CXCR7 complex, the chemokine core domain is oriented at ∼90°, while the N-terminus adopts an open, slender conformation that penetrates more deeply into the orthosteric pocket of CXCR7 (**Supplementary Figure S11D).** This is also reflected in a comparison of CXCL11-CXCR7 with previously determined structure of CXCL11-CXCR3^38^, which reveals a large shift in the positioning of the same ligand i.e. CXCL11 on the two receptors (**Supplementary Figure S11E**). Specifically, the core domain of CXCL11 on CXCR7 adopts a torsional conformation rotated by approximately 90°, which facilitates more extensive interactions with the receptor N-terminus, compared to CXCR3 (**Supplementary Figure S11E**). Moreover, while N-terminus of CXCL11 aligns closely within the orthosteric pocket of both receptors, the N-loop residues in CXCL11 diverge by nearly 180° between the two receptors (**Supplementary Figure S11E**). These pronounced differences in the binding of CXCL11 to CXCR3 vs. CXCR7 likely contribute to their functional divergence in terms of transducer-coupling and functional outcomes.

In case of VUN701, the extended CDR3 penetrates into the orthosteric pocket of the receptor, and surprisingly, the N-terminus of CXCR7 wraps around the CDR3 to stabilize the binding further (**Supplementary Figure S11F**). This is in stark contrast with CXCL11-CXCR7 structure, wherein the N-terminus of the receptor adopts a vertical orientation to interact with, and stabilize, the core domain of CXCL11 (**Supplementary Figure S11F**). In addition, CXCL11 penetrates approximately 8Å deeper in the orthosteric binding pocket compared to the CDR3 of VUN701 (**Figure 2A**). This observation underscores a relatively superficial binding of VUN701 and likely suggests that it antagonizes CXCR7 by sterically obstructing the entry and docking of agonists including chemokines. Interestingly, TC14012 adopts a slender conformation despite being a cyclic peptide, and inserts deep into the orthosteric pocket of the receptor, almost to a comparable depth with CXCL11, and makes similar interactions as that of the N-terminus of CXCL11 (**Figure 2A and Supplementary Figure S11G**).

### Distinct ionic-locks stabilize inactive CXCR7 conformation

Class A GPCRs typically harbor a TM3-TM6 ionic-lock where Arg^3.50^ of the conserved DRY motif in TM3 forms an interhelical salt-bridge with a conserved Asp/Glu^6.30^ in TM6, and helps maintain the receptors in their basal conformation^39,40^. This salt-bridge is disrupted upon receptor activation leading to large TM movements, especially TM6, leading to subsequent coupling and activation of heterotrimeric G-proteins^41,42^. Interestingly, in CXCR7, the position 6.30 in TM6 is occupied by a lysine residue, which makes the formation of a typical salt-bridge with Arg^3.50^ unfeasible, and VUF16840-bound structure indeed corroborates the same (**Figure 2B**). A previous study has proposed that Glu^6.29^ in CXCR7 may serve as a surrogate for Asp/Glu^6.30^ present in other class A GPCRs, and thereby, still allow the formation of a slightly altered TM3-TM6 ionic-lock^43^. In contrast, VUF16840-bound CXCR7 structure elucidates that not only Arg^3.50^ is oriented towards the receptor core and away from Glu^6.29^ but the side-chain of Glu^6.29^ is also oriented in an opposite direction, which in turn makes any contact with Arg^3.50^ implausible (**Figure 2B**). Instead, we observed that Glu^6.29^ forms a distinct tripartite lock by making a salt-bridge with Arg^6.34^, which in turn interacts with Tyr^5.58^ via an ionic-interaction (**Figure 2C**). In addition, we also observed that Tyr^7.53^ in TM7, which is also a part of the conserved NPxxY motif, interacts with Asn^8.47^ in helix 8, and this engagement is likely to restrict the mobility of TM7 and helix8 in the basal conformation of CXCR7 (**Figure 2C**). Interestingly, these two distinct ionic-locks are also conserved in both, monomeric and dimeric structures of VUN701-bound CXCR7 (**Supplementary Figure S12A-B**), and therefore, establish them as a bonafide feature that maintains CXCR7 in its basal conformation.

### Atypical activation of CXCR7

Structural comparison of the inactive and active CXCR7 structures reveals large conformational changes in the receptor, especially, an outward movement of cytoplasmic side of TM6 (∼7Å) accompanied by a significant rotation of helix 8 (∼50°) towards the cytoplasmic core of the receptor, and a relatively smaller movement of TM7 (∼4Å) (**Supplementary Figure S12C**). Unlike other class A GPCRs, the movement of TM5 upon CXCR7 activation appears to be restricted (<1Å). The movement of cytoplasmic end of TM6 upon receptor activation appears to be essential for CID24 binding, and the position of TM6 in the inactive state is likely to clash with CID24 engagement. Interestingly, upon receptor activation, Tyr^5.58^ undergoes an upward rotation (∼75°) towards the receptor core, while Arg^6.34^ and Glu^6.29^ shift by approximately 5-6Å away from the core, and these structural changes result in complete disruption of the tripartite ionic-lock (**Figure 2B**). Concomitantly, Tyr^7.53^ in TM7 and Asn^8.47^ in helix8 also undergo a large upward and downward rotation, respectively, which disrupts their interaction by positioning their side-chains in the opposite directions (**Figures 2B**). Collectively, these structural changes in CXCR7 disrupt the non-canonical ionic-locks and thereby, relieve the restraints on TM6, 7 and helix8, and consequently, result in receptor activation. Importantly, these activation-dependent structural changes in CXCR7 are consistent across all the active state structures stabilized by different agonists, and therefore, unequivocally establish the mechanistic basis of receptor activation. Interestingly, the cytoplasmic surface of CXCR7 upon activation also reveals two distinctive features compared to prototypical GPCRs. First, the overall dimension of the cytoplasmic cavity generated upon receptor activation is significantly smaller than that observed for prototypical GPCRs including other chemokine receptors such as CXCR3, which may not be sufficient to support a stable interaction with the α5-helix of the G-proteins (**Figures 2E and Supplementary Figure S12D**). This constricted intracellular pocket likely results from a relatively smaller outward movement of TM5 and TM6 in CXCR7, compared to prototypical GPCRs. Second, the cytoplasmic surface of CXCR7 in active conformation lacks a positively charged patch of residues that is apparent in other chemokine receptors and accommodates the αN helix of Gβ subunit (**Figure 2E and Supplementary Figure S12E**). Taken together, these two features provide a plausible explanation for the lack of G-protein-coupling and activation by CXCR7.

### Distinct dimerization of CXCR7

Several class A GPCRs have a propensity to homo- and hetero-dimerize as assessed using cellular assays, and this impacts their functional outcomes in terms of trafficking and downstream signaling^44–46^. CXCR7 is reported to form homodimers as well as heterodimers with other chemokine receptors such as CXCR4 with distinct functional consequences^47,48^. As mentioned earlier, we have captured CXCR7 dimeric arrangement in three different structures in complex with three distinct ligands **(Supplementary Figure S13A-B)**. The dimeric structure in complex with VUF16840 encompasses a dimension of ∼80Å along the longer axis and ∼65Å along the broader axis, and harbors a buried surface area and interface area of ∼2200Å^2^ and ∼1100Å^2^, respectively. The dimer interface is mediated primarily by TM1 and helix8 from each protomer, wherein TM1 is positioned diagonally at an angle of ∼90° while helix8 is aligned almost parallel to each other (**Figure 3A-B**). Three hydrogen bonds and a large number of non-bonded contacts stabilize the dimer interface suggesting a robust interaction. These include, for example, hydrophobic interactions of Met59^1.47^ and Trp67^1.55^ from one protomer with Leu91^2.51^ and Ile60^1.48^/Val64^1.52^ in the other protomer (**Figure 3C**) and hydrogen binding between Ser63^1.51^ from each protomer across the interface (**Figure 3C**). In addition, the helix8 interface involves a hydrogen bond between Arg323^8.51^ of one protomer with Met327^8.55^ of the other as well as a cation-π interaction between Arg320^8.48^ and Phe330^8.58^ of the two protomers, respectively (**Figure 3C**). Interestingly, the dimeric assembly of VUN701-bound CXCR7 closely resembles that observed in VUF16840-CXCR7 complex, with the dimerization interface mediated primarily by TM1 and helix8, and a comparable cross-angle between the protomers (**Figure 3A-B**).

**Figure 3.**
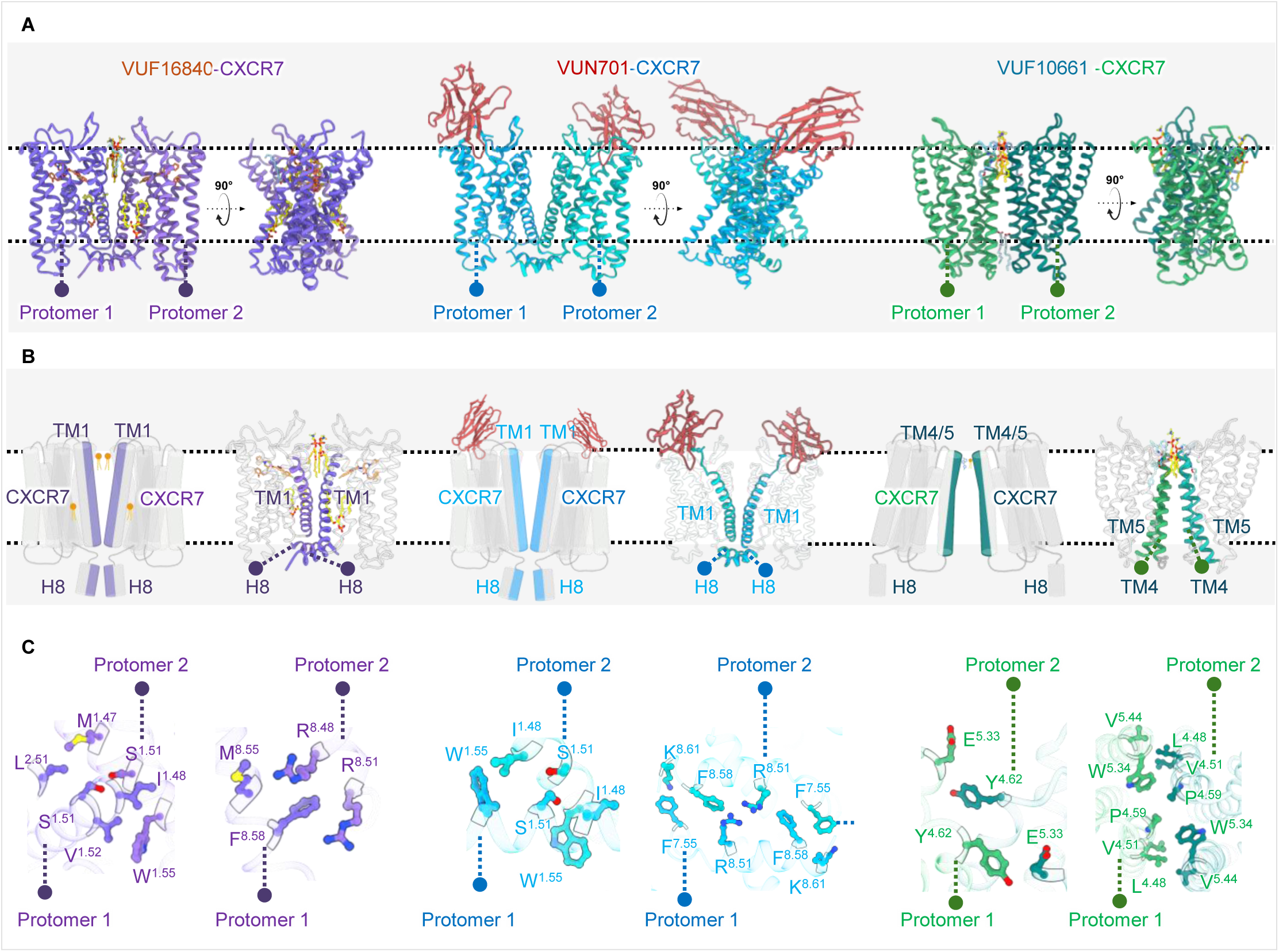
Dimerization of CXCR7 via distinct interfaces. **(A)** Structural representation of CXCR7 dimers as captured by cryo-EM in complex with VUF16840, VUN701, and VUF10661. Ribbon diagram at two different orientations are presented for visualization (PDB IDs: 9WLG for VUF16840-CXCR7; 9WLO for VUN701-CXCR7; 9WLM for VUF10661-CXCR7). **(B)** Schematic illustration and ribbon diagram of CXCR7 dimers showing the dimerization interface. Although the dimerization interface between VUF16840-CXCR7 and VUN701-CXCR7 are similar (i.e., TM1-TM1 and H8-H8), the two protomers are oriented differently with respect to each other. **(C)** The key interacting residues in CXCR7 dimers, contributed by each protomer, are presented based on the corresponding structural snapshots.

On the other hand, the dimeric assembly of VUF10661-bound CXCR7 is distinct from that observed in the VUF16840/VUN701-bound CXCR7 dimers. Here, the two protomers align parallel to each other with TM4 and TM5 constituting the primary dimer interface with minor contribution from the cytoplasmic region of TM3 (**Figure 3A-B**). The absence of CID24 in this structure suggested an inactive-like receptor conformation, and further structural analysis as discussed subsequently indeed corroborates this. The overall buried surface area at the dimer interface is approximately 1000Å², and Tyr181^4.62^ of each protomer forms a hydrogen bond with Glu207^5.33^ of the opposing protomer, and vice-versa. A prominent hydrophobic cluster at the interface further contributes to the stability of the oligomeric complex, which includes Leu167^4.48^, Val170^4.51^, Pro178^4.59^, Trp208^5.34^, and Val218^5.44^ from both protomers (**Figure 3C**). The overall conformation of each protomers in the dimeric structure are nearly identical with a rmsd of <1Å. Interestingly however, while the receptor conformations between VUN701-CXCR7 monomer vs. dimer are mostly similar except a slight shift in CDR3 positioning (**Supplementary Figure S13C)**, there are stark differences between the receptor conformation between the monomer vs. dimer of VUF10661-bound CXCR7 as discussed subsequently.

### Interaction of native lipids to CXCR7 and a molecular glue-like mechanism

Several cryo-EM structures of GPCRs determined previously have shown a presence of bound cholesterol molecules, and for some, specific functional impact has also been demonstrated using cellular and biophysical assays^49,50^. Surprisingly, the cryo-EM structures of CXCR7 determined here exhibit clear densities consistent with several lipid molecules, not only at the dimeric interface but also bound to the receptor monomers (**Figure 4A-D**). As there is no prior information on the interaction of non-cholesterol lipids to CXCR7, their modelling and placement are guided by the observed densities, assignment in other membrane protein structures in the Protein Data Bank, and experimentally confirmed abundance in cultured insect cells. In the active state CXCR7 structures, there are two distinct lipid densities, which correspond to dipalmitoyl phosphatidic acids (LPP) molecules. The first LPP molecule occupies a hydrophobic sub-pocket formed with contributions of residues from TM2, TM3, and TM4 (**Figure 4A-B**). The phosphate head group of this LPP molecule forms hydrogen bonds with Lys73^ICL1^ and Arg162^4.43^, while the aliphatic chains engage with several residues from TM2, TM3, and TM4 via hydrophobic interactions (**Figure 4A and 4D**). The second LPP molecule occupies a similar hydrophobic sub-pocket composed of residues from TM3, TM4, and TM5, and it is stabilized by a series of non-bonded contacts with the receptor. Importantly, in the previously published CXCL12-CXCR7 structure^29^, two cholesterol molecules were modelled in the densities corresponding to the lipid moieties (**Supplementary Figure S14A-B**). However, the structures presented here at improved resolution clearly suggest that these densities correspond to phospholipids with two aliphatic chains. In order to corroborate these findings, we carried out mass spectrometry (MS)-based lipidomics analysis to identify the specific lipid moieties that are bound to CXCR7. Interestingly, we identified several phospholipids that are abundantly present in the *Sf*9 membrane, and some of these are enriched in purified receptor sample (**Supplementary Figure S14C**). Importantly, we noticed a robust enrichment of phosphatidic acid (16:0/16:1) in the purified receptor sample, despite relatively lower abundance in the membranes. This is in an excellent agreement with the placement of LPP in the cryo-EM density, which is repeatedly observed in every CXCR7 structure determined here including different batches of purified receptor.

**Figure 4.**
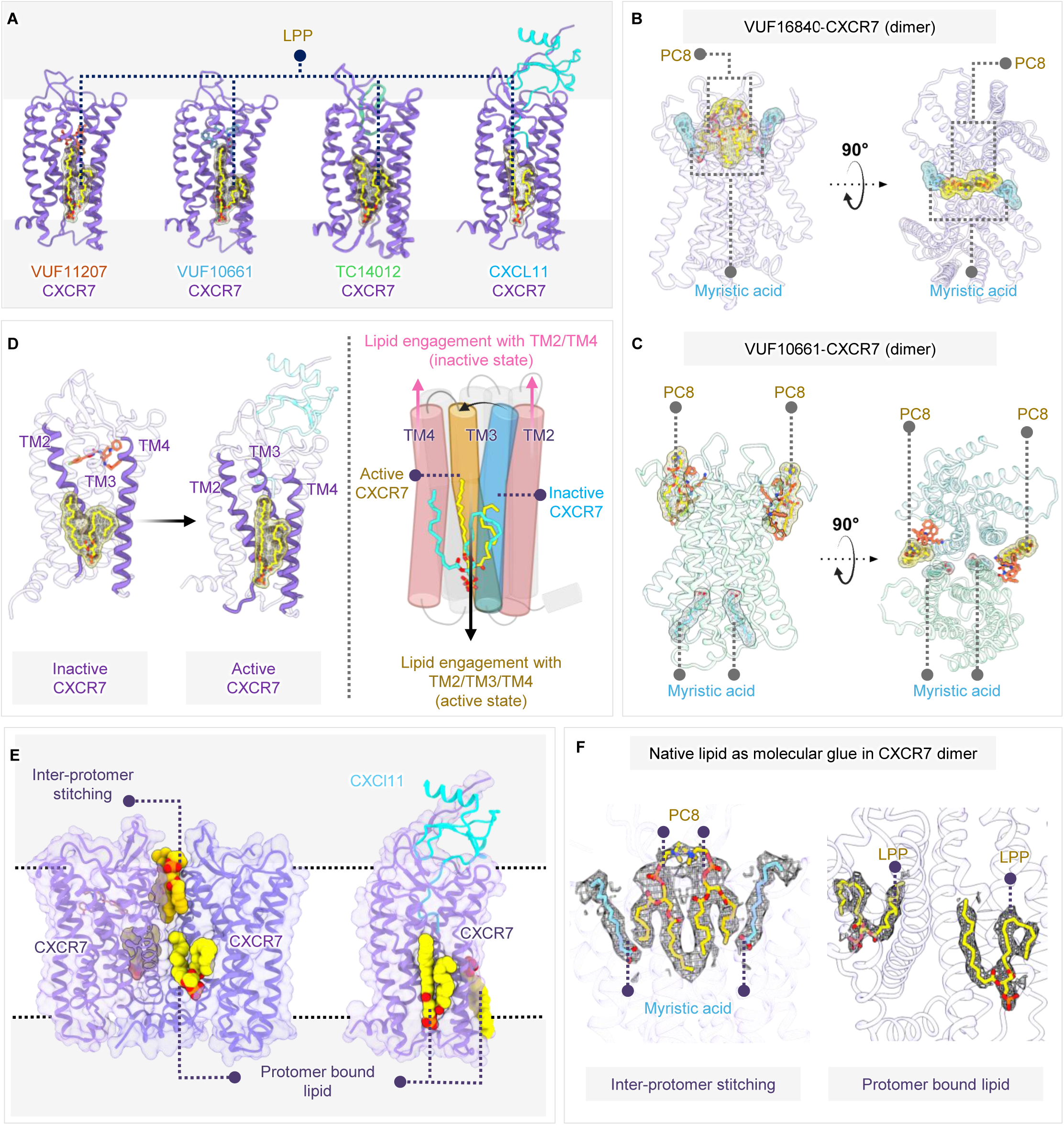
Interaction of native lipids with CXCR7. **(A)** Ribbon diagram of CXCR7 structures depicting the binding of native lipids to the receptor monomer in the agonist-bound state. The lipid-binding interface in terms of the TM helices are highlighted. **(B-C)** Ribbon diagram of CXCR7 dimeric structures depicting the binding of native lipids to each protomer and at the dimer interface. The lipid-binding interface in terms of the TM helices contributed by each protomer is highlighted. **(D)** Ribbon diagram and schematic illustration depicting the rearrangement of lipid-binding interface and the aliphatic chains due to conformational change in CXCR7 upon activation. **(E-F)** Lipid-mediated inter-protomer stitching as observed in VUF16840-CXCR7 and VUF10661-CXCR7 structures. The cross-interaction of the protomers mediated by the lipid moieties is indicated.

In addition, we also observed two putative phosphatidylcholine (PC) molecules at the dimer interface in the inactive state structures, each coming from the individual protomers (**Figure 4B and Supplementary Figure S15A-B**). The lipid-binding microenvironment at the dimeric interface is predominantly hydrophobic, formed by several residues contributed primarily by TM2 and TM3 (**Supplementary Figure S15C)** These two phosphatidylcholine molecules located at this interface form extensive hydrophobic interactions with residues from both protomers, thereby, effectively stitching the two protomers together by serving as a molecular glue and stabilizing the dimeric arrangement. In addition to these lipids, we also observed additional density corresponding to a lipid moiety in some of the structures such as VUF11207-bound CXCR7 and VUF16840/10661-bound dimers, which we infer to represent myristic acid (**Figure 4F**). For example, in the VUF10661-CXCR7 structure, myristic acid molecule is present at the dimeric interface, and it is positioned between the cytoplasmic segment of TM3 of one protomer and the ICL2 of the adjacent protomer. It is further stabilized via non-bonded interactions involving Tyr143^3.51^ in the DRY motif of one protomer and Arg156^ICL2^ of the other protomer. We did not model lipid density in the VUN701-bound CXCR7 structures due to relatively lower resolution, although, at lower contour levels, the corresponding density appears to emerge. We also note that a precise identification of these lipids, and their functional roles receptor activation and signaling remains to be determined in future studies.

### A novel allosteric site and an intermediate CXCR7 conformation

As mentioned earlier, the monomeric structure of VUF10661-bound CXCR7 exhibits an active conformation of the receptor as reflected by the large conformational changes compared to the inactive state, and VUF10661 occupies the orthosteric binding pocket (**Supplementary Figure S10B and S10D)**. Interestingly however, in the dimeric structure, VUF10661 binds to an extra-helical allosteric site located at the dimer interface formed by the extracellular segments of TM3, TM4, and part of ECL2 (**Figure 5A-B**). Specifically, each copy of the ligand interacts with Glu114^3.22^, Lys118^3.26^, Val119^3.27^, and Thr180^4.61^ of one protomer and Glu207^5.33^ of the other (**Figure 5C**). Additionally, we also observed that the phosphatidylcholine molecule is positioned adjacent to VUF10661, and engages with Lys206^5.32^, Ile210^5.36^ and Leu214^5.40^, thereby, providing further stabilizing the binding of the ligand to the allosteric site and serving as a molecular glue-like (**Figure 5C**). Interestingly, both CXCR7 protomers in this structure are present in an inactive-like conformation, and they align well with the inverse-agonist-bound conformation with an overall rmsd of <1Å. Still however, some notable conformational differences are observed in the extracellular loop regions. For example, ECL1 exhibits an outward linear displacement of approximately 11Å (measured at the Cα atom of Gln^109^) while the β-hairpin of ECL2 shifts laterally away from the orthosteric pocket by approximately 6Å. Surprisingly, a part of ECL2, spanning the residues from Tyr^195^ to His^203^, folds back and penetrate the orthosteric pocket (**Figure 5D**). Interestingly, ECL2 fold-back structure and positioning in the orthosteric binding pocket is reported as a self-activating mechanism for a couple of orphan receptors such as GPR161^51^ and GPR52^52^. Contrary to this, ECL2 in VUF10661-bound CXCR7 appears to block the orthosteric binding pocket of the receptor, and thereby, pushing the ligand to an allosteric site, stabilizing an inactive-like conformation **(Supplementary Figure S16A)**. To our knowledge, this is the first example of an autoinhibitory mechanism mediated by ECL2 in any GPCR structure reported till date, and it underscores the versatility and ligand-specific conformational adaptation encoded by the 7TM scaffold. Moreover, the newly identified ionic-locks in CXCR7 are present in an intermediate state in this structure. Specifically, the Arg251^6.34^-Glu246^6.29^ and Tyr315^7.53^-Asn319^8.47^ interactions are maintained, while Arg251^6.34^ is positioned more than 4Å away from Tyr232^5.58^, thereby, weakening the tripartite lock compared to the fully inactive structure (**Supplementary Figure S16B)**. We acknowledge that future studies are required to further explore the functional consequences of this allosteric site in CXCR7 and the role of ECL2 in receptor activation.

**Figure 5.**
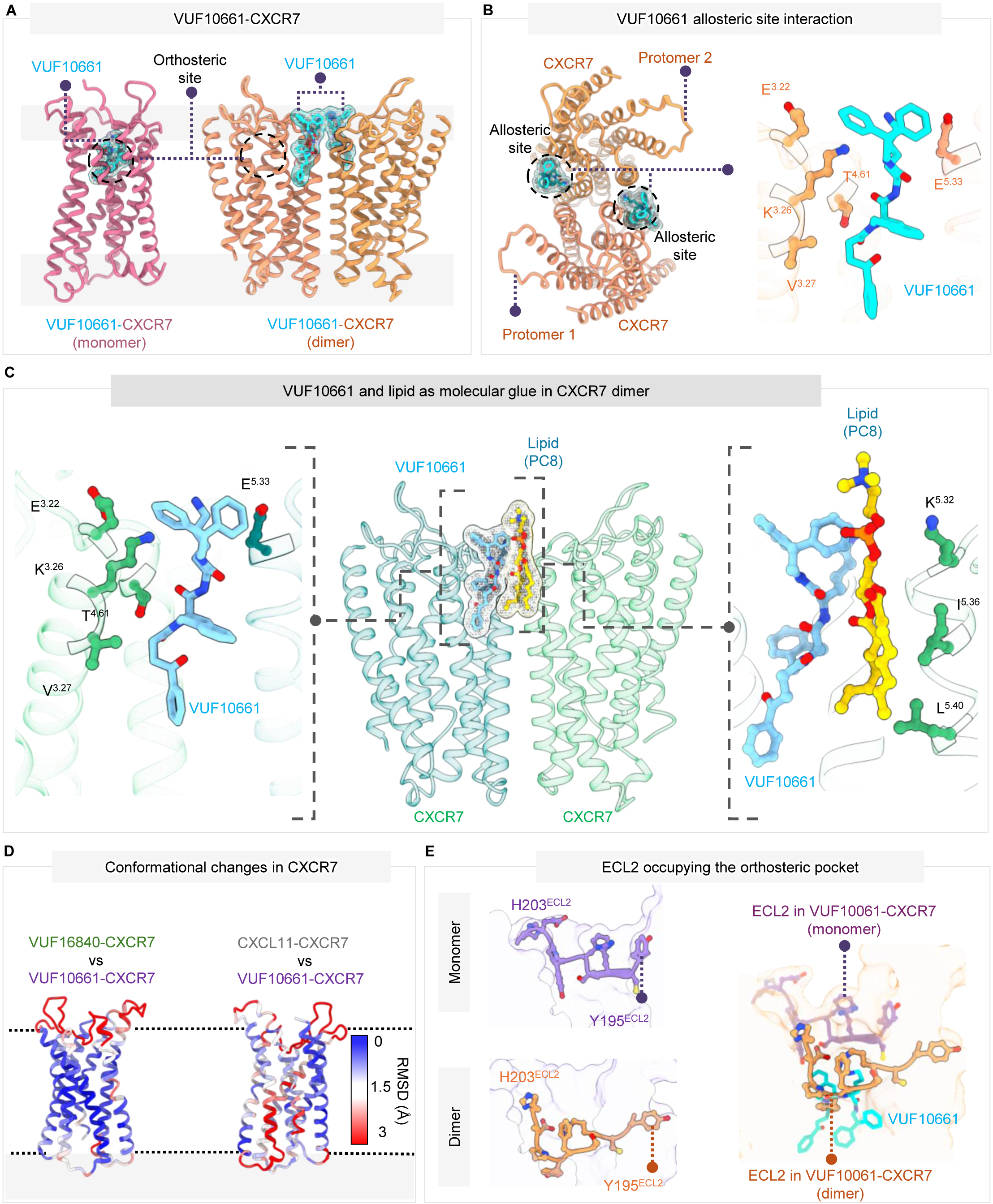
An allosteric site as a molecular glue and an intermediate conformation of CXCR7. **(A)** Ribbon diagram of the VUF10661-bound monomeric and dimeric structures showing the orthosteric and extra-helical allosteric binding sites (side-view) (PDB ID: 9WLF for VUF10661-CXCR7 monomer; 9WLM for VUF10661-CXCR7 dimer). **(B)** Key interactions of VUF10661 with CXCR7 in the extra-helical allosteric binding site as visualized in the cryo-EM structure (PDB ID: 9WLM) and stabilization of VUF10661 in the extra-helical binding pocket by the positioning and interaction of PC at the dimer interface. **(C)** Top-view of VUF10661-bound CXCR7 showing the allosteric sites occupied by VUF101661 on each protomer and the key residues interacting with VUF101661. **(D)** Structural differences between the VUF101661-bound CXCR7 and inactive/active conformations of CXCR7 depicted in terms of RMSD (VUF10661-CXCR7 vs. VUF16840/CXCL11-CXCR7 structures) mapped onto a ribbon representation. **(E)** Distinct ECL2 conformations in the VUF10661-bound monomeric and dimeric CXCR7 structures. A stretch of ECL2 protrudes into the orthosteric binding pocket of the receptor protomers in the dimeric (intermediate) state, thereby, blocks of the pocket for VUF10661 entry.

### CXCR7 as an atypical opioid receptor

In addition to chemokines, CXCR7 is also proposed to recognize opioid peptides and scavenge them to potentially modulate their physiological availability for the canonical opioid receptors^31-33^. In order to assess the specificity of CXCR7 for opioid peptides, we screened a library of >1000 biologically active peptides and analogues, covering a wide variety of ligand families including >150 orphan peptides, on cells expressing CXCR7 using a Nano-BRET-based ligand competition assay (**Figure 6A**). We observed that only about a dozen different peptides were able to displace a significant level of CXCL12 from CXCR7, and most of these were opioid peptides from the enkephalin and dynorphin family (**Figure 6A**). We selected a set of these peptides and measured their ability to induce βarr1/2 recruitment in a dose dependent manner, which further confirmed their robust agonism at CXCR7 **(Supplementary Figure S17A-B**).

**Fig. 6:**
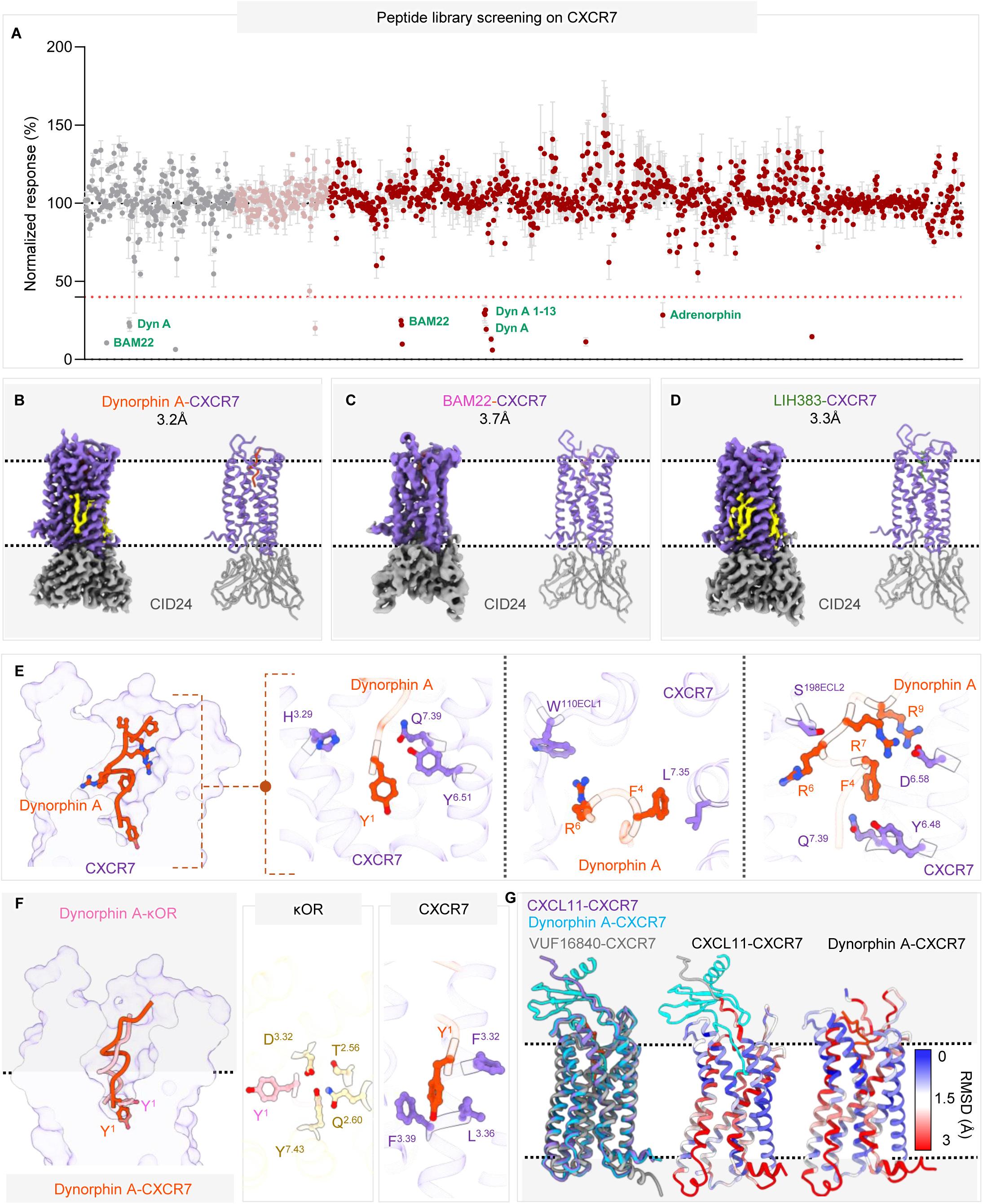
CXCR7 as an atypical opioid receptor. **(A)** Screening of peptide libraries on CXCR7-expressing cells using the displacement of a fluorescently-labeled CXCL12 as a readout. A displacement of 60% of CXCL12 used as a cut-off and the clusters indicated in different colors correspond to three distinct libraries. **(B-D)** Structures of CXCR7 in complex with indicated ligands determined using cryo-EM, stabilized using CID24 as indicated. **(E)** Dynorphin A stabilizing interactions in the CXCR7 orthosteric pocket. (**F),** Relative binding depth and key interactions of Dynorphin A with respect to Dynoprhin A in the orthosteric pocket of KOR and CXCR7 respectively. **(G),** Ribbon diagram showing overall structural changes in CXCR7 upon activation via endogenous ligand CXCL11 and opioid Dynorphin A.

Next, we reconstituted CXCR7 complexes with dynorphin A (dynorphin), BAM22, and an adrenorphin-based CXCR7-selective synthetic peptide (LIH383), and stabilized them using CID24 for structural analysis (**Supplementary Figure S17C-E)**. We determined the structures using cryo-EM at resolutions ranging from 3.2Å to 3.7Å (**Figure 6B-D**). The technical details of cryo-EM data collection and processing are presented in **Table S1** and **Supplementary Figure S18A-C**, and the cryo-EM densities for the TMs and ligands are presented in the **Supplementary Figure S19A-C**. Overall, these structures reveal a clear binding of the ligands, and the lipid moieties in a similar position as described earlier. As dynorphin-bound CXCR7 structure was resolved at a higher resolution, we primarily used this for subsequent analysis as described below. In this structure, the first ten residues of dynorphin were clearly discernible, which adopted a “Z-shaped” conformation with residues Gly3-Phe4-Leu5-Arg6 taking a helical turn, which penetrates deep into the orthosteric pocket (**Figure 6E**). In particular, the N-terminal tyrosine residue of dynorphin (Tyr1^Dyn^) is oriented vertically downwards, forming an extensive receptor-peptide interface with an interface area of more than 800Å^2^ and buried surface area of about 2,200Å^2^. We observed that three hydrogen bonds and multiple non-bonded contacts stabilize the positioning of dynorphin within the amphipathic orthosteric pocket. Tyr1^Dyn^ is nestled within a hydrophobic patch formed by Phe124^3.32^, Leu128^3.36^, and Phe129^3.37^ of CXCR7 (**Figure 6F**). Molecular interactions that anchor dynorphin include the cationic hydroxyl of Tyr1^Dyn^ forming ionic bonds with His121^3.29^ and Gln301^7.39^, while Tyr268^6.51^ engages with the backbone nitrogen of Tyr1^Dyn^ (**Figure 6E**). The Phe4^Dyn^ residue participates in hydrophobic interactions with Leu297^7.35^, while the guanidino group of Arg6^Dyn^ forms a cation-π interaction with Trp110^ECL1^ (**Figure 6E**). In addition, Arg7^Dyn^ and Arg9^Dyn^ establish hydrogen bonds with Asp275^6.58^ in CXCR7 **Figure 6E**).

A direct comparison of dynorphin-bound CXCR7 with the inverse-agonist-bound CXCR7 displays similar conformational changes in the receptor as that observed for CXCL11, indicating that dynorphin is able to fully activate the receptor (**Figure 6G**). In line with this interpretation, the structures of CXCR7 in complex with dynorphin and CXCL11 look nearly-identical with an overall rmsd of <1Å (**Supplementary Figure S20A)**. Moreover, dynorphin and CXCL11 are positioned in the orthosteric binding pocket at comparable depths with similar spatial placement of the terminal ligand residues (Tyr1 in dynorphin vs. Phe1 in CXCL11), leading to similar interaction patterns (**Supplementary Figure S20A**). However, the three residues following the two terminal residues i.e. Gly3-Phe4-Leu5 in dynorphin-CXCR7 and Met3-Phe4-Lys5 in CXCL11, deviate by ∼5.5Å although the subsequent stretches of both peptides realign again within the orthosteric pocket (**Supplementary Figure S20A**). While the backbone atoms of Arg7 and Arg9 in dynorphin occupy similar sites as Arg6 and Arg8 in CXCL11, their side chains are oriented in opposite directions, thereby, engaging different receptor residues. Specifically, Arg6 and Arg9 in dynorphin interact with Asp275^6.58^ in CXCR7 via hydrogen bonds, whereas Arg6 and Arg8 in CXCL11 engage with Asp179^4.60^ and Ile279^6.62^ in the receptor via a hydrogen bond and hydrophobic interaction, respectively (**Supplementary Figure S20B**). These structural observations underline an overall conserved mode of interaction of dynorphin and CXCL11 with CXCR7 with respect to the orthosteric site, and observed differences in their interaction networks are likely to fine-tune their respective potencies for βarr recruitment.

Structural comparison of dynorphin-bound CXCR7 with that of kappa-opioid receptor (kOR)^53^ reveals some interesting patterns of ligand-interaction and receptor conformation. Overall, the two receptors are similar to each other in dynorphin-bound conformation with an overall rmsd of <1Å (**Supplementary Figure S20C)**, and dynorphin also penetrates to a similar depth in the orthosteric binding pockets (**Figure 6F**). However, the side-chain rotamer of Tyr1 in dynorphin differs significantly. In kOR, Tyr1 points in an upward direction while in CXCR7, it points downward, which in turn, results in a divergent interaction network (**Figure 6F**). Despite this deviation at the N-terminus, the other end of dynorphin appears to converge in a similar fashion on the two receptors as reflected by an identical interaction of Arg9 in dynorphin with Asp297^6.58^ and Asp275^6.58^ in the kOR and CXCR7, respectively (**Supplementary Figure S20C-D)**. Importantly, distinct structural arrangements are also evident in the intracellular loops (ICLs) of the two receptors in complex with dynorphin. For example, in the dynorphin-bound kOR structure, the ICL1, ICL2, and ICL3 extend outward from the receptor core, creating a wide cytoplasmic pocket that accommodates heterotrimeric G-proteins (**Supplementary Figure S20E)**. On the other hand, in the dynorphin-bound CXCR7 structure, these ICLs shift linearly towards the receptor core, resulting in a constricted cytoplasmic pocket as observed in the other agonist-bound structures **(Supplementary Figure S2E)**.

The other two opioid peptides namely BAM22 and LIH383 also exhibit an overall similar binding to CXCR7. For example, similar to dynorphin, residues “Gly3-Phe4-Met5-Arg6” of LIH383 form a helical turn, and the overall conformation of LIH383 within the orthosteric pocket closely mirrors that of dynorphin and engages a comparable buried surface area (**Supplementary Figure S20F)**. Specifically, Phe1 in LIH383 penetrates deepest into the orthosteric pocket, with its phenyl ring occupying a spatial position analogous to that of Tyr1 in dynorphin, and the following residues also interact in analogous manner to that of dynorphin, leading to receptor activation as assessed by outward movement of TM6 and TM7 (**Supplementary Figure S20G)**. Taken together, these findings unequivocally establish CXCR7 as an atypical opioid receptor and thereby, expand the opioid receptor subfamily beyond the conventional receptors.

### A pivotal ligand-recognition switch in the chemoattractant receptors

The analysis of CXCR7 structures expectedly revealed several residues that are involved in ligand interactions, and some of these appear to be mostly conserved across different ligands while others seem to show a ligand-specific pattern of interaction (**Figure 7A**). Of these, Tyr268^6.51^ stands out in particular as it appears to make a contact with nearly every ligand used here for structural analysis. We generated a series of alanine mutants of CXCR7 by replacing the key residues involved in ligand-interaction, and measured agonist-induced βarr recruitment (**Figure 7B and Supplementary Figure S21A-B**). We observed that the mutation of some of the residues such as Tyr268^6.51^ affected βarr recruitment across all ligands while the others impacted only selected ligands (**Figure 7B and Supplementary Figure S21C-H**). For example, the mutation of Leu128^3.36^ resulted in a significant attenuation of βarr recruitment for CXCL11 and dynorphin but not for CXCL12 and VUF11207. These findings corroborate the structural observations, and underscore ligand-specific receptor contacts potentially fine-tuning downstream functional responses. Considering the robust impact of Tyr268^6.51^ mutation, we explored if this residue is conserved across other chemokine and non-chemokine peptide receptors. We observed that the majority of the chemokine receptors and all three complement anaphylatoxin receptors harbor Tyr^6.51^, and it also makes a robust interaction with the corresponding natural agonists of these receptors (**Figure 7C**). On other hand, only very few of the non-chemokine peptide receptors harbor the conserved Tyr^6.51^ in TM6. Taken together with site-directed mutagenesis data and structural analysis, we propose that Tyr^6.51^ serves as a pivotal switch for ligand-binding and activation of a broad repertoire of chemo-attractant receptors, underscoring a previously unknown conserved structure feature of these receptors.

**Fig. 7:**
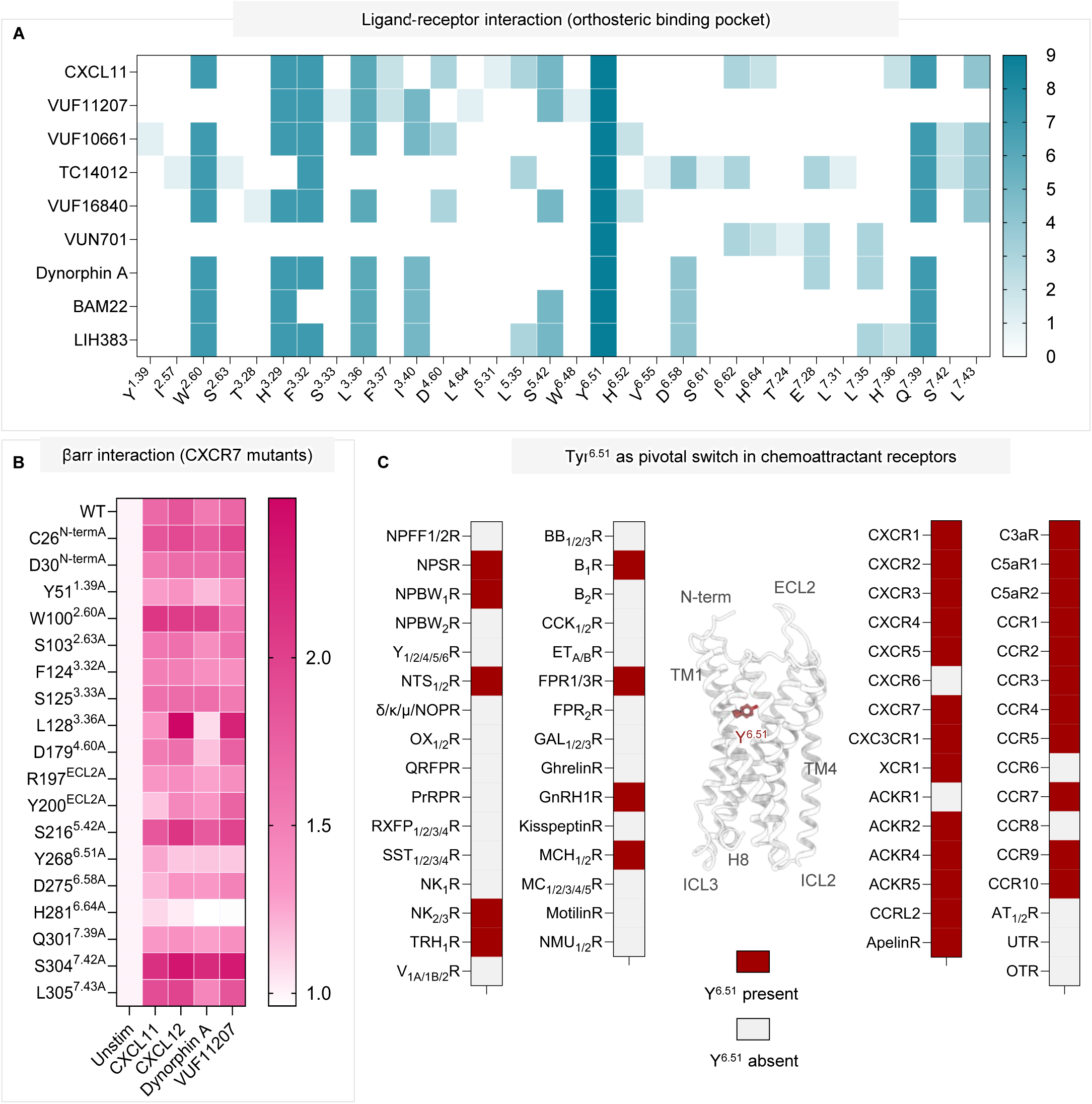
A pivotal tyrosine switch in CXCR7 and chemoattract receptors. **(A)** A heatmap representation depicting the ligand-interacting residues in the TM domain of CXCR7 based on structural snapshots determined here. Color-coding indicates the relative frequencies of the interaction. **(B)** Ligand-induced βarr-recruitment for CXCR7 mutants measured using NanoBiT-based bystander assay in transfected HEK-293T cells presented as a heatmap (mean; n=3; fold-normalized with basal; CXCL11, 1μM; CXCL12, CXCL11, 1μM; VUF11207, 10μM; dynorphin, 31.6μM.**(C)** Sequence analysis of peptide GPCRs uncovers a broadly conserved occurrence of Tyr6.51 in TM6 of the chemoattractant receptors.

## Discussion

The conceptual framework of biased agonism has redefined GPCR pharmacology, signaling, and therapeutic design considering the inherent potential to minimize the side-effects of GPCR-targeting drugs^51^. Biophysical and structural studies using synthetic biased ligands and their cognate receptors such as the angiotensin II subtype 1 receptor (AT1R) have elucidated distinct conformational changes in the receptor as an underlying principle to impart signaling-bias^12–14^. On the other hand, the molecular basis of distinct activation of naturally-encoded, signaling-biased 7TMRs, also referred to as Arrestin-Coupled Receptors (ACRs), which lack measurable G-protein activation, remains mostly speculative. Earlier studies have unequivocally demonstrated a lack of functional engagement of heterotrimeric G-proteins to ACRs, and have suggested distinct conformational signatures in βarrs upon their interaction with ACRs vs. GPCRs^19^. A previous study on CXCR7 has proposed distinct conformation and structural features of the 2^nd^ and 3^rd^ intracellular loops (ICL2 and ICL3) as a plausible mechanism to preclude G-protein-coupling^29^. However, replacing these loops either individually or in combination with that of a prototypical chemokine receptor, CXCR2, did not allow any measurable gain in G-protein-coupling. Therefore, it is likely that additional structural features may contribute to the inherent defect in G-protein activation. In fact, our study reveals a constricted intracellular pocket in CXCR7, resulting particularly from a relatively smaller movements of TM5 and 6 upon activation, compared to other GPCRs. In addition, unlike prototypical GPCRs, the lack of positively charged patch on the intracellular surface in CXCR7 is also readily apparent. Taken together, these distinctive features in activated CXCR7 may not be suitable for efficient docking and alignment of the α5 and αN helices of the Gα subunit, respectively, resulting in the lack of G-protein activation. These observations provide a testable hypothesis for future experiments, and may also guide the investigation on other ACRs going forward (**Figure 8**).

**Figure 8.**
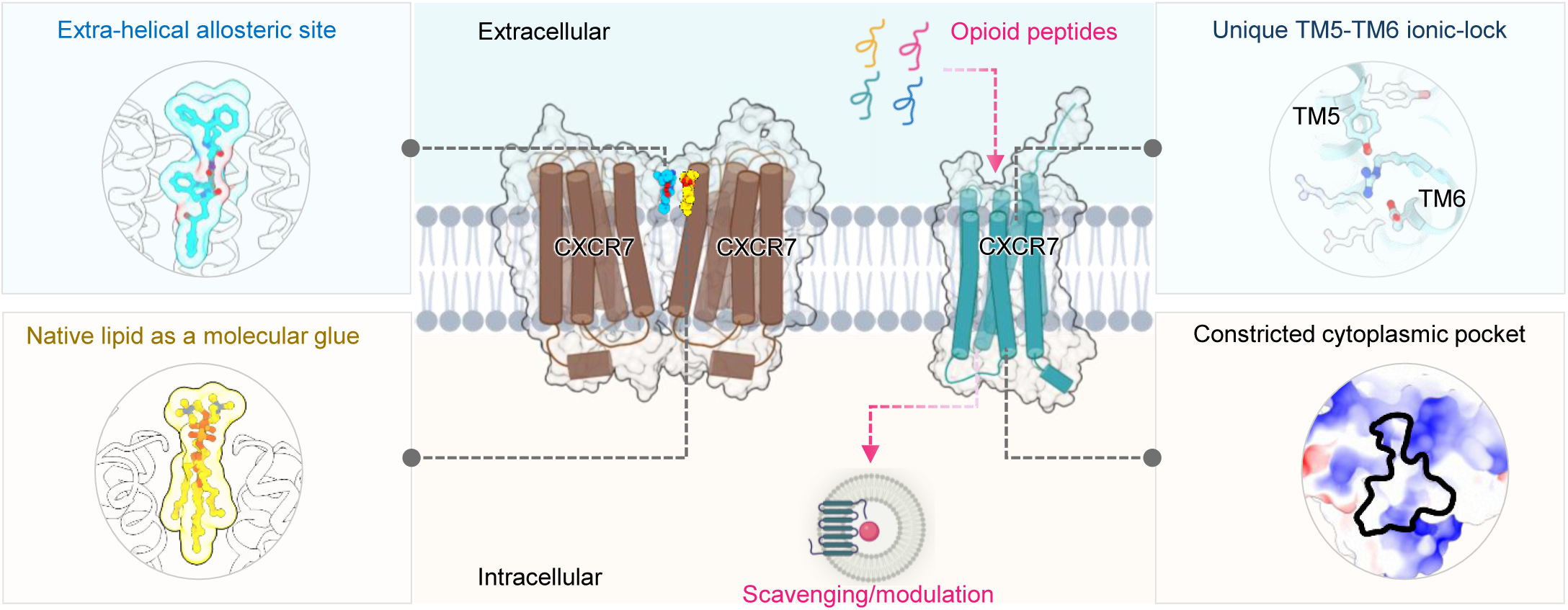
Atypical activation and functional divergence of CXCR7. A schematic illustration depicting a unique activation mechanism, constricted cytoplasmic pocket, allosteric site for a small molecule agonist, dimerization, and recognition of endogenous opioid peptides.

The functional specialization of ACRs in terms of selective βarr-coupling potentially argues for a distinct molecular mechanism of activation compared to prototypical GPCRs, possibly at the level of receptor conformation and micro-switches. This is further substantiated for CXCR7 considering that broadly conserved GPCR sequence motifs such as DRY and NPxxY are conserved in this receptor. Due to the lack of CXCR7 structure in an inactive state, previous inferences about receptor activation have been drawn using the inactive state structures of other chemokine receptors as a reference. However, this is likely to provide only a close approximation at best, and therefore, should be interpreted with caution. For example, a previous study using HDX-MS and site-directed mutagenesis has proposed that Arg^3.50^ in TM3 of CXCR7 may form an ionic lock with Glu^6.29^ in TM6, which serves as an alternative to broadly conserved TM3-TM6 ionic in GPCRs formed between Arg^3.50^ in TM3 and Glu^6.30^ in TM6^43^. Our study now demonstrates that the formation of TM3-TM6 ionic lock in CXCR7 is structurally not plausible, at least based on the receptor conformations captured here using cryo-EM. This, in turn, raises the possibility of alternative inter-helical restraints to maintain the receptor in its basal state. Indeed, CXCR7 structures highlight two novel inter-helical ionic-locks spanning TM5-TM6 and TM7-H8, which are present in the antagonist/inverse-agonist-bound states of the receptor, but are disrupted upon agonist-binding and receptor activation. This observation also aligns with the previous experimental data where the mutation of Glu^6.29^ to alanine imparts higher constitutive activity of CXCR7 compared to the wild-type receptor^43^. We also note that unlike CXCR7, other ACRs lack the conserved sequence motifs such as DRY and NPxxY, and thus, may utilize an analogous mechanism as described here. However, direct structural visualization and biochemical analysis of these additional ACRs is essential to probe this hypothesis.

Dimerization of class A GPCRs are well documented in cellular context using proximity-based assays, and in some cases, also appear to have a direct functional relevance^44–46^. There is a growing list of GPCR cryo-EM structures that also capture GPCR-G-proteins and GPCR-βarr complexes in dimeric arrangements^6^. While some of these may simply represent concentration-dependent non-physiological dimers, others appear to use a specific interface and may have direct functional consequences. On the other hand, direct visualization of distinct dimeric interface and/or arrangements adopted by the same receptor, wherein the protomers are stitched together by a native phospholipid originating from the membrane bilayer, is rather scarce. Previous studies using cell-based experiments have reported homo-dimerization of CXCR7 as well as hetero-dimerization with other GPCRs, and some of these also appear to be sensitive to ligand stimulation^47,48^. Moreover, in case of CXCR7-CXCR4 heterodimer, CXCR7 is also proposed to modulate the activation and signaling of CXCR4, presumably by scavenging the ligand and/or βarrs^47,48^. CXCR7 structures resolved here show three distinct dimeric arrangements, each stabilized via native phospholipid molecules positioned at the inter-protomer interface. While two of them utilize primarily TM1-TM1 interface, the inter-protomer rotation are significantly different between these two dimers, and the other dimeric assembly is stabilized by TM4/5-TM4/5 interface. Still however, the overall positioning of the native phospholipid molecule, designated as phosphatidylcholine here based on the resolved EM density, is nearly identical in all three dimers. A striking observation is that all three dimers exhibit an inactive conformation of the receptor although in one of them, the receptor is bound to an agonist albeit at a non-orthosteric binding pocket. On the other hand, all the active state structures are predominantly present as a monomer, which was also reflected in the 2D class averages during data processing. It would be interesting to probe in future studies, especially in cellular context, if the propensity of receptor dimerization correlates with the transition from inactive to active state (**Figure 8**).

Allosteric sites on GPCRs are emerging as a hot-spot for therapeutic targeting as they can be leveraged to modulate receptor signaling, presumably with a better precision than orthosteric binding pocket^54–56^. Binding of VUF10661, a dual agonist of CXCR3 and CXCR7, to an extra-helical allosteric site is intriguing, especially considering that it can also bind to the orthosteric pocket of CXCR7 and CXCR3^37^. Previous studies have also revealed dual binding mode of ligands for the muscarinic M5 receptor^57^ and the bitter taste receptor TASR2^58^. However, the ability of VUF10661 to stabilize an active receptor conformation when occupying the orthosteric pocket vs. an intermediate, inactive-like conformation from the allosteric site is unique and first-of-its-kind efficacy switch to the best of our knowledge. Future studies are required to probe the unanticipated ECL2 conformation in this structure that blocks the orthosteric site, and thereby, likely pushes VUF10661 to occupy the allosteric site on the receptor. Nonetheless, this self-blocking mechanism is in stark contrast with ECL2-mediated self-activating mechanism observed for a couple of class A GPCRs reported recently^58^, and in turn, underscores a remarkable versatility in structural mechanisms inherently encoded by 7TM scaffold.

The first evidence of opioid peptide recognition by CXCR7 and an ensuring modulation of emotional behaviour circuitry via adrenal-opioid signaling cascade surface more than a decade ago^31^. Subsequently, a comprehensive study reaffirmed the ability of CXCR7 to recognize multiple opioid peptides, which in turn results in βarr recruitment, potentially leading to opioid peptide scavenging^32,33^. This is analogous to chemokine scavenging activity proposed for ACKRs, and may in turn be a natural mechanism to regulate opioid peptide availability in physiological context. Still however, follow-up studies to exploit CXCR7 as a non-canonical opioid receptor from therapeutic perspective have remained mostly non-existent, possibly due to lack of a direct visualization of the binding to opioid peptides to CXCR7. We now demonstrate the CXCR7 not only selectively recognizes primarily opioid peptides from a large library of biologically active peptides, but also that the binding of opioid peptides such as dynorphin and BAM22 fully activates the receptor. These structural observations should alleviate any doubts about CXCR7 being a bonafide receptor for opioid peptides, and thereby, expand the opioid receptor subfamily beyond conventional opioid receptors namely the μ-OR, κ-OR, and δ-OR^53^. More importantly, the structural template of LIH383-CXCR7 should also facilitate the design and optimization of the new generation of synthetic molecules that are more suitable from therapeutic perspective by modulating the opioid concentration under *in-vivo* condition for their cognate receptors.

In summary, our study presents a series of novel insights into atypical CXCR7 activation and functional divergence, and also unravels novel allosteric sites for ligands and native phospholipids. Importantly, our findings also firmly establish CXCR7 as a non-canonical opioid receptor, thereby, expanding the endogenous repertoire of opioid receptors beyond the conventional members. These findings have direct and immediate implications to better understand the framework of biased GPCR signaling and designing novel therapeutics.

## Acknowledgements

Research in A.K.S.’s laboratory is supported by the Senior Fellowship of the DBT Wellcome Trust India Alliance (IA/S/20/1/504916) awarded to A.K.S., the Science and Engineering Research Board (SPR/2020/000408 and IPA/2020/000405), the Indian Council of Medical research (F.NO.52/15/2020/BIO/BMS), and IIT Kanpur. A.K.S. is the Sonu Agrawal Memorial Chair Professor. All the cryo-EM data collection described here was carried at the ANRF-sponsored National cryo-EM Facility at IIT Kanpur (IPA/2020/000405). This study was also supported in part by the Luxembourg Institute of Health (LIH) through the NanoLux Platform, the Cancer Foundation Luxembourg, Luxembourg National Research Fund (INTER/FNRS CXCL12 20/15084569, CORE IMPACTT C23/BM/18068832, INTER/AUDACE/25/19352183 and AFR Hope-ioid 17191672) to AC, Natural Science Foundation of China (grant #22077012) to XC, and a Wellcome Trust grant (number 221795/Z/20/Z) to CVR. We thank Dr. Martin Gustavsson for kindly providing the CID24 expression plasmid and Calvin D’Souza for help with CID24 purification.

## Authors’ contributions

NB and AR purified the receptor, prepared, and characterized the complexes for cryo-EM analysis with help from MK and AD; MG and RB processed the cryo-EM data, determined the structures, and carried out structural analysis with help from NR; SM and AD characterized the ligands and receptor mutants in cellular assays with help from SS and DT; XG synthesized and characterized VUF16840 with help from HG under the supervision of XC; MM and CBP carried out the peptide library screening under the supervision of AC; all authors contributed to manuscript writing and editing; AKS supervised and coordinated the overall study.

## Data availability

The cryo-EM maps and structures have been deposited in the PDB and EMDB with accession numbers 9WLD and EMD-66055 for CXL11-CXCR7-CID24, 9WLE and EMD-66056 for VUF11207-CXCR7-CID24, 9WLF and EMD-66057 for VUF10661-CXCR7-CID24, 9WLL and EMD-66065 for Dynorphin-CXCR7-CID24, 9WLJ and EMD-66061 for LIH383-CXCR7-CID24, 9WLH and EMD-66059 for TC14012-CXCR7-CID24, 9WLK and EMD-66062 for BAM22-CXCR7-CID24, 9WLM and EMD-66066 for VUF10661-CXCR7-dimer, 9WLG and EMD-66058 for VUF16840-CXCR7, 9WLI and EMD-66060 for VUN701-CXCR7-monomer, and 9WLO and EMD-66067 for VUN701-CXCR7-dimer respectively. The accession codes of other PDB coordinate files referenced in this study are 7SK3 (CXCL12-CXCR7-Fab), 7SK5 (CXCL12-CXCR7-Fab), 7SK9 (CCX662-CXCR7-Fab), 8HNK (CXCL11-CXCR3), 8F7W (Dynorphin-KOR-Gi), 8XYI (VUF10661-CXCR3), 8IC0 (CXCL8-CXCR1-Gi), 6LFM (CXCL8-CXCR2-Gi), 8XWV (CXCL1-CXCR2-Go), 8XVU (CXCL2-CXCR2-Go), 8XWA (CXCL1-CXCR2 receptor-ligand focused map), 8XWF (CXCL3-CXCR2-Go), 8XWM (CXCL6-CXCR2-Go), 8XWN (CXCL8-CXCR2), and 8XWS (CXCL5-CXCR2).

## Materials and methods

### Human cell line

HEK293T cells used in this study were purchased from ATCC (Cat. No. CRL-3216). They were cultured in DMEM, supplemented with 10% FBS, at 37°C under 5% CO₂. The cell line was routinely monitored under the microscope for appropriate morphology, but they were not authenticated. In this study no knockout or knockdown cell lines were generated.

### Insect cell line

*Sf*9 cell line was obtained from Expression Systems (Cat. No: 94-001F), maintained in a shaker incubator at 27°C with 135 rpm shaking speed, and sub-cultured in protein-free insect cell media purchased from Gibco (Cat. No: 10902-088). The cells were regularly inspected under microscopy for appropriate morphology, but they were not authenticated.

### General reagents and chemicals

Most standard molecular biology reagents were purchased from Sigma-Aldrich, unless otherwise specified. Dulbecco’s Modified Eagle Medium (DMEM), trypsin-EDTA, Penicillin-Streptomycin solution, Phosphate-Buffered Saline (PBS), Fetal Bovine Serum (FBS), and Hanks’ Balanced Salt Solution (HBSS) were purchased from ThermoFisher Scientific. HEK-293 cells were purchased from ATCC (Cat. no: CRL-3216) and maintained in 10 cm dishes (Corning, Cat. no: 430167) in DMEM (Gibco, Cat. no: 12800-017) supplemented with 10% (v/v) FBS (Gibco, Cat. no: 10270-106) and 100 U/mL penicillin and 100 µg/mL streptomycin (Gibco, Cat. no: 15140122) at 37°C and 5% CO2. Plasmid constructs for NanoBiT-based assays have been described previously^59^. VUF11207 was synthesized as described previously^60^ while VUF10661 was purchased from Sigma-Aldrich (Cat. No: SML0803) and TC14012 was purchased from MedChemExpress (Cat. no: HY-P1102). The opioid peptides used in βarr recruitment assays and structural studies were obtained from Genscript.

### NanoBiT-based G-protein-coupling and βarr recruitment

The standard methods for plasmid transfection and NanoBiT assays were carried out as described previously^59^. Briefly, HEK-293T cells were seeded in the 10 cm dishes (Corning, Cat. no: 430167) at a density of 3 million, 24 h prior to the transfection. For preparing the transfection mixture, plasmid DNA and polyethyleneimine, linear (PEI) (Polysciences, Cat. no: 23966) were mixed in 1:3 ratio in 500 µL serum-free DMEM (Cellclone, Cat. no: CC3004) in a microcentrifuge tube. This mix was incubated for 10 min. Prior to the addition of the transfection mixture, the FBS-supplemented DMEM of the cell culture plate was replaced with serum-free DMEM. Subsequently, the transfection mix was added to the cells, and serum-free DMEM was replaced with FBS-supplemented DMEM 6 h post-transfection.

To measure ligand-induced ꞵarr recruitment, cells were transfected with the indicated receptors tagged at the C-terminus with SmBiT and N-terminally LgBiT-tagged ꞵarr1/2. After 16-18 h of transfection, cells were washed with 1X PBS, trypsinized, and resuspended in the assay buffer containing 1X HBSS, 5 mM HEPES, pH 7.4, 0.01% BSA, and 10 µL coelenterazine (GoldBio, Cat. no: CZ05). The cells were seeded in a white, flat-bottom 96 well plate at the density of 0.1 million cells per well, and incubated at 37°C for 90 min, followed by a 30 min incubation at room temperature. Afterwards, basal luminescence values were recorded for three cycles, followed by addition of ligands at indicated concentrations, and luminescence readings were taken for additional ten cycles using a multimode plate reader (LUMIstar/FLUOstar microplate reader, BMG Labtech). The analysis was carried out using GraphPad Prism v 10.4.0 software, taking an average of luminescence values for cycles 5-10. Normalization was carried out by taking luminescence values of the lowest dose as 1.

CXCR7 mutants were generated on wild-type template using a site-directed mutagenesis kit (NEB, Cat no. E0554S) as indicated, and all constructs were verified by sequencing (Macrogen). Subsequently, βarr recruitment was measured as described above.

To measure G-protein dissociation, cells were transfected using the indicated receptor constructs and heterotrimeric G-proteins, and 16-18 h post-transfection, cells were harvested by washing with 1X PBS followed by trypsinization. Subsequently, cells were resuspended in the assay buffer containing 1X HBSS, 5 mM HEPES, pH 7.4, 0.01% BSA and 10 µL coelenterazine (GoldBio, Cat. no: CZ05), and seeded in a white, flat-bottom 96-well plate at a density of 0.1 million cells per well. The plate was incubated at 37°C for 90 min followed by incubation at room temperature for an additional 30 min. Post-incubation, basal luminescence reading was taken for three cycles, followed by ligand stimulation at indicated concentrations and luminescence was recorded for ten additional cycles using a multimode plate reader (LUMIstar/FLUOstar microplate reader, BMG Labtech). Data analysis was carried out using GraphPad Prism v 10.4.0 software, and the luminescence value at 10 min post-ligand stimulation was corrected for basal signal, followed by percentage normalization in terms of the decrease in luminescence value.

In cell-based assays, the surface expression of the receptors was measured using a previously described whole cell-based surface ELISA assay^61^. Briefly, cells were seeded in a poly-D-lysine-coated 24 well plate at a density of 0.2 million cells per well, and allowed to adhere for 24 h. Afterwards, growth media was removed, cells were washed with ice-cold 1X Tris-buffered saline (TBS), and then fixed using 4% paraformaldehyde for 20 min on ice. Subsequently, cells were washed three times with 1X TBS, incubated with 1% BSA (w/v, prepared in 1X TBS) for 1 h at room temperature, followed by 1 h incubation with anti-FLAG M2-HRP (Sigma-Aldrich, Cat. no: A8592; RRID:AB_439702) at 1:10,000 dilution (prepared in 1% BSA). Then, cells were washed three times with 1% BSA, followed by addition of TM Ultra TMB-ELISA (Thermo Fisher Scientific, Cat. no: 34028), the reaction was quenched using 1 M H2SO4. Signal was measured at 450 nm using the PerkinElmer VictorTM X4 multimode plate reader, and to normalize the signal, Janus Green staining was performed (Sigma-Aldrich, Cat. no: 201677) with absorbance measured at 595 nm. Surface expression was calculated by taking a ratio of absorbance at 450 nm and 595 nm, and normalizing with respect to mock-transfected cells.

### Expression and purification of CXCR7

CXCR7 was expressed and purified using baculovirus-mediated infection of *Sf*9 cells following the protocols published previously^62–65^. Briefly, a full-length CXCR7 construct with N-terminal FLAG-tag was generated using baculovirus DNA (Cat. No: 91-002). Cells were harvested 72 h post-infection, homogenized in a hypotonic buffer (20 mM HEPES pH 7.4, 20 mM KCl, 10 mM MgCl₂, 1 mM PMSF, 2 mM benzamidine), followed by a hypertonic buffer (20 mM HEPES pH 7.4, 20 mM KCl, 10 mM MgCl₂, 1 M NaCl, 1 mM PMSF, 2 mM benzamidine). Subsequently, the receptor was solubilized in lysis buffer (20 mM HEPES pH 7.4, 450 mM NaCl, 1 mM PMSF, 2 mM benzamidine, 0.1% cholesteryl hemisuccinate, 2 mM iodoacetamide) containing 1% L- MNG (Anatrace, Cat. no: NG310) for 2 h at 4°C. Cell lysate containing the solubilized receptor was centrifuged (∼48,000 X g for 30 min), filtered, and loaded onto pre-equilibrated M1-FLAG column. The column was washed alternately with low salt buffer (20 mM HEPES pH 7.4, 150 mM NaCl, 2 mM CaCl2, 0.01% cholesteryl hemisuccinate, 0.01% L-MNG) and high salt buffer (HSB; 20 mM HEPES pH 7.4, 350 mM NaCl, 2 mM CaCl2, 0.01% L-MNG). Subsequently, the receptor was eluted using elution buffer (20 mM HEPES pH 7.4, 150 mM NaCl, 0.01% L-MNG, 2 mM EDTA, and 250 ug/mL FLAG peptide), free cysteines were alkylated by incubation with 2 mM iodoacetamide, and any remaining iodoacetamide was quenched with 2 mM L-cysteine. Purified receptor was analyzed by SDS-PAGE, flash-frozen with 10% glycerol, and stored at −80°C until further use.

### Expression and purification of CXCL11

CXCL11 was purified following a previously published protocol^66^. Briefly, *E. coli* (Shuffle) cells were transformed with CXCL11 expression plasmid, and transformed cells were first grown overnight at 30°C in 50 mL of TB media containing 100 μg/mL ampicillin. The primary culture was then used to inoculate 1 L of TB media and grown at 30°C until the OD600 reached 1.2, and subsequently, protein expression was induced with IPTG (1mM). After an additional growth at 20°C for 48 h, cells were harvested and lysed in lysis buffer (50 mM Tris-HCl pH 8, 150 mM NaCl, 6 M Guanidine hydrochloride, pH 8) for 1 h at 4°C followed by sonication at 40% amplitude for 20 min. Cell lysate was then centrifuged at ∼48,000 x g at 4°C for 1 h, supernatant was filtered, and then loaded onto a pre-equilibrated Ni-NTA column. The beads were washed with the same buffer, and the bound protein was eluted with elution buffer (50 mM Tris-HCl pH 8, 150 mM NaCl, 500 mM imidazole pH 8). Eluted protein was incubated with 20 mM DTT for 1 h at room temperature and then added dropwise in renaturation buffer (20 mM Tris-HCl pH 8, 200 mM NaCl, 550 mM Arginine pH 8, 1 mM EDTA, 1 mM reduced glutathione, 0.1 mM oxidized glutathione) followed by an incubation for 48 h at 4°C. After renaturation, the protein solution was dialyzed against 20 mM Tris-HCl pH 8, 150 mM NaCl, and hexa-histidine tag was cleaved using enterokinase in presence of 10 mM CaCl₂ for 16 h at 4°C. The Cleaved CXCL11 was purified on Resource S cation exchange chromatography with elution using a linear gradient of NaCl, and the elution fractions were analyzed by SDS-PAGE. Subsequently, fractions containing purified CXCL11 were pooled, dialyzed overnight against 20 mM HEPES pH 7.4, 150 mM NaCl buffer at 4°C, flash-frozen with 10% glycerol, and stored at −80°C until further use.

### Expression and purification of CID24

CID24 was expressed and purified following previously published protocol^67^. Briefly, CID24 expression plasmid was transformed in *E. coli* BL21 (DE3) cells, and the primary culture was grown in 2XYT media containing 100 μg/mL ampicillin at 37°C and expression was induced with 1 mM IPTG at OD600 of 1.0. The culture was grown for an additional 4 h at 37°C followed by harvesting and lysis of cells in lysis buffer (20 mM HEPES pH 7.4, 500 mM NaCl, 0.5 mM MgCl2, 1 mM PMSF, 2 mM Benzamidine, and 0.5% (v/v) Triton X-100). Cell lysis was performed using sonication for 20 min at 40% amplitude, and the lysate was heated at 65°C for 30 min. Subsequently, lysate was centrifuged at ∼48,000 x g for 40 min at 4°C, supernatant was clarified by filtration, and then loaded onto a pre-equilibrated Capto-L (Cytiva) column. The beads were washed with washing buffer (20 mM HEPES pH 7.4, 500 mM NaCl), Fab was eluted with 0.1 M acetic acid, and immediately neutralized with 10% (v/v) 1M HEPES pH 7.4. Purified protein was dialyzed against 20 mM HEPES pH 7.4, 150 mM NaCl buffer, analyzed using SDS-PAGE, and flash-frozen with 10% glycerol and stored at −80°C until further use.

### Expression and purification of VUN701

The gene encoding VUN701 was cloned in pET-22b (+) vector with a hexa-histidine tag at the N-terminus, and transformed in *E. coli* (Rosetta) cells. Freshly transformed cells were used to inoculate primary culture at 37°C in 50 mL of 2XYT media supplemented with 100 μg/mL ampicillin. The culture was grown overnight and further expanded in 1L 2XYT media and grown in 37°C until OD600 reached 1.1. Afterwards, the culture was induced with 100 μM IPTG and incubated for 16 h at 18°C. Subsequently, cells were harvested and resuspended in lysis buffer (20 mM HEPES pH 8, 150 mM NaCl, 1 mM PMSF, 2 mM Benzamidine and 1 mg/mL Lysozyme) and homogenized for 1 h at 4°C. Cells were lysed by ultrasonication at 40% amplitude for 20 min, and the lysate was centrifuged at ∼48,000 x g for 40 min at 4°C. The supernatant was filtered and loaded onto a pre-equilibrated Ni-NTA column, and the column was washed with 20 mM HEPES pH 8, 150 mM NaCl, and 30 mM imidazole pH 8 to remove unbound protein. Subsequently, bound protein was eluted with 20 mM HEPES pH 8, 150 mM NaCl, and 500 mM imidazole pH 8, and dialyzed against 20 mM HEPES pH 8, 150 mM NaCl at 4°C for 16 h. After dialysis, protein was loaded onto a pre-equilibrated HiTrap Q FF column, fractions corresponding to VUN701 were pooled, and dialyzed against 20 mM HEPES pH 7.4 and 150 mM NaCl. Finally, the purified protein was flash-frozen with 10% glycerol and stored at −80°C until further use.

### Chemical synthesis of VUF16840

#### Chemical reagents and instruments

All chemical reagents were purchased from Energy Chemicals and Aladdin Chemicals (Shanghai, China) and used as received. All moisture-sensitive reactions were carried out using anhydrous solvents and under nitrogen atmosphere. ^1^H and ^13^C NMR spectra were recorded on a Bruker Avance III spectrometer (400 MHz) using TMS as the internal standard. High resolution mass spectrometry (HRMS) was performed on Agilent Technologies 6540 UHD Accurate-Mass Q-TOF. Flash column chromatography was performed with silica gel (300-400 mesh).

1-(tert-Butyl) 3-methyl (S)-4-((1-phenylethyl)amino)-5,6-dihydropyridine-1,3(2H)-dicarboxylate(2): To a solution of 1-(tert-butyl) 3-methyl 4-oxopiperidine-1,3-dicarboxylate (5 g, 19.4 mmol) and Na2SO4 (5.53 g, 38.8 mmol) in 2-methyltetrahydrofuran (2-MeTHF, 25 mL) was added (S)-1-phenylethan-1-amine (2.6 g, 21.4 mmol) at rt. After being stirred at rt for 18h, the reaction mixture was filtered via a celite pad, and the filter cake was washed with 2-MeTHF (2×5 mL). The combined filtrates were collected and kept under N₂, used for next step without further purification.

1-(tert-Butyl) 3-methyl (3R,4S)-4-(((S)-1-phenylethyl)amino)piperidine-1,3-dicarboxylate TFA salt(3): NaBH4 (1.5 g, 23.3 mmol) in 2-MeTHF (15 mL) was cooled to −5°C, followed by dropwise addition of TFA (3.1 g, 27.2 mmol) over 5 min. The freshly prepared compound 2 2-MeTHF solution described above was added dropwise at −15°C. After being stirred at −15°C for 15 min, the reaction mixture was quenched by adding 4 M NaOH (18 mL). The resulting mixture was allowed to warm to ambient temperature, and then extracted with 2-MeTHF (3 × 30 mL). The combined organic layers were washed with brine, dried over Na2SO4, and concentrated. The residue was dissolved in TBME (40 mL) at 65°C, then cooled to ambient temperature, and was treated with TFA (2.67 g, 23.3 mmol) to induce crystallization. The precipitate was filtered, washed with 2-MeTHF/TBME (1:1, 2 × 30 mL), and dried under vacuum at 55°C to afford 3 as a white solid (4.53 g, 63% yield for two steps). ^1^H NMR (400 MHz, MeOD): δ 7.56-7.48 (m, 5H), 4.69-4.41 (m, 2H), 3.80 (s, 3H), 3.81-3.24 (m, 5H), 2.90-1.60 (m, 4H), 1.69 (d, J = 6.8 Hz, 3H), 1.43 (s, 9H).^13^C NMR (101 MHz, MeOD): δ 173.00, 155.10, 137.20, 130.93, 130.68, 128.81, 81.71, 58.50, 56.31, 42.15, 28.49, 26.90, 20.24.

1-(tert-Butyl) 3-methyl (3R,4S)-4-aminopiperidine-1,3-dicarboxylate TFA Salt(4): A mixture of 3 (1.9 g, 5.2 mmol) and 20% Pd/C (0.42 g, 0.78 mmol) in MeOH (20 mL) was evacuated and purged with hydrogen gas (3 times). After being stirred for 3 h under hydrogen, the reaction mixture was filtered through a celite pad, and the filter cake was washed with MeOH (2×15 mL). The combined filtrates were concentrated and recrystallized from TBME (20 mL), giving 4 (1.09 g, 81% yield) as a white solid. ^1^H NMR (400 MHz, DMSO-d₆): δ 4.00 (m, 2H), 3.89 (s, 3H), 3.65 (m, 3H), 3.49 (dt, J = 10.5, 4.1 Hz, 1H), 3.23 (dd, J = 14.0, 3.8 Hz, 1H), 3.11–2.75 (m, 2H), 1.90 (m, 1H), 1.76–1.72 (m, 1H), 1.37 (s, 9H).

1-(tert-Butyl) 3-methyl (3R,4S)-4-(5-(2,4-difluorophenyl)isoxazole-3-carboxamido)piperidine-1,3-dicarboxylate(5): A mixture of 5-(2,4-difluorophenyl)isoxazole-3-carboxylic acid (455 mg, 2 mmol) and HOAt (330 mg, 2.4 mmol) in DMF (8 mL) were stirred at 0°C for 10 min. Amine 4 (720 mg, 2 mmol) and NMM (142 mg, 1.4 mmol) were then added at 0°C, followed by EDCI (465 mg, 2.4 mmol). The ice bath was removed, and the reaction mixture was then stirred at ambient temperature for 12 h. After the solvent and volatiles were evaporated under the reduced pressure, H2O (10 mL) was added, and the mixture was extracted with EtOAc (3×15 mL). The combined organic phases were washed with brine (20 mL) and dried (anhydrous Na2SO4). The crude products were purified by flash column chromatography (eluting with petroleum ether/EtOAc, 6:1) to afford 5 (580 mg, 62% yield) as a white solid. ^1^H NMR (400 MHz, MeOD): δ 8.08-7.97 (m, 1H), 7.21 (m, 2H), 7.05 (s, 1H), 4.52-4.27 (m, 2H), 4.04 (brs, 1H), 3.71 (s, 3H), 3.37-3.26 (m, 4H), 2.25-2.05 (m, 1H), 1.78 (m, 1H), 1.46 (s, 9H), 1.36-1.25 (m, 1H). ^13^C NMR (101 MHz, MeOD): δ 174.09, 161.00, 160.77, 130.90, 114.20 (d, J = 22.1 Hz), 113.50, 106.57, 106.30 (d, J = 24.5 Hz), 103.66, 81.89, 52.95, 44.97, 29.12.

(3S,4S)-1-(tert-Butoxycarbonyl)-4-(5-(2,4-difluorophenyl)isoxazole-3-carbox-amido)piperidine-3-carboxylic acid(7)A 22% KOⁱPr solution (1.11 g, 2.5 mmol) in ^i^PrOH (25 mL) was azeotropically dried via co-evaporation at 70°C. ^i^PrOAc (127 mg, 1.2 mmol) was added, followed by 5 (580 mg, 1.2 mmol) at rt. The reaction mixture was stirred at rt for 2 h, and the solution containing the resulting ester 6 was then cooled to −5°C with an ice-bath. 2 M KOH (1.8 mL, 3.6 mmol) was added to the cooled mixture, and the whole reaction mixture was stirred at the same temperature for 24 h. The pH value of the reaction solution was adjusted to 3.5 by adding 1 M HCl, resulting in some precipitates. The mixture was heated to 75°C until a clear solution formed, and some solid crystallized from the solution upon cooling. The solid was filtered, washed with H₂O/iPrOH (3:1, 2 × 15 mL), and dried at 65°C under vacuum to give 7 (410 mg, 73% yield). ^1^H NMR (400 MHz, MeOD): δ 8.41-7.86 (m, 1H), 7.36-7.14 (m, 2H), 7.12-6.99 (m, 1H), 4.39 (m, 1H), 4.12 (m, 1H), 2.95 (m, 2H), 2.69 (m, 1H), 1.98 (m, 1H), 1.61 (m, 1H), 1.48 (s, 9H). ^13^C NMR (101 MHz, MeOD): δ 174.68, 165.92, 160.53 (d, J = 26.4 Hz), 156.13, 130.27 (d, J = 13.6 Hz), 113.66 (d, J = 22.3 Hz), 106.03 (t, J = 26.2 Hz), 103.09, 81.63, 50.20, 49.90, 28.60. tert-Butyl(3S,4S)-4-(5-(2,4-difluorophenyl)isoxazole-3-carboxamido)-3-((2-methoxyphenethyl)carbamoyl)piperidine-1-carboxylate(8): A solution of 7 (410 mg, 0.9 mmol) and 2-(2-methoxyphenyl)ethan-1-amine (165 mg, 1.1 mmol) in MeCN (8 mL) was treated with NMI (298 mg, 3.6 mmol) at rt. TCFH (383 mg, 1.4 mmol) in MeCN (3 mL) was added dropwise. After being stirred at rt for 30 min, the reaction mixture was diluted with H₂O, heated at 75°C for 45 min, and then allowed to cool to ambient temperature, resulting in formation of some precipitate. The solid was filtered, washed with H₂O/MeCN (4:1, 2 × 10 mL), and dried at 65°C under vacuum to afford 8 (376 mg, 71% yield) as a white solid. ^1^H NMR (400 MHz, DMSO-d6): δ 8.65 (d, J = 8.6 Hz, 1H), 8.09-7.84 (m, 2H), 7.46 (m, 1H), 7.25 (d, J = 2.5 Hz, 1H), 7.13-7.03 (m, 2H), 6.99 (d, J = 7.4 Hz, 1H), 6.85 (d, J = 8.2 Hz, 1H), 6.74 (t, J = 7.4 Hz, 1H), 4.23 (d, J = 11.2 Hz, 1H), 3.69 (s, 3H), 3.46 (m, 4H), 3.20 (t, J = 6.9 Hz, 2H), 2.67 (d, J = 1.3 Hz, 2H), 2.60 (q, J = 7.5 Hz, 1H), 1.88-1.80 (m, 1H), 1.40 (s, 9H). ^13^C NMR (101 MHz, DMSO-d6): δ170.49, 165.42, 164.15, 162.53 (J = 12.5 Hz), 160.38 (J = 12.8 Hz), 159.59, 157.90, 157.77, 157.25, 153.91, 129.86, 127.34 (J = 49.2 Hz), 120.22, 113.03 (J = 20.8 Hz), 112.93, 105.47 (t, J = 26.1 Hz), 102.56, 79.20, 59.90, 55.18, 48.03, 38.65, 38.29, 29.91, 28.10, 20.78, 14.12.

5-(2,4-Difluorophenyl)-N-((3S,4S)-3-((2-methoxyphenethyl)carbamoyl)piperidin-4-yl)isoxazole-3-carboxamide(9): To a solution of compound 8 (170 mg, 0.3 mmol) in ^i^PrOH (4 mL) was added 4 M HCl/^i^PrOH (0.4 mL, 1.2 mmol), and the mixture was stirred at 50°C for 5 h. After the solvent and volatiles were evaporated under the reduced pressure, the residue was dissolved in EtOAc (30 mL), washed with saturated NaHCO3 solution (10 mL) and brine (10 mL), and dried (anhydrous Na2SO4). The crude product was purified by flash column chromatography (eluting with petroleum ether/EtOAc) to afford 9 (61 mg, 43%) as light off-white solid. ^1^H NMR (400 MHz, DMSO-d6): δ 8.96 (d, J = 8.5 Hz, 1H), 8.19 (s, 1H), 7.98 (q, J = 8.0 Hz, 1H), 7.49 (m, 1H), 7.35-7.19 (m, 2H), 7.08 (m, 1H), 6.99 (d, J = 7.3 Hz, 1H), 6.83 (m, 1H), 6.73 (m, 1H), 4.31-4.14 (m, 4H), 3.64 (s, 3H), 3.42-3.12 (m, 2H), 3.11-2.93 (m, 2H), 2.55 (d, J = 9.1 Hz, 2H), 2.01 (m, 1H), 1.82 (m, 1H). ^13^C NMR (101 MHz, DMSO-d6): δ169.22, 164.94, 164.12, 162.50 (d, J = 12.4 Hz), 160.33 (d, J = 12.8 Hz), 159.40, 157.85, 157.71, 157.19, 129.65 (d, J = 20.1 Hz), 127.30 (d, J = 71.8 Hz), 120.28, 113.19, 111.59, 110.63, 105.79, 102.63, 102.54, 55.24, 47.21, 44.54, 44.13, 42.41, 29.76, 28.42, 24.16.

N-((3S,4S)-1-Cyclohexyl-3-((2-methoxyphenethyl)carbamoyl)piperidin-4-yl)-5-(2,4-difluorophenyl)isoxazole-3-carboxamide(VUF16840)A mixture of 9 (316 mg, 0.65 mmol), cyclohexanone (128 mg, 1.3 mmol), MeOH (5 mL) and catalytic amount of AcOH (1 drop) was stirred at rt for 3 h. NaBH4 (49 mg, 1.3 mmol) was then added, and the stirring was continued for an additional 12 h. The reaction was quenched by adding H₂O (3 mL). The resulting mixture was concentrated to dryness, and the residue was extracted with EtOAc (3×10 mL). The combined organic phases were washed with brine (10 mL) and dried (anhydrous Na₂SO₄). The crude product was purified by flash column chromatography (eluting with DCM/MeOH) to afford VUF16840 (116 mg, 32% yield) as a grey solid. ^1^H NMR (400 MHz, DMSO-d6): δ 8.85 (brs, 1H), 8.06-8.01 (m, 2H), 7.58 (ddd, J = 11.5, 9.2, 2.5 Hz, 1H), 7.32 (td, J = 8.6, 2.5 Hz, 1H), 7.19 (d, J = 2.8 Hz, 1H), 7.17-7.09 (m, 1H), 7.05-7.00 (m, 1H), 6.88 (d, J = 8.1 Hz, 1H), 6.77 (m, 1H), 4.05 (m, 1H), 3.69 (s, 3H), 3.27-3.12 (m, 3H), 2.64-2.55 (m, 3H), 2.08-1.95 (m, 3H), 1.81-1.78 (m, 2H), 1.61-1.57 (m, 2H), 1.34-1.28 (m, 5H), 1.24-1.10 (m, 2H). ^13^C NMR (101 MHz, DMSO-d6): δ172.19, 164.23, 160.42, 159.45, 157.68, 157.21, 130.14, 129.88, 129.72, 128.54, 127.65, 126.93, 124.61, 120.21, 113.09 (J = 19.6 Hz), 110.80 (J = 31.0 Hz), 105.56 (J = 25.7 Hz), 102.56, 68.65, 60.83, 55.24, 47.43, 43.41, 38.59, 32.62, 29.84, 28.71, 26.90, 25.57. HRMS (ESI, positive): Calcd. for C31H37F2N2O4 [M+1]+ 567.2783; observed: 567.2787.

### Reconstitution of CXCR7 complexes for structural analysis

For preparing CXCR7 complexes, purified CXCR7 was incubated with 3-5 fold molar excess of the indicated ligands, and approximately 1.5 fold molar excess of CID24, when applicable, for 1.5 hours at room temperature. Following incubation, the protein complex was concentrated using a 100 MWCO concentrator (Cytiva, Cat. no: 28932363) and injected into SuperoseTM 6 Increase 10/300 GL column. Peak fractions corresponding to the target complexes were analyzed by SDS-PAGE, pooled, concentrated to 15-18 mg/mL, and used for freezing cryo-EM grids. In those cases where the cryo-EM grids were not frozen immediately after sample preparation, the samples were flash-frozen in liquid N2 and stored at −80°C until further use.

### Cryo-EM grid preparation and data collection

Purified complex samples were applied onto glow-discharged Quantifoil R1.2/1.3 gold grids (300 mesh, holey carbon) at a protein concentration of 12-18 mg/ml (∼3 μl per grid). Grids were blotted for 4 s at 4°C and 100% relative humidity (blot force 0) using a Vitrobot Mark IV (Thermo Fisher Scientific) and immediately plunge-frozen in liquid ethane. Cryo-EM data were collected on a Titan Krios G4 electron microscope (Thermo Fisher Scientific) operated at 300 kV, equipped with a Gatan K3 direct electron detector and BioQuantum K3 energy filter. Automated data acquisition was performed using EPU software, with movies recorded in counting mode at a nominal magnification corresponding to a pixel size of 0.86 Å. Dose rates were maintained at ∼15.6 e⁻/Å²/s, with a total dose of 55 e⁻/Å² distributed over 50 frames for each 1 s exposure. For the VUN701-CXCR7 monomer complex, movies were recorded at a nominal magnification of 165,000x corresponding to a pixel size of 0.53 Å with a total dose of 75 e⁻/Å². Images were collected across a defocus range of −0.8 to −1.8 μm. The total number of micrographs for each complex are indicated in the **Figure S3-S5 and S18**.

### Cryo-EM data processing

All datasets for CXCR7 complexes were processed following a standardized workflow in cryoSPARC v4.5-4.6, unless otherwise specified. Dose-fractionated movie stacks were corrected for beam-induced motion using the Patch motion correction (multi) sub-program, and contrast transfer function parameters were determined with Patch CTF (multi)^68^. Particles were automatically picked using the blob-picker sub-program, applying a circular mask with a diameter range of 80-160 Å. The auto-picked particles were extracted with a box size of 320 px (fourier cropped to 72 px) and subjected to reference-free 2D classification and heterogeneous refinement to eliminate ice contamination and distorted particles. Particles corresponding to the best 3D class were selected and re-extracted using a box size of 320 px (fourier cropped to 256 px) and subjected to non-uniform (NU) refinement and local refinement. A complete data processing pipeline including the number of particles selected at different stages are presented in Figure S3-S5 and S13. Local resolution estimations for all cryo-EM maps were performed using the LocRes module in cryoSPARC, with half-maps provided as input. Protein-protein interfaces were analyzed using PDBsum^69^.

### Model building and refinement

Coordinates for CXCR7 were generated using SWISS-MODEL and the atomic coordinates of CID24 were derived from the previously determined structure of the CCX662-CXCR7-CID24 complex (PDB: 7SK9)^29^. Initial models for the chemokines were generated with SWISS-MODEL, employing the previously determined structure of CXCL11 (PDB: 1RJT) as a template. All initial models were docked into the respective cryo-EM maps using UCSF Chimera^70,71^, followed by flexible fitting using the “all-atom refine” function in COOT^72^. Subsequent model refinement was performed with phenix.real_space_refine, applying secondary structure restraints, combined with iterative manual adjustments in COOT^72^. The quality of the final models was assessed using MolProbity as implemented in Phenix^73,74^. Data collection, processing, and model refinement statistics are summarized in **Supplemental Table 1**. All structural figures were prepared with UCSF Chimera^70^ and ChimeraX^69^. Chemokine residues are numbered from the first residue following signal sequence cleavage site.

### Lipidomics analysis

The lipidomics analysis was performed based on the method described previously^75^. Briefly, the purified CXCR7 sample (40 µL) and *Sf*9 cell lysate with overexpressed CXCR7 (40 µL) were mixed with 300 µL methanol and 1mL MTBE and incubated for 1 h on a roller at room temperature. 250 µl of HPLC H2O was added. The sample was vortex for 2-5 min to induce phase separation and centrifuged at 400g, at room temperature for 10 min. The supernatant was transferred to a new tube and further dried by a SpeedVac vacuum concentrator (Thermo Fisher Scientific). The lipids mixture was dissolved in 80% lipidomics mobile phase A/20% lipidomics mobile phase B (100 µL) and sonicated for 10 min. Mobile phase A consists of acetonitrile/H2O (60/40, vol/vol) supplemented with 10 mM ammonium formate and 0.1% (vol/vol) formic acid. Mobile phase B contains isopropanol/acetonitrile (90/10, vol/vol) supplemented with 10 mM ammonium formate and 0.1% (vol/vol) formic acid. For LC-MS/MS analysis, lipids were analyzed using a Dionex UltiMate 3000 RSLC nano system (Thermo Scientific) coupled to an Eclipse Tribrid Orbitrap mass spectrometer (Thermo Scientific). The samples were loaded onto a C18 column (Acclaim PepMap 100, C18, 75 µm × 150 mm, 3 µm particle size). A binary solvent system was applied. Lipid separation was performed at 40 °C using a gradient from 30% to 99% buffer B over 30 minutes, at a flow rate of 300 nL/min. The electrospray voltage was set to 2.2 kV, with a funnel RF level of 40 and a heated capillary temperature of 320 °C. For data-dependent acquisition (DDA), the full MS scan range was set from m/z 300 to 2,000, with a resolution of 120,000 and an AGC target of 100%. Fragmentation spectra were collected in the Orbitrap at a resolution of 15,000 using higher-energy collisional dissociation (HCD) with stepped collision energies of 25%, 30%, and 35%. Phospholipids were detected in the negative ion mode. The raw data were processed using MSDIAL for phospholipid identification and quantification^76^.

### Peptide library screening on CXCR7 using NanoBRET

Peptide library screening on ACKR3 was monitored by NanoBRET on living cells. HEK293T cells, stably expressing ACKR3, N-terminally fused to Nanoluciferase were distributed into white 384-well plates (1.5 × 10^4^ cells per well). Peptides from the Chop-Suey/Variety Peptide Library (L-001, 238 peptides), Bioactive Secretory Peptide Library (L-009, 1016 peptides) and Orphan Receptor Peptide Ligand Library (L-005, 150 peptides) from Phoenix Peptides were added to the cells at a final concentration of 1 µM and incubated for 5 min on ice. Cells were subsequently incubated with CXCL12-AZ568 (2 nM) and incubated for 2 h on ice. NanoGlo Live Cell Assay System substrate was then added and donor emission (450/8 nm BP filter) and acceptor emission (600 nm LP filter) were immediately measured on a GloMax Discover plate reader (Promega). BRET binding signal was defined as acceptor/donor ratio, and cells not treated with CXCL12-AZ568 were used to define 0% BRET binding, whereas cells that were treated with CXCL12-AZ568 alone were used to define 100% BRET binding.

