## Supplemental Figures for "Atypical activation and molecular glue-like dimerization mechanism of an intrinsically-biased chemokine receptor"

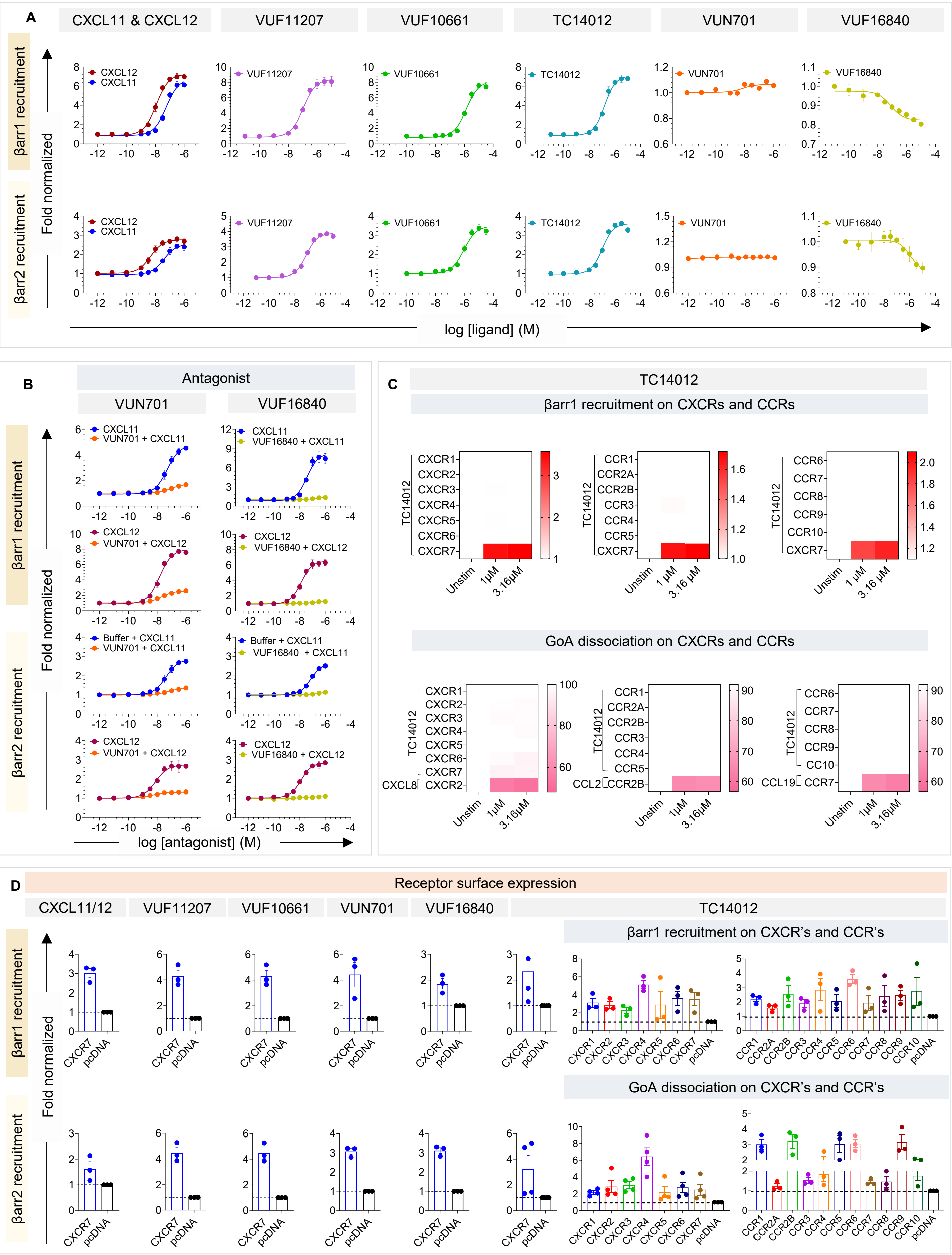
A
CXCL11 & CXCL12
VUF11207
VUF10661
TC14012
VUN701
VUF16840
βarr1 recruitment
βarr2 recruitment
Fold normalized
CXCL12
CXCL11
log [ligand] (M)
B
Antagonist
VUN701
VUF16840
VUN701 + CXCL11
VUF16840 + CXCL11
VUN701 + CXCL12
VUF16840 + CXCL12
Buffer + CXCL11
log [antagonist] (M)
C
TC14012
βarr1 recruitment on CXCRs and CCRs
GoA dissociation on CXCRs and CCRs
CXCR1
CXCR2
CXCR3
CXCR4
CXCR5
CXCR6
CXCR7
CCR1
CCR2A
CCR2B
CCR3
CCR4
CCR5
CCR6
CCR7
CCR8
CCR9
CCR10
CXCL8
CCL2
CCL19
Unstim
1μM
3.16μM
D
Receptor surface expression
CXCL11/12
βarr1 recruitment on CXCR's and CCR's
GoA dissociation on CXCR's and CCR's
CXCR7
pcDNA

**Figure S1. Pharmacological characterization of CXCR7 ligands. (A)** Ligand-induced βarr1/2 recruitment as measured using a NanoBiT assay (mean±sem; n=3; fold normalized with the response at lowest agonist concentration). **(B)** Antagonism of VUN701 and VUF16840 on CXCR7 as assessed using their ability to block CXCL11/CXCL12-induced βarr1/2 recruitment (mean±sem; n=3). **(C)** Selectivity of TC14012 at CCRs and CXCRs as measured using NanoBiT-based G-protein dissociation and βarr1/2 recruitment, respectively (mean; n=3; normalized with respect to unstimulated condition). **(D)** Surface expression of CXCR7 in the βarr1/2 recruitment assay (presented in panel A-C) as measured using whole cell surface ELISA (mean±sem; n=3; fold normalized with mock-transfected condition).

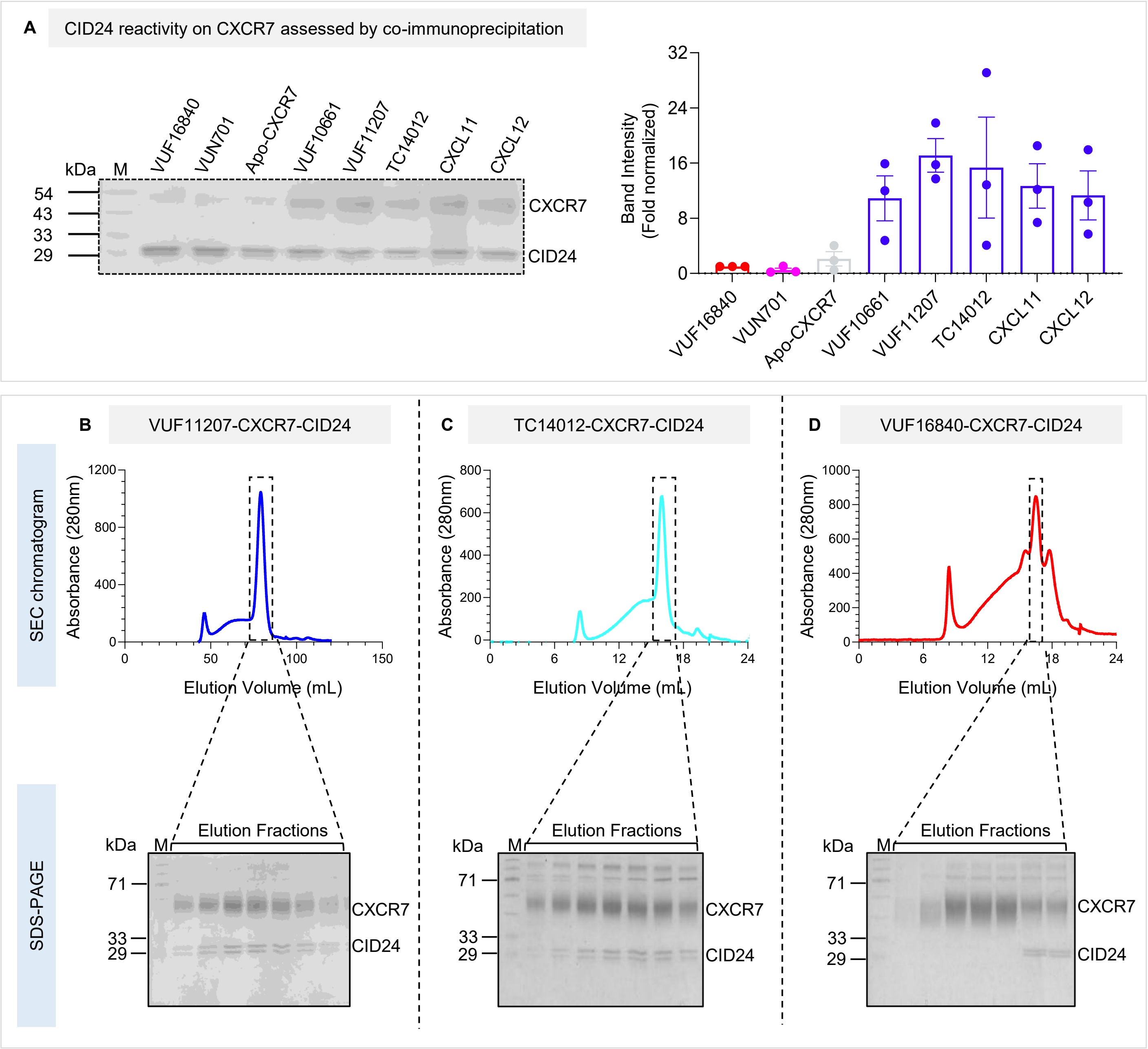


**Figure S2. Reconstitution of agonist-CXCR7-CID24 complexes for structural analysis. (A)** CID24 selectively recognizes agonist-bound conformation of CXCR7 as assessed using a co-immunoprecipitation assay with purified proteins. A representative gel image and densitometry-based quantification (mean±sem; n=3; % normalized with CXCL12 as 100). **(B-D)** Size exclusion chromatography (SEC) profiles and SDS-PAGE analysis of VUF11207-CXCR7-CID24 and TC14012-CXCR7-CID24 complexes. The other CXCR7 complexes also exhibited a similar SEC profile.

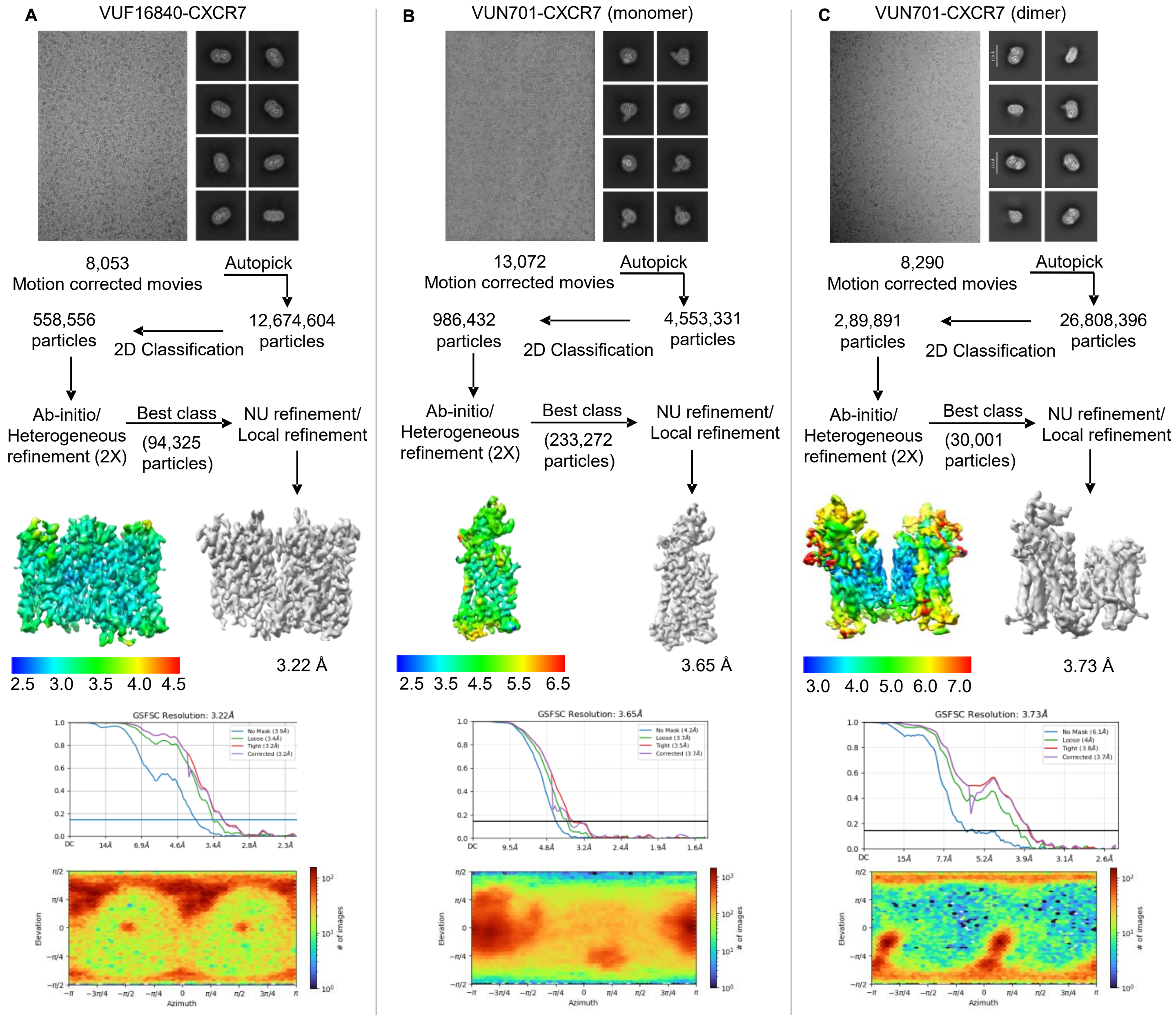


**Figure S3: Workflow of cryo-EM data processing for CXCR7 complexes. (A-C)** Representative cryo-EM micrographs, selected 2D class averages indicating different orientations, step-wise pipelines for data processing, local resolution maps of the 3D reconstructions, gold standard Fourier shell correlation curves at a threshold of 0.143, and angular distribution plots of the particles against the final reconstruction of the indicated samples.

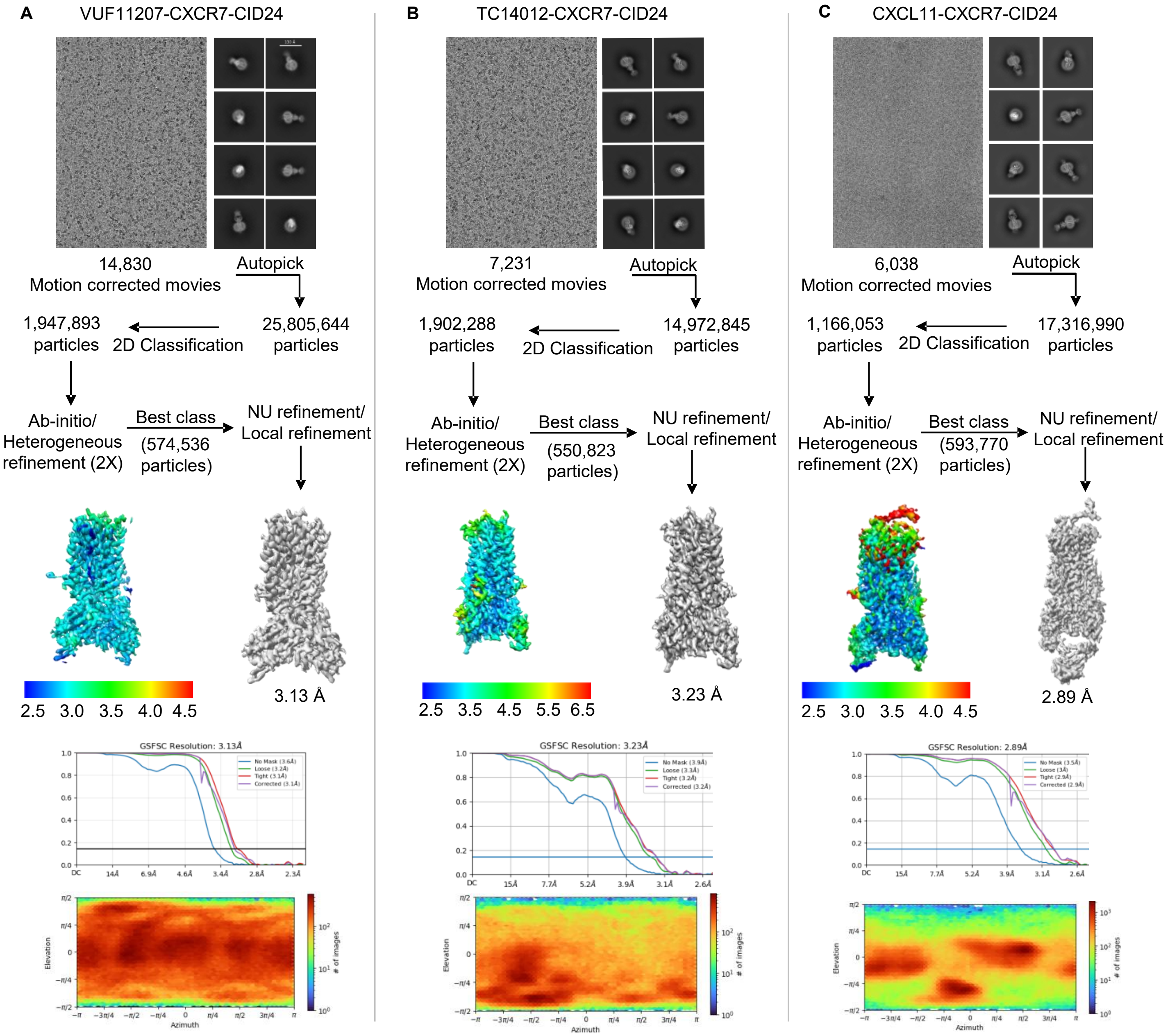


**Figure S4: Workflow of cryo-EM data processing for CXCR7 complexes. (A-C)** Representative cryo-EM micrographs, selected 2D class averages indicating different orientations, step-wise pipelines for data processing, local resolution maps of the 3D reconstructions, gold standard Fourier shell correlation curves at a threshold of 0.143, and angular distribution plots of the particles against the final reconstruction of the indicated samples.

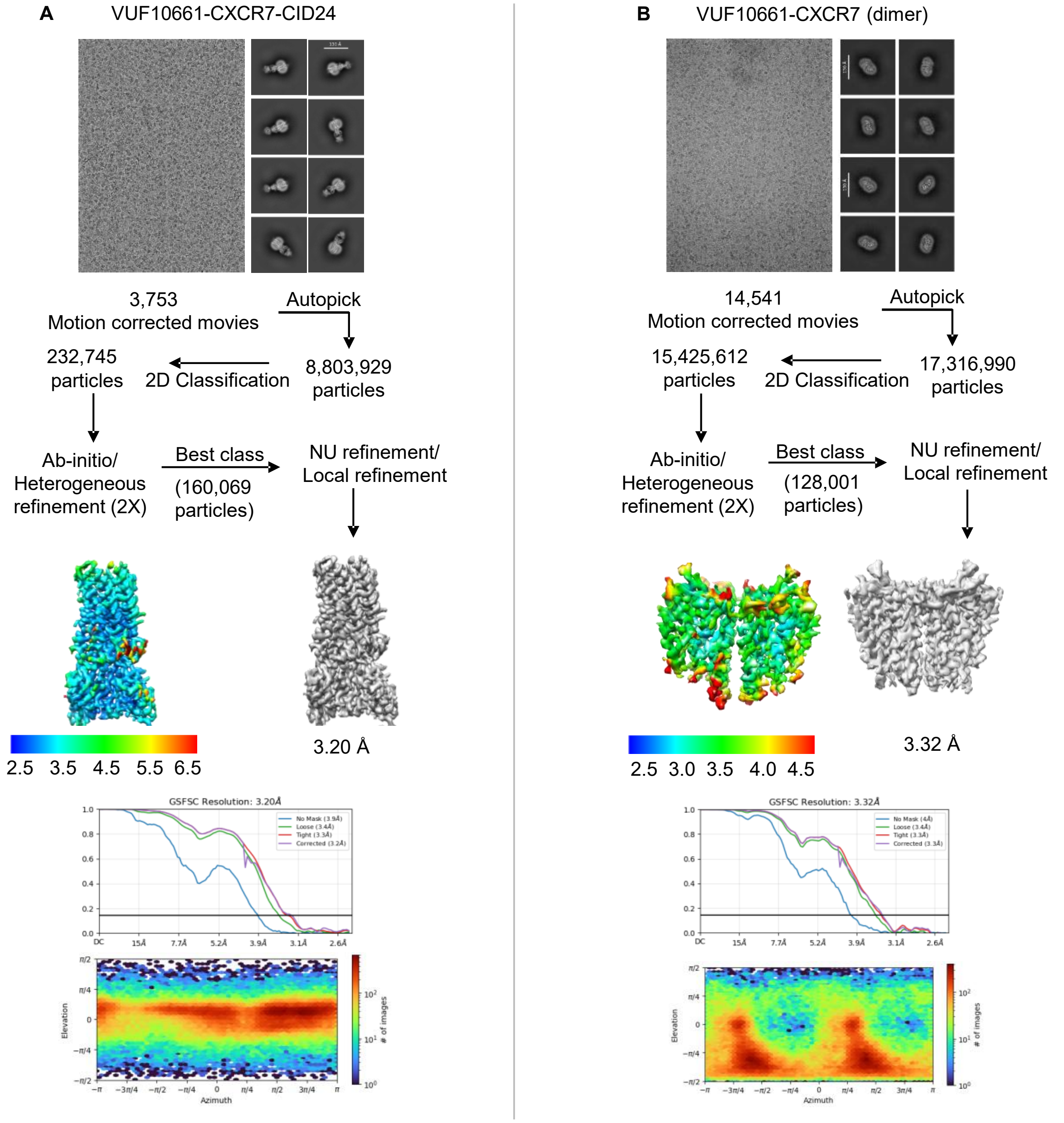


**Figure S5: Workflow of cryo-EM data processing for CXCR7 complexes. (A-B)** Representative cryo-EM micrographs, selected 2D class averages indicating different orientations, step-wise pipelines for data processing, local resolution maps of the 3D reconstructions, gold standard Fourier shell correlation curves at a threshold of 0.143, and angular distribution plots of the particles against the final reconstruction of the indicated samples.

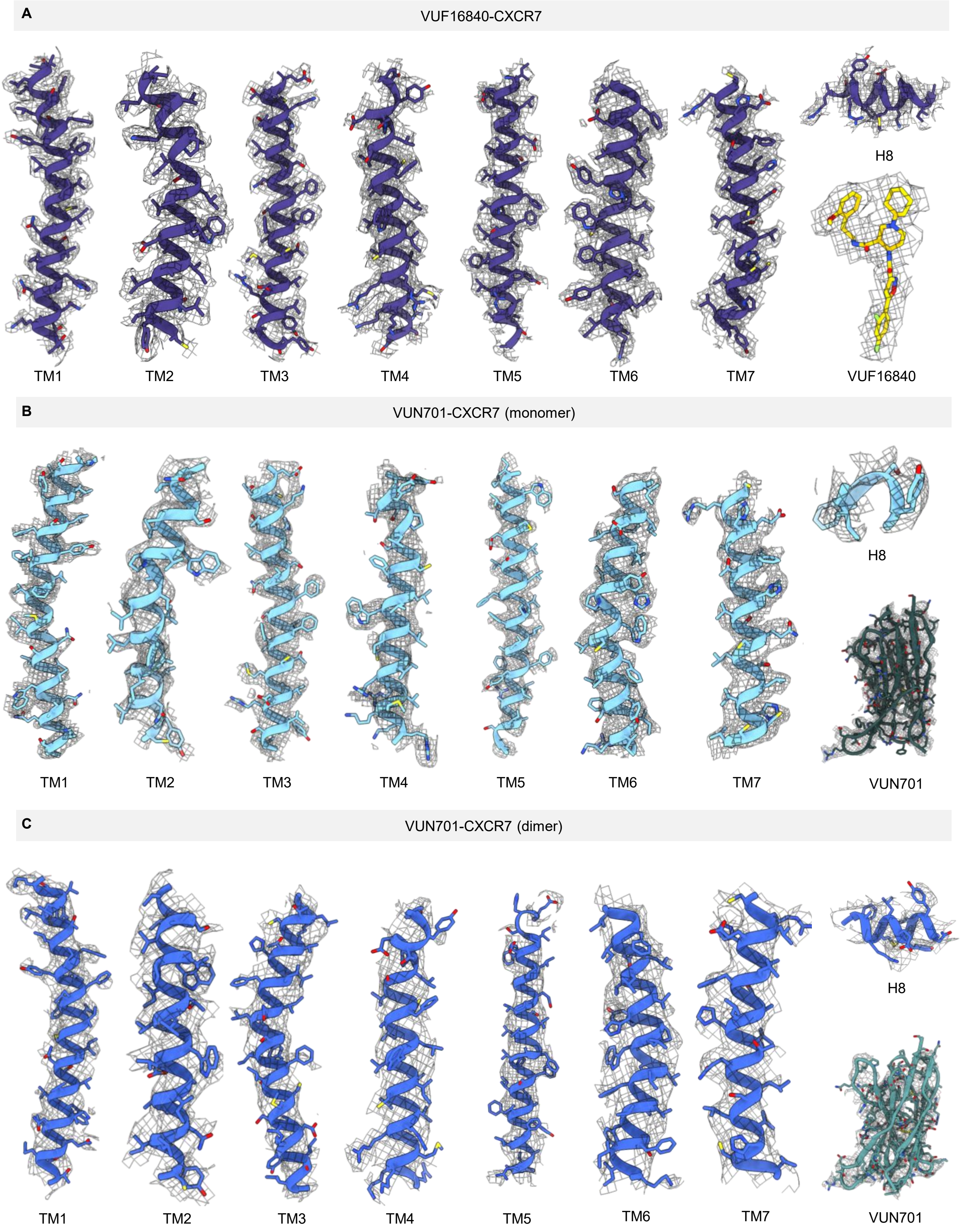


**Figure S6. cryo-EM densities of the TM segments and ligands in CXCR7 structures. (A-C)** cryo-EM densities of TM1-7, helix8, and the ligands are presented for the indicated structures.

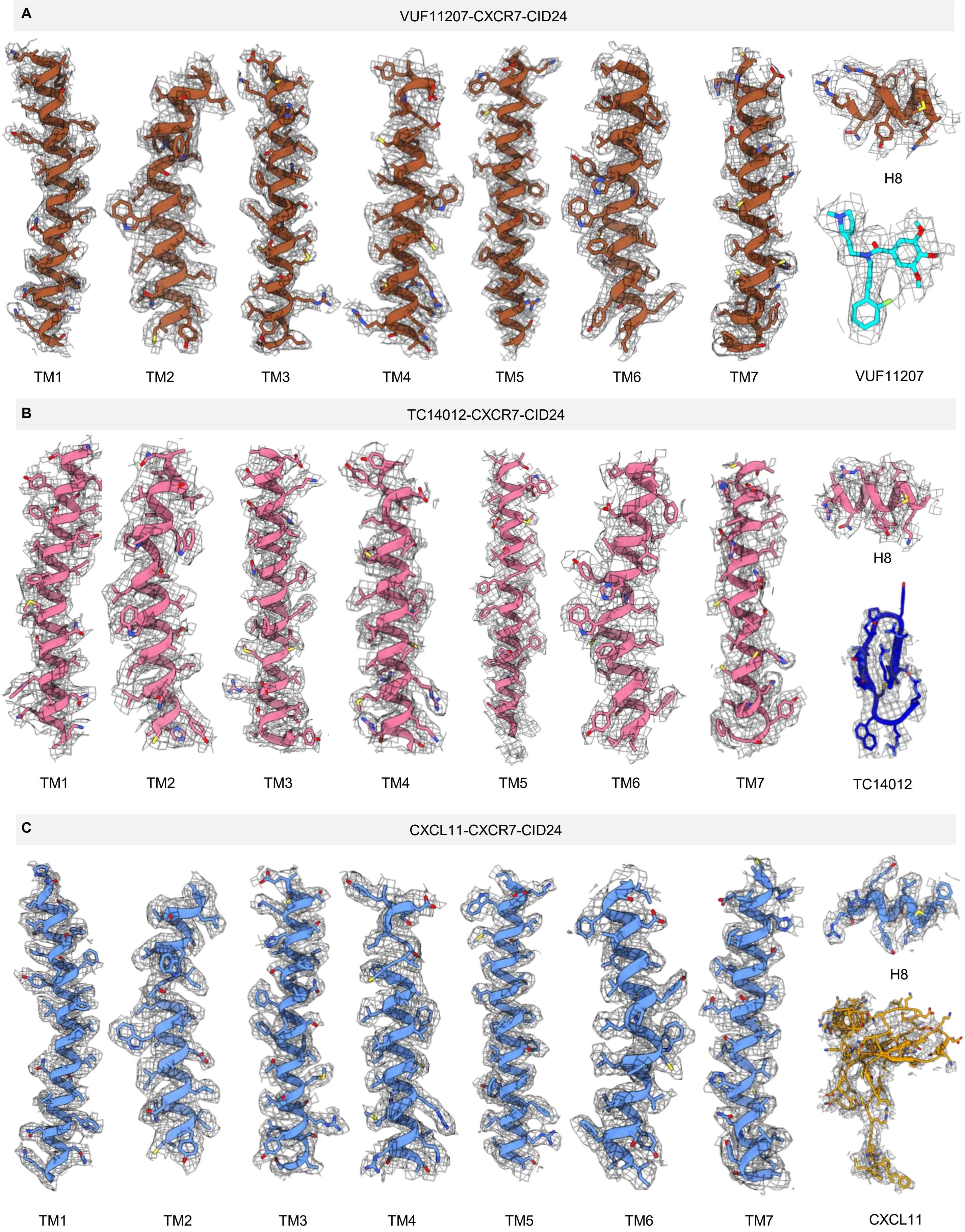


**Figure S7. cryo-EM densities of the TM segments and ligands in CXCR7 structures. (A-C)** cryo-EM densities of TM1-7, helix8, and the ligands are presented for the indicated structures.

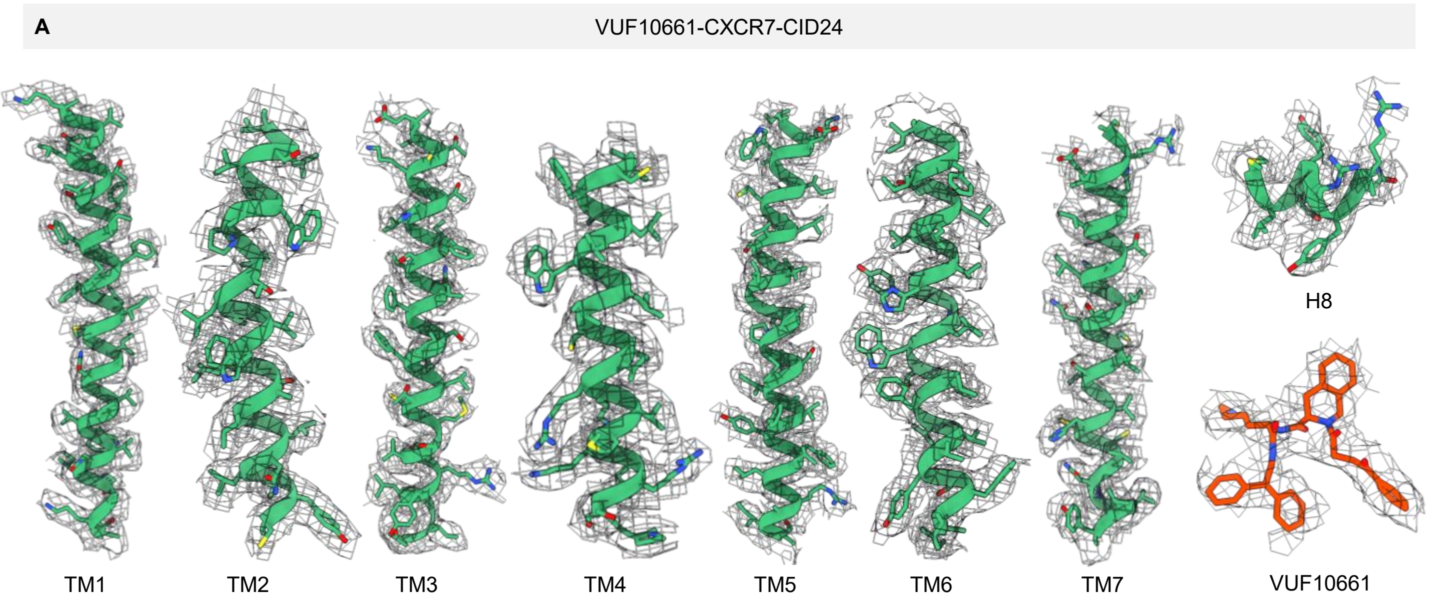


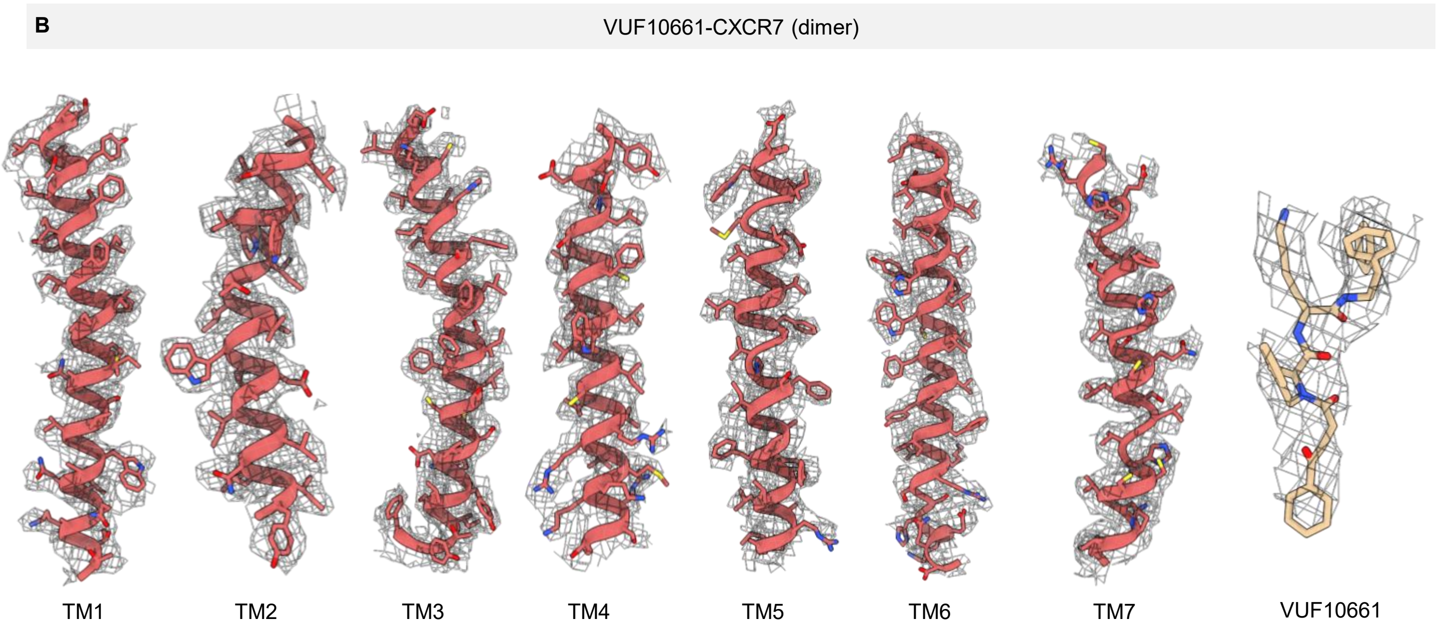


**Figure S8. cryo-EM densities of the TM segments and ligands in CXCR7 structures. (A-C)** cryo-EM densities of TM1-7, helix8, and the ligands are presented for the indicated structures.

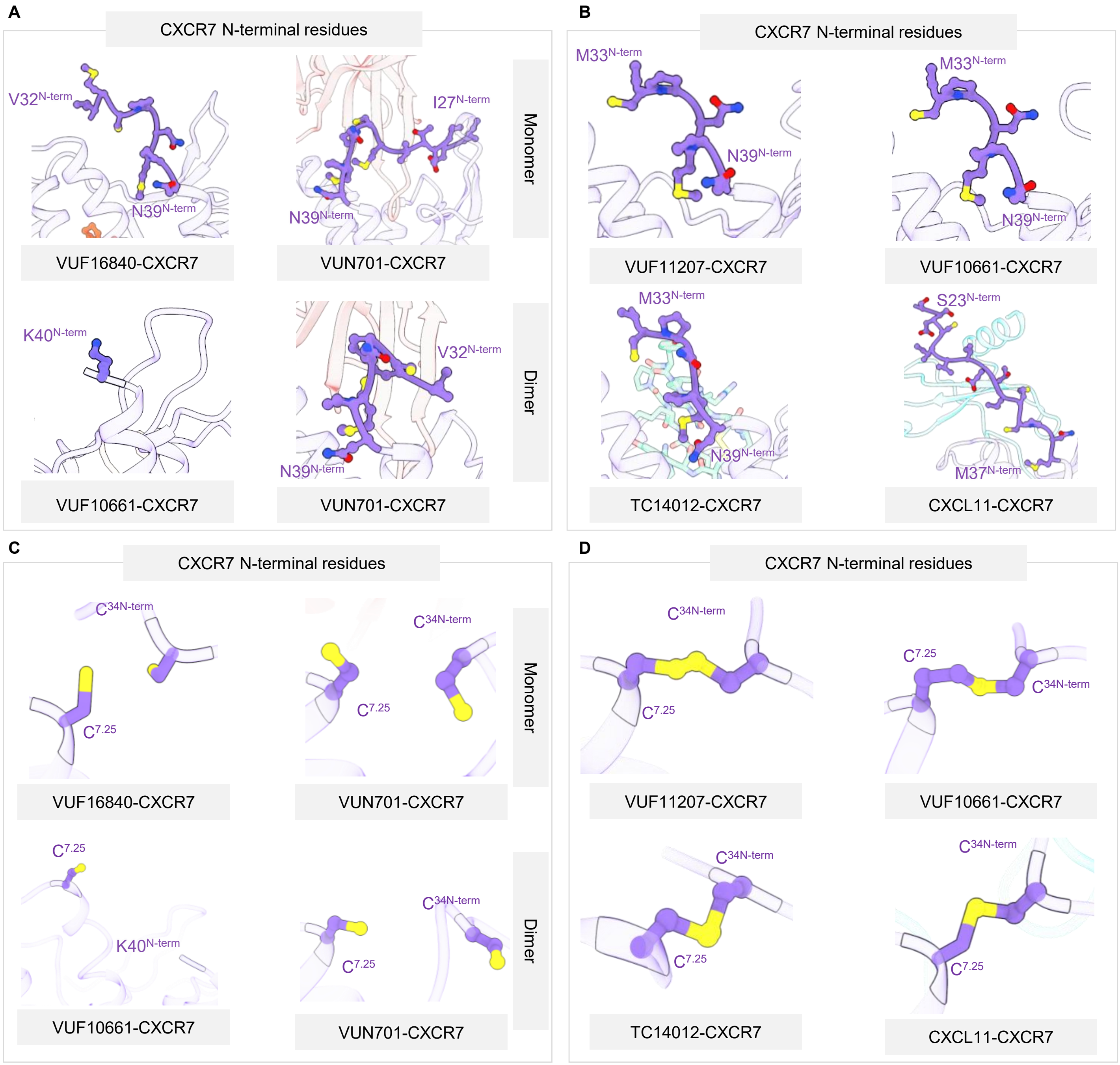


**Figure S9. The N-terminus and disulfide bridges in CXCR7 structures. (A-B)** Ribbon diagram of CXCR7 structures indicating the resolved segments of the N-terminus and their orientations. **(C-D)** The status of $Cys^{34}$-$Cys^{287}$ disulfide bridge in CXCR7 as resolved in the cryo-EM structures presented here.

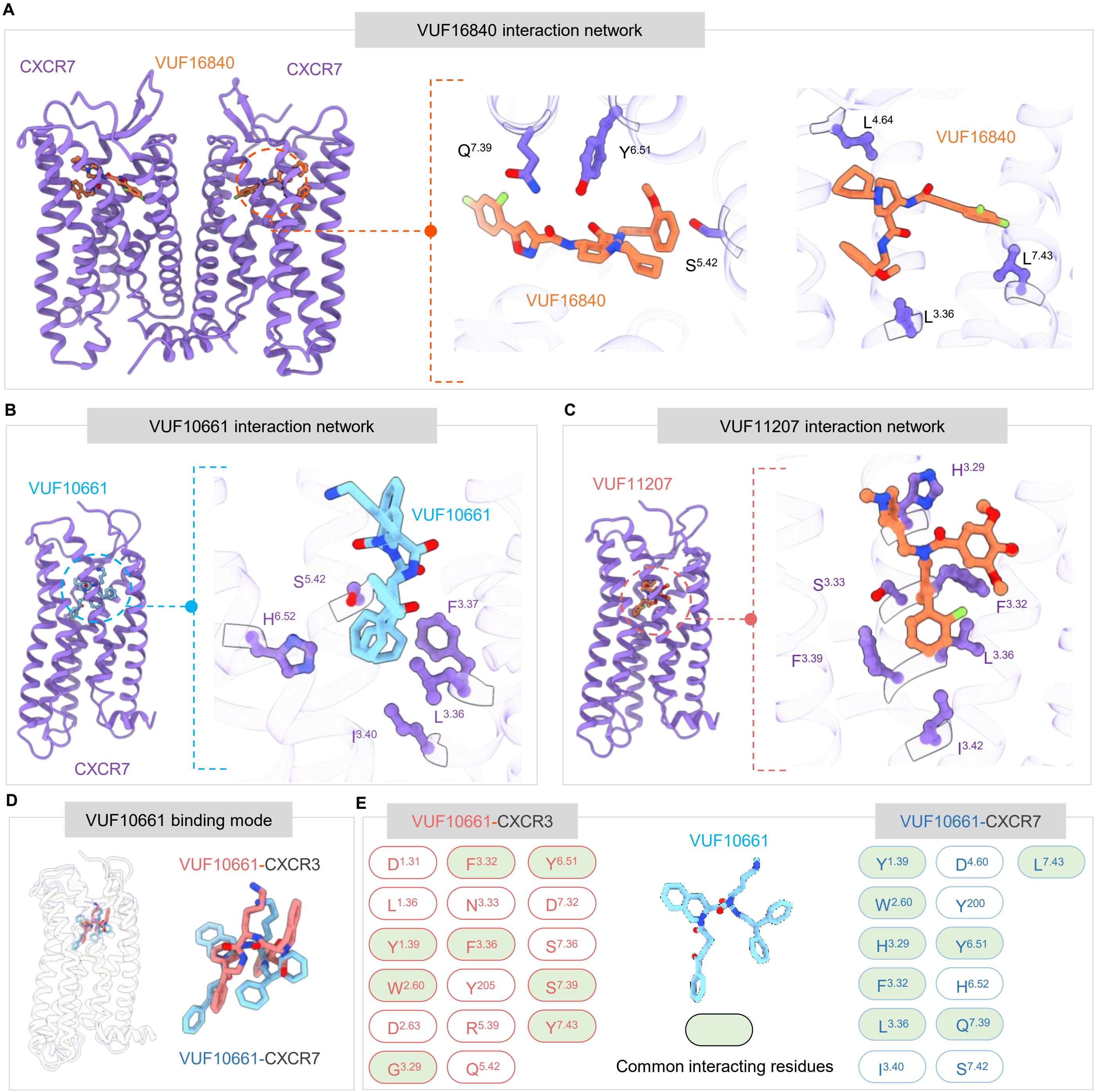


**Figure S10. Overall interaction of small molecule agonists with CXCR7 and comparison with CXCR3. (A-C)** The interaction network and hydrophobic binding pockets for VUF16840, VUF10661 (monomer), and VUF11207 in CXCR7 based on the cryo-EM structures presented here (PDB ID: 9WLG for VUF16840-CXCR7; 8WLF for VUF10661-CXCR7; 9WLE for VUF11207-CXCR7). **(D)** Structural superimposition of VUF10661-bound CXCR7 and CXCR3 depicting the binding of VUF10661 in the orthosteric pocket of the receptors (PDB ID: 8XYI). **(E)** Key interaction of VUF10661 with CXCR7 and CXCR3 are presented based on PDBSum. The common (analogous) interactions are indicated in blue, and the unique interactions are in red.

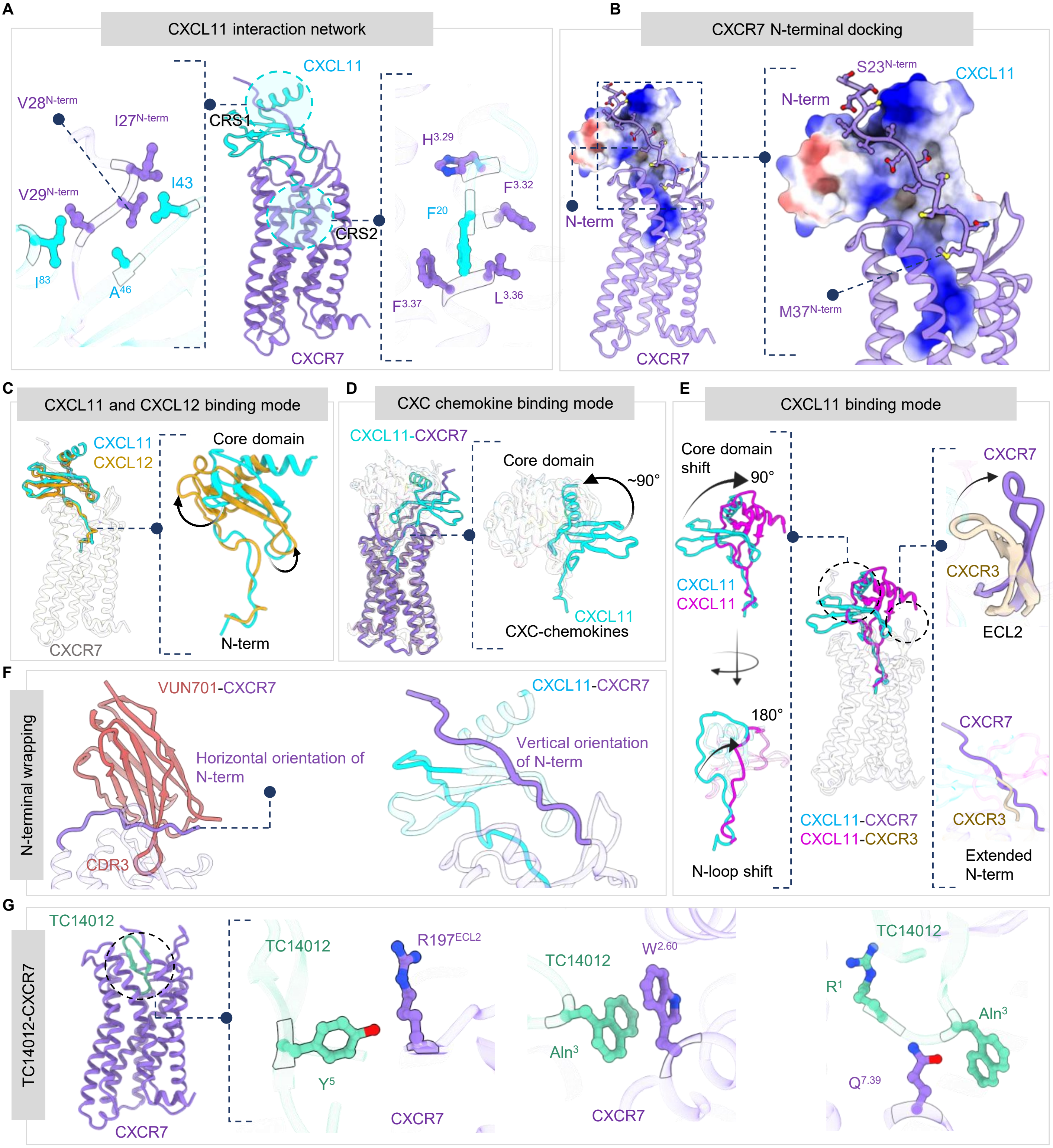
A
CXCL11 interaction network
CXCL11
CRS1
CRS2
CXCR7
V28$^{N-term}$
I27$^{N-term}$
V29$^{N-term}$
I43
I$^{83}$
A$^{46}$
H$^{3.29}$
F$^{3.32}$
F$^{20}$
F$^{3.37}$
L$^{3.36}$
B
CXCR7 N-terminal docking
N-term
CXCR7
S23$^{N-term}$
CXCL11
M37$^{N-term}$
C
CXCL11 and CXCL12 binding mode
CXCL11
CXCL12
Core domain
N-term
CXCR7
D
CXC chemokine binding mode
CXCL11-CXCR7
Core domain
~90°
CXCL11
CXC-chemokines
E
CXCL11 binding mode
Core domain shift
90°
CXCL11
CXCL11
180°
N-loop shift
CXCL11-CXCR7
CXCL11-CXCR3
CXCR7
CXCR3
ECL2
CXCR7
CXCR3
Extended N-term
F
N-terminal wrapping
VUN701-CXCR7
Horizontal orientation of N-term
CDR3
CXCL11-CXCR7
Vertical orientation of N-term
G
TC14012-CXCR7
TC14012
CXCR7
TC14012
R197$^{ECL2}$
Y$^{5}$
CXCR7
TC14012
W$^{2.60}$
Aln$^{3}$
CXCR7
TC14012
R$^{1}$
Aln$^{3}$
Q$^{7.39}$

**Figure S11. Overall interaction of CXCL11 with CXCR7 and comparative analysis. (A)** Ribbon diagram of CXCL11-CXCR7 structure depicting the two binding site mode of interaction. CRS1, Chemokine Recognition Site 1; CRS2, Chemokine Recognition Sequence 2 and key interactions involved in CXCL11-CXCR7 interaction based on the cryo-EM structure. **(B)** Interaction of the N-terminus of CXCR7 with the core domain of CXCL11 via a complementary charge interface as visualized in the CXCL11-CXCR7 structure. **(C)** A structural comparison of CXCL11 vs. CXCL12 binding to CXCR7 depicting a lateral displacement between their positioning on the receptor (PDB ID: 7SK5 for CXCL12-CXCR7). **(D)** A distinct binding pose of CXCL11 on CXCR7 compared to other C-X-C chemokines on CXCRs based on cryo-EM structures (PDB ID: 8IC0 for CXCR1; 8XWA, 8XWV, 8XVU, 8XWF, 8XWS, 8XWN, 8XWM, for CXCR2, for CXCR4 used for comparison). **(E)** Comparison of CXCL11-binding to CXCR7 and CXCR3 depicting distinct binding poses, and relative orientations of ECL2 between the two structures and Key differences in the binding of CXCL11 with CXCR7 vs. CXCR3 in terms of overall positioning and interactions. **(F)** Key structural differences in the interaction of VUN701 vs. CXCL11 with CXCR7 including CDR3 insertion in the orthosteric binding pocket and distinct conformation of CXCR7 N-terminus. **(G)** Overall positioning and key interactions of TC14012 in the orthosteric binding pocket.

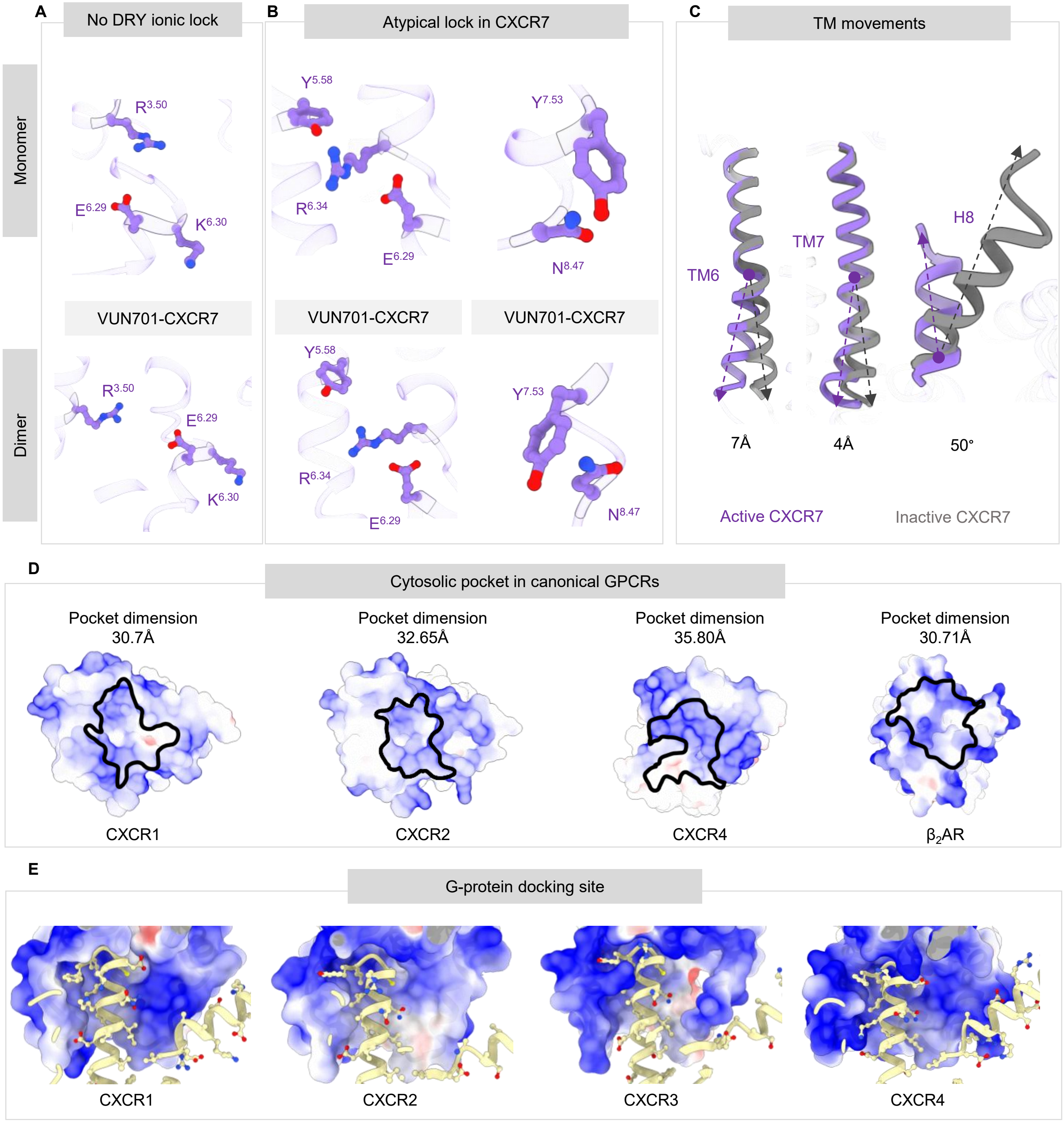


**Figure S12. Atypical ionic-locks and activation of CXCR7. (A)** Structural representation depicting the absence of TM3-TM6 lock in VUN701-bound CXCR7 (upper panel, monomer; lower panel, dimer). **(B)** Structural representation depicting the newly-identified TM5-TM6 tri-partite lock and TM7-H8 ionic lock in VUN701-CXCR7 (upper panel, monomer; lower panel, dimer). **(C)** Activation-dependent outwards movement of TM6, TM7 and helix8 in CXCR7 based on structural comparison of VUF1684-CXCR7 and CXCL11-CXCR7 structures. **(D-E)** Comparison of the cytoplasmic surface of active CXCR7 with selected prototypical GPCRs in terms of the dimension of the cytoplasmic cavity and the surface for the docking of α5 and αN helices of Gα subunit, respectively.

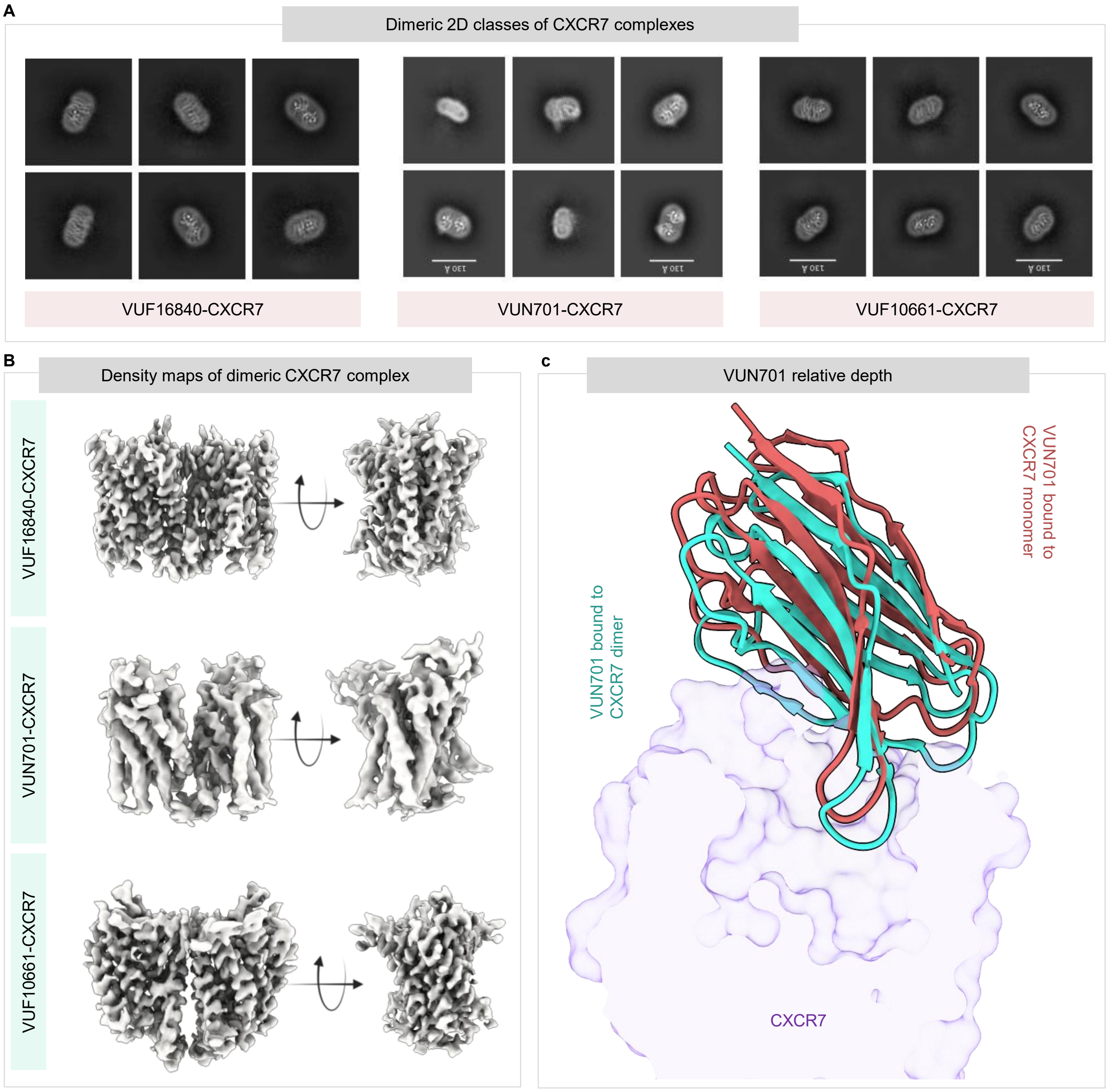


**Figure S13. Dimerization of CXCR7. (A)** 2D class averages of the indicated CXCR7 samples depicting dimeric arrangement of the receptor. **(B)** Overall, cryo-EM map of CXCR7 dimeric structures shown as the front and side views. **(C)** Overall positioning of VUN701 in monomeric and dimeric CXCR7 structures as visualized by structural superimposition (PDB ID: 9WLI and 9WLO for VUN701-CXCR7 monomer and VUN701-CXCR7 dimer, respectively).

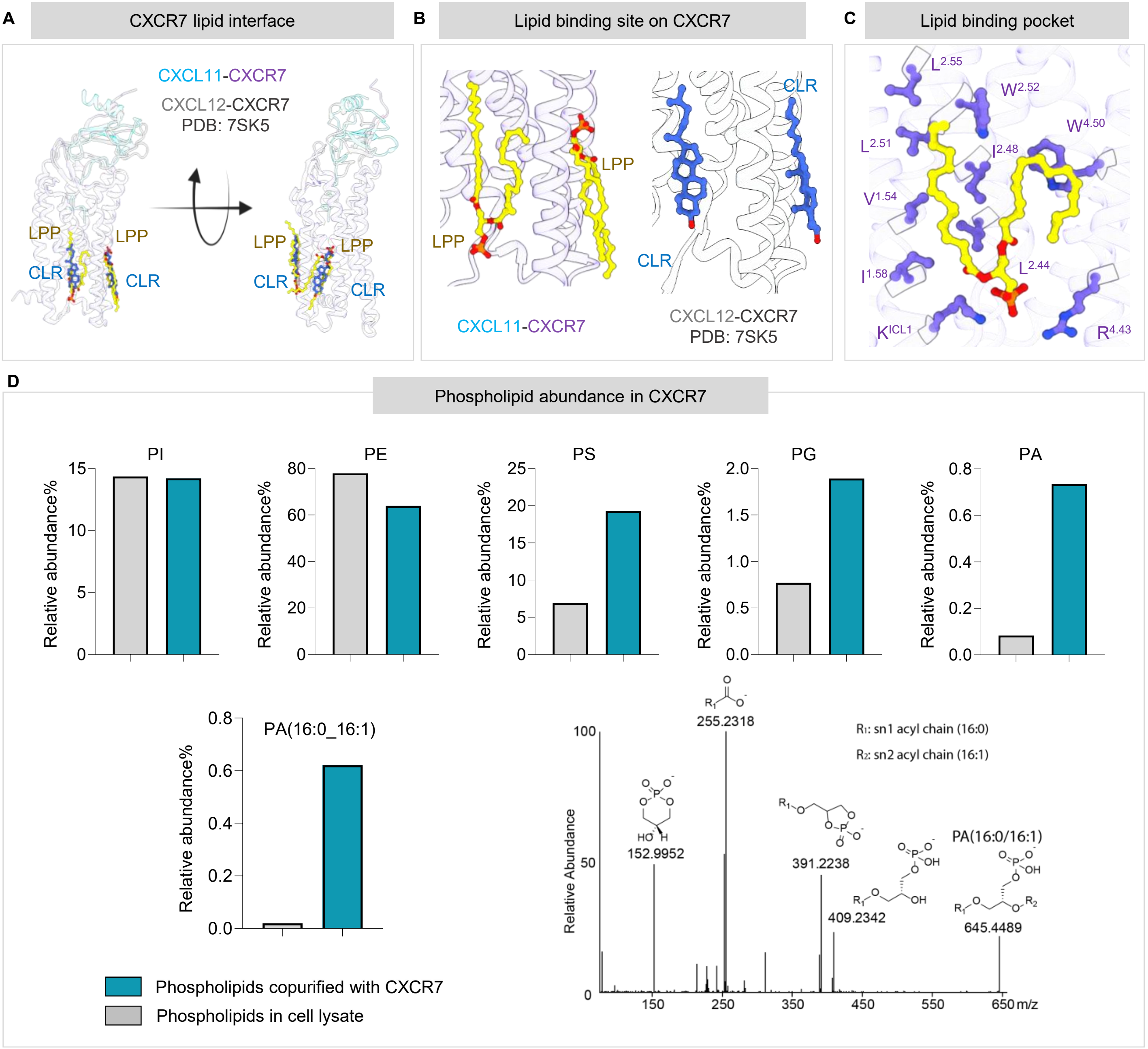


**Figure S14. Interaction of the native lipids with CXCR7. (A-B)** Comparison of LPP binding sites on the receptor in the CXCL11-CXCR7 structure with previously reported CXCL12-CXCR7 structure, where two cholesterol molecules were modeled in analogous EM densities. **(C)** Key interaction of the LPP lipid moieties with the CXCR7 residues based on the CXCL11-CXCR7 structure. **(D)** Lipidomics-based identification of the relative abundance of indicated phospholipids in *Sf*9 membrane and purified CXCR7 sample.

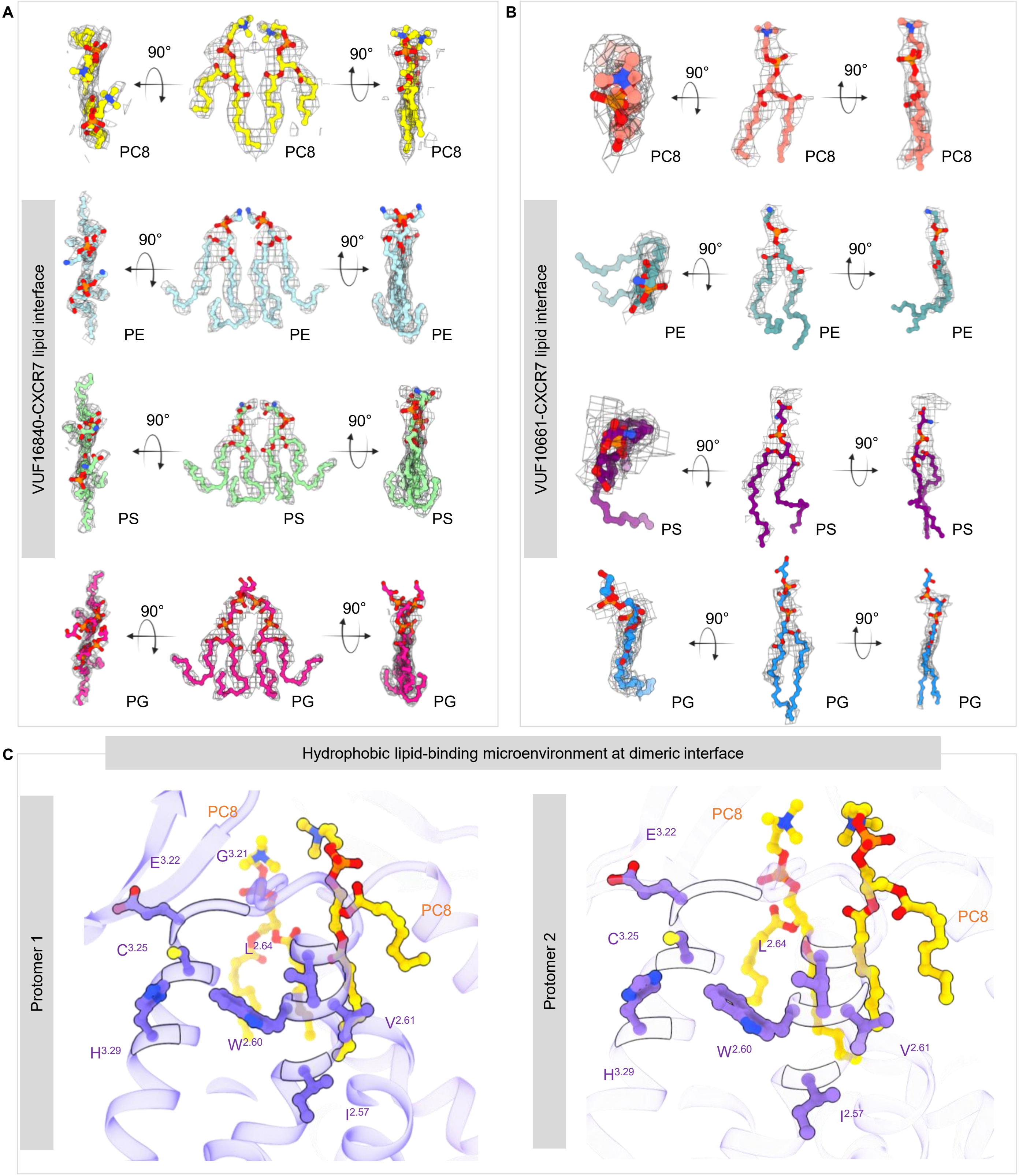


**Figure S15. Interaction of the native lipids with CXCR7. (A-B)** Structural modeling, guided by lipidomics analysis, of the indicated native lipids in the cryo-EM densities observed at the dimer interface in VUF16840 and VUF10661-bound CXCR7 structures. **(C)** Hydrophobic pocket accommodating phosphatidylcholine moieties at the dimer interfaces in VUF16840- and VUF10661-bound CXCR7 structures.

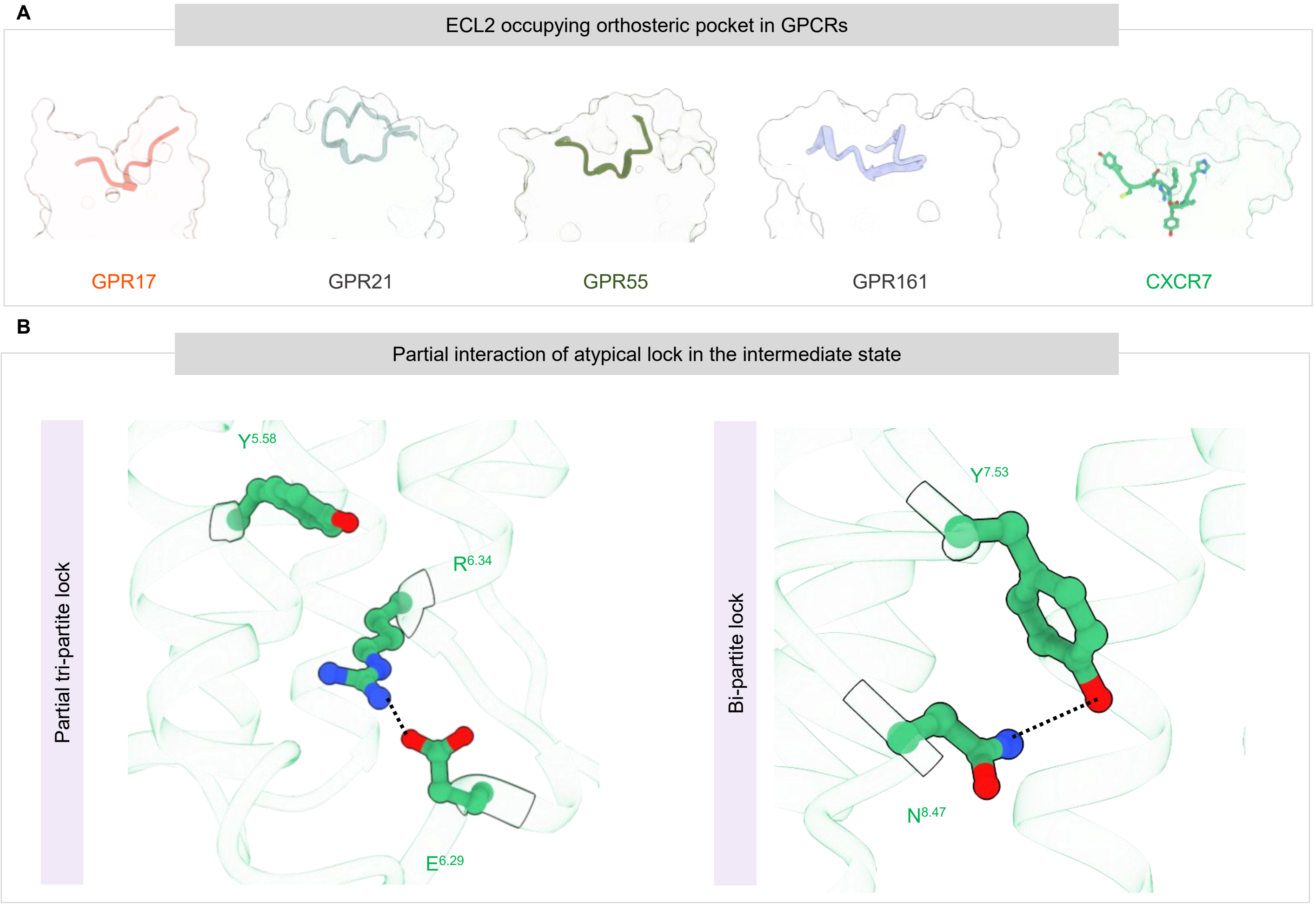


**Figure S16. A novel allosteric site and an intermediate conformation of CXCR7. (A)** A comparison of ECL2 orientation in VUF10661-CXCR7 dimeric structure that in other orphan GPCRs namely, GPR17, GPR21, GPR52, GPR55, and GPR161. **(B)** An intermediate conformation of the newly identified ionic-locks in CXCR7, wherein the Arg251$^{6.34}$-Glu246$^{6.29}$ and Tyr315$^{7.53}$-Asn319$^{8.47}$ interactions are maintained while the Arg251$^{6.34}$-Tyr232$^{5.58}$ lock is disrupted.

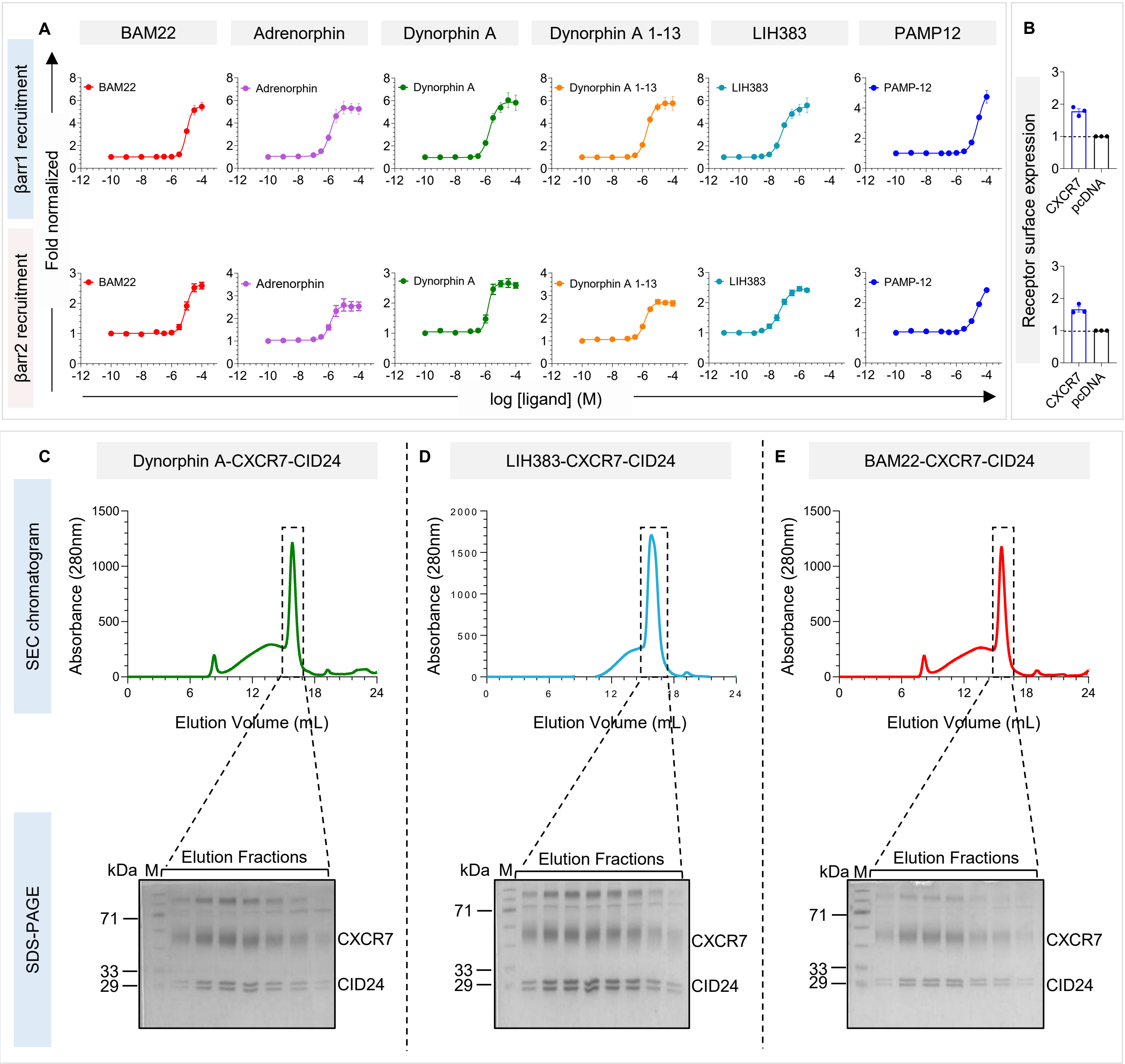


**Figure S17. Binding and activation of CXCR7 by opioid-peptides. (A)** Ligand-induced βarr1 and 2 recruitment as measured using a NanoBiT assay (mean±sem; n=3; fold normalized with the response at lowest agonist concentration). **(B)** Surface expression of CXCR7 as measured in the βarr1/2 recruitment assay (presented in panel A-B) (mean±sem; n=3; fold normalized with mock-transfected condition). **(C-E)** Reconstitution of agonist-CXCR7-CID24 complexes for structural analysis. Size exclusion chromatography (SEC) profiles and SDS-PAGE analysis of the indicated complexes are presented.

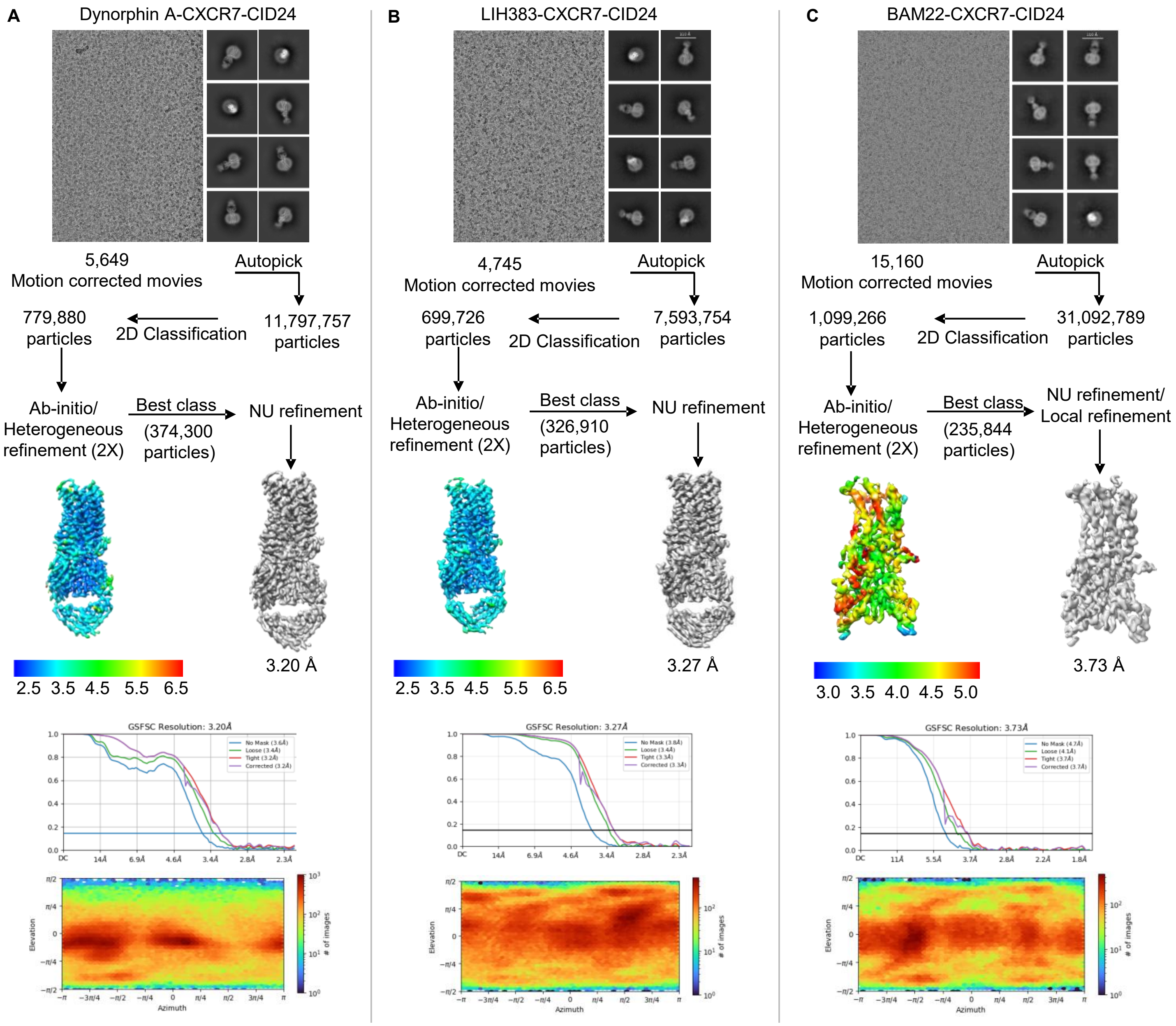


**Figure S18. Workflow of cryo-EM data processing for CXCR7 complexes. (A-C)** Representative cryo-EM micrographs, selected 2D class averages indicating different orientations, step-wise pipelines for data processing, local resolution maps of the 3D reconstructions, gold standard Fourier shell correlation curves at a threshold of 0.143, and angular distribution plots of the particles against the final reconstruction of the indicated samples.

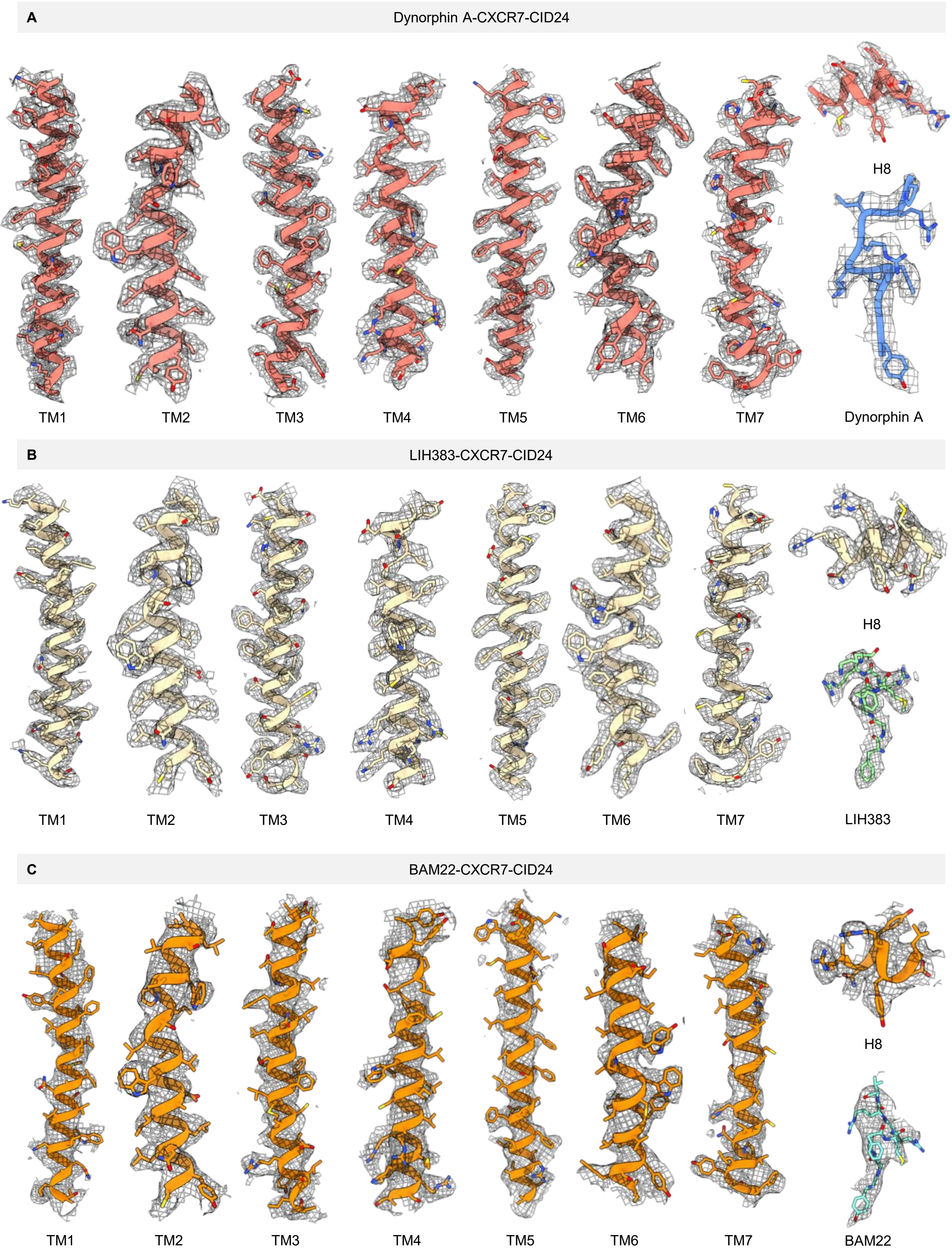


**Figure S19. cryo-EM densities of the TM segments and ligands in CXCR7 structures. (A-C)** cryo-EM densities of TM1-7, helix8, and the ligands are presented for the indicated structures.

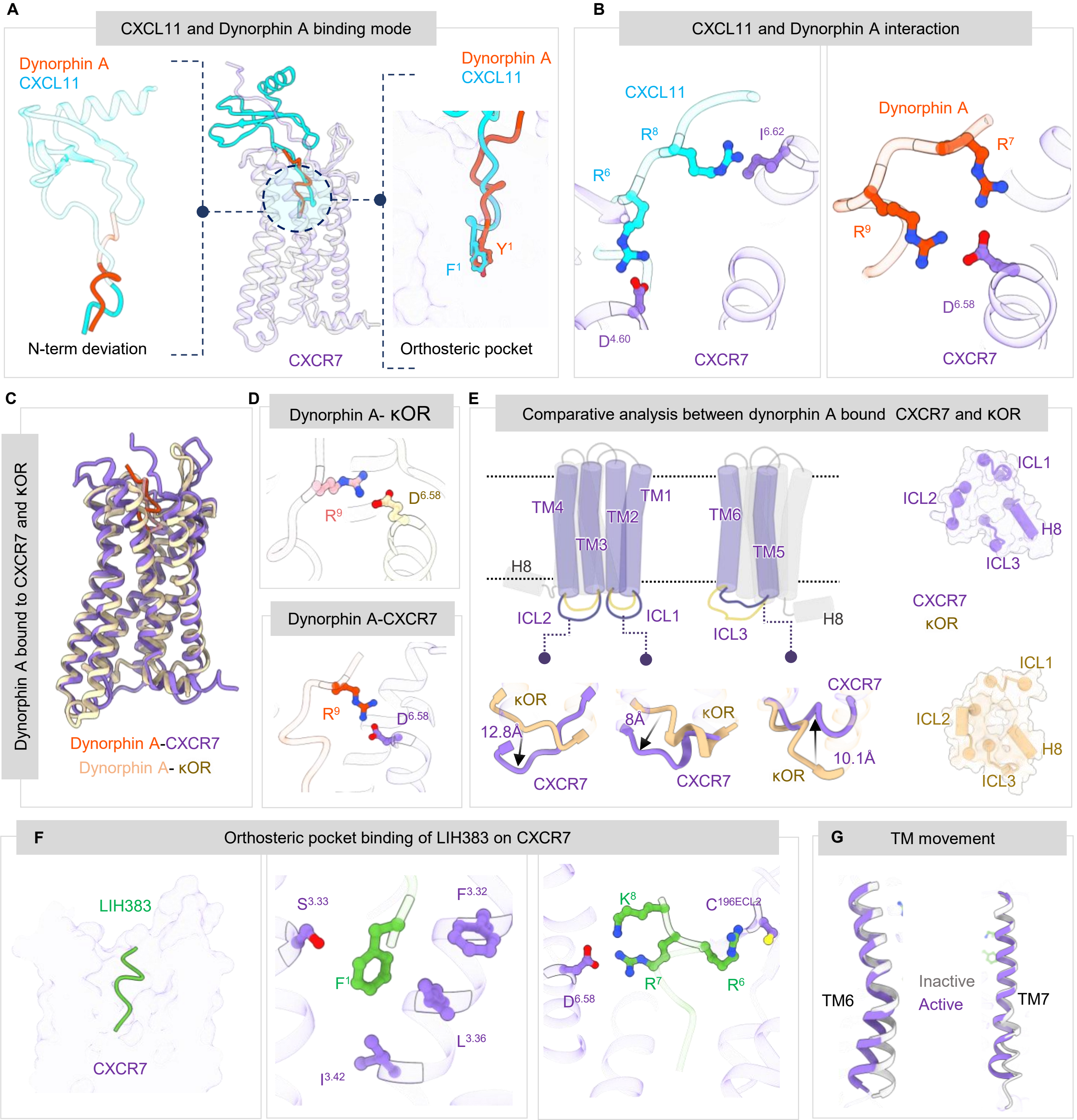


**Figure S20. Comparison of dynorphin-CXCR7 and CXCL11-CXCR7 structures and recognition of opioid peptides by CXCR7 and receptor activation. (A)** Structural superimposition of dynorphin-bound CXCR7 and CXCL11-bound CXCR7 structures (PDB ID: 9WLL for dynorphin-CXCR7; 9WLD for CXCL11-CXCR7) and overall positioning of the terminal residues i.e. Tyr$^{1}$ and Phe$^{1}$ of dynorphin and CXCL11, respectively, in the orthosteric binding pocket of CXCR7. **(B)** Structural comparison of overall positioning and interaction of dynorphin and the N-terminus of CXCL11 in the orthosteric binding pocket of CXCR7. **(C)** Structural superimposition of dynorphin-bound CXCR7 and kappa-opioid receptor (κ-OR) (PDB ID: 9WLL for dynorphin-CXCR7; 8F7W for dynorphin-κ-OR). **(D)** Structural comparison of dynorphin binding to CXCR7 and κ-OR indicating a convergent position of R9 in dynorphin on the two receptors. **(E)** Schematic illustration and ribbon representation to depict the key differences in the ICL1/2/3 and DRY motif between the dynorphin-bound CXCR7 and κ-OR followed by structural comparison of the cytoplasmic surface of the dynorphin-bound CXCR7 vs. κ-OR indicating a relatively constricted pocket in CXCR7. **(F)** Overall binding pose and key interactions of LIH383 with CXCR7 based on the cryo-EM structures (PDB ID: 9WLJ for LIH383-CXCR7). **(G)** The outward movement of TM7 and TM7 in BAM22/LIH383-bound CXCR7 compared to VUF16840-CXCR7.

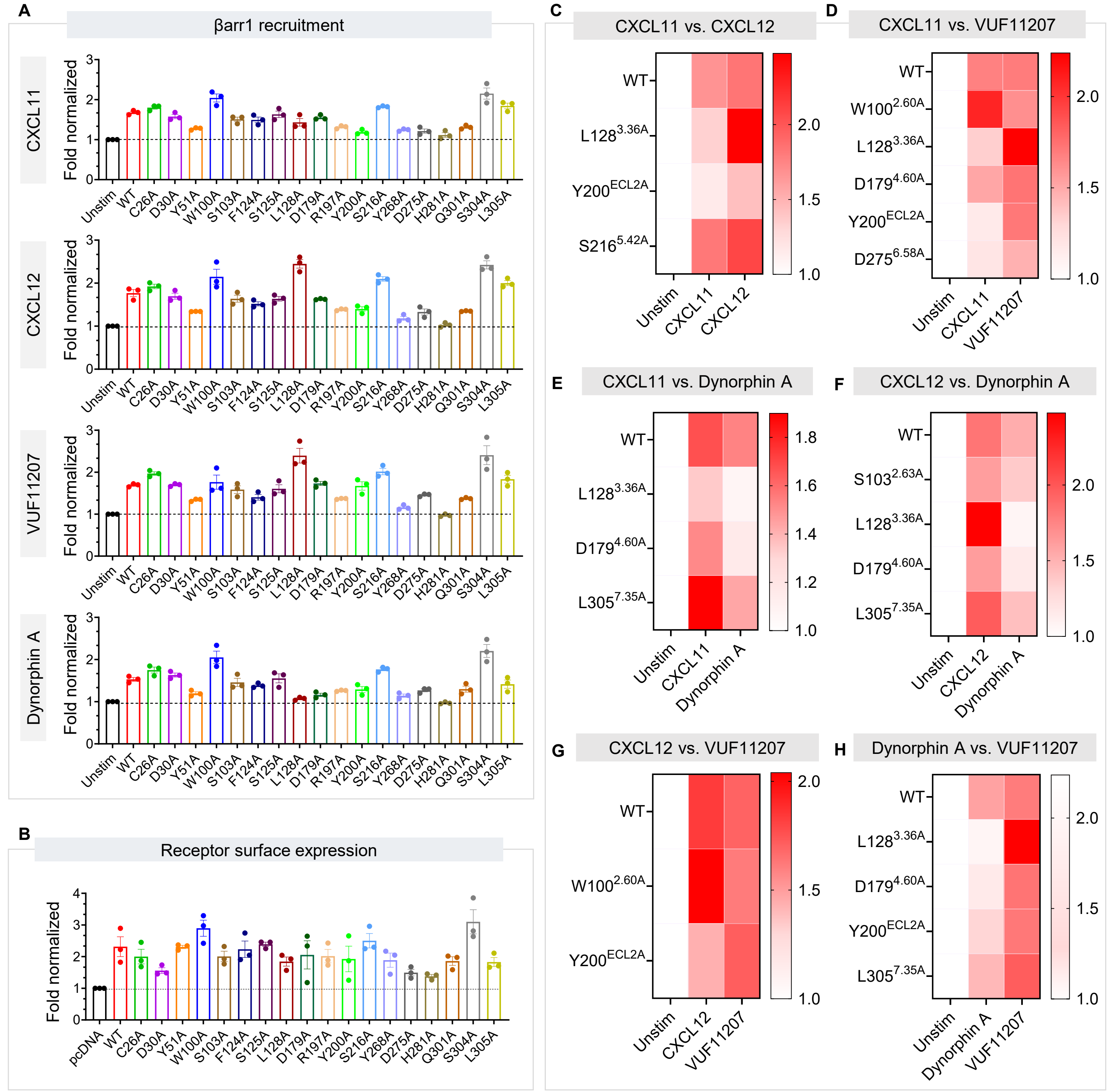


**Figure S21. Agonist-induced βarr recruitment at CXCR7 mutants. (A)** Ligand-induced βarr1 recruitment for the CXCR7 mutants as measured using a NanoBiT assay in transfected HEK-293T cells (mean±sem; n=3; fold normalized with the response at lowest agonist concentration). **(B)** Surface expression of CXCR7 mutants in the βarr1 recruitment assay (presented in panel A) as measured using whole cell surface ELISA (mean±sem; n=3; fold normalized with mock-transfected condition). **(C-H)** Heatmap representation of βarr1 recruitment response for selected mutants and ligands as indicated. The data points are taken from panel A and replotted (mean; n=3; fold-normalized with unstimulated condition).
